# Coordination of cytochrome *bc*_1_ complex assembly at MICOS

**DOI:** 10.1101/2024.02.22.580370

**Authors:** Ralf M. Zerbes, Lilia Colina-Tenorio, Maria Bohnert, Christian D. Peikert, Karina von der Malsburg, Carola S. Mehnert, Inge Perschil, Rhena Klar, Ida van der Klei, Silke Oeljeklaus, Bettina Warscheid, Heike Rampelt, Martin van der Laan

**Author notes:** Corresponding authors. (M.v.d.L.); (H.R.).

## Abstract

The boundary and cristae domains of the mitochondrial inner membrane are connected by crista junctions. Most cristae membrane proteins are nuclear-encoded and inserted by the mitochondrial protein import machinery into the inner boundary membrane. Thus, they must overcome the diffusion barrier imposed by crista junctions to reach their final location. Here, we show that respiratory chain complexes and assembly intermediates are physically connected to the mitochondrial contact site and cristae organizing system (MICOS) that is essential for formation and stability of crista junctions. We identify the inner membrane protein Mar26 (Fmp10) as determinant in the biogenesis of the cytochrome *bc*_1_ complex (complex III). Mar26 couples a Rieske Fe/S protein-containing assembly intermediate to MICOS. Our data indicate that Mar26 maintains an assembly-competent Rip1 pool at crista junctions where complex III maturation likely occurs. MICOS facilitates efficient Rip1 assembly by recruitment of complex III assembly intermediates to crista junctions. We propose that MICOS, via interaction with assembly factors such as Mar26, directly contributes to the spatial and temporal coordination of respiratory chain biogenesis.

## INTRODUCTION

Conversion of the energy contained in nutrients into the universal cellular energy currency adenosine triphosphate (ATP) is a fundamental metabolic program in all living organisms. In eukaryotic cells the majority of ATP is synthesized in mitochondria via a process termed oxidative phosphorylation (OXPHOS). The protein machineries that carry out the underlying chemical reactions - the respiratory chain complexes and the F_1_F_o_- ATP synthase - are embedded into the mitochondrial inner membrane that is composed of two subcompartments with distinct functions and protein compositions. The inner boundary membrane is in close proximity to the outer membrane and is enriched in transporters, like the components of the mitochondrial protein import machinery (Zick *et al*, 2009; Horvath *et al*, 2015). The oxidative phosphorylation system of mitochondria is mainly localized to the cristae of the inner membrane that shape a specialized microcompartment for chemi-osmotic coupling (Gilkerson *et al*, 2003; Mannella, 2006; Vogel *et al*, 2006; Wurm & Jakobs, 2006; Zick *et al*, 2009; Davies *et al*, 2011; Appelhans *et al*, 2012; Wilkens *et al*, 2013; Cogliati *et al*, 2016; van der Laan *et al*, 2016; Kondadi *et al*, 2020; Colina Tenorio *et al*, 2020; Kondadi & Reichert, 2024). Cristae membranes are highly folded invaginations protruding from the inner boundary membrane into the central matrix compartment. They are connected to the boundary membrane via narrow tubular openings, termed crista junctions, that are thought to act as a diffusion barrier (Perkins *et al*, 1997; Frey *et al*, 2002; Mannella, 2006; Zick *et al*, 2009; Wollweber *et al*, 2017; Wolf *et al*, 2019; Colina Tenorio *et al*, 2020). The asymmetric protein distribution between inner boundary and cristae membranes imposes a logistical challenge for the mitochondrial protein sorting system. Many of the membrane-integral subunits of the OXPHOS machinery are encoded by nuclear genes and synthesized in the cytosol as cleavable precursor proteins with an amino-terminal presequence. These proteins are recognized by dedicated mitochondrial surface receptors and enter the organelle via the general protein translocase of the outer membrane (TOM complex) (Harbauer *et al*, 2014; Araiso *et al*, 2022; Busch *et al*, 2023). Insertion into the inner boundary membrane is mediated by the presequence translocase of the inner mitochondrial membrane (TIM23 complex) and its partner protein complexes (Mokranjac & Neupert, 2010; Schulz *et al*, 2015; Moulin *et al*, 2019; Busch *et al*, 2023; Fielden *et al*, 2023; Zhou *et al*, 2023). Thus, somewhere on the way from membrane insertion of OXPHOS proteins at the inner boundary membrane to their final destination in the cristae membranes, the diffusion barrier imposed by the crista junctions must be crossed. In fact, early versus late assembly steps of complex III and IV are localized asymmetrically: While early steps preferentially take place in the inner boundary membrane, late ones are more prevalent in the cristae membranes (Stoldt *et al*, 2018).

Crista junctions with their high local membrane curvature require for their stability the mitochondrial contact site and cristae organizing system (MICOS) (Harner *et al*, 2011; Hoppins *et al*, 2011; Malsburg *et al*, 2011; Friedman & Nunnari, 2014; Kozjak-Pavlovic, 2017; Rampelt *et al*, 2017; Wollweber *et al*, 2017). The MICOS complex is highly conserved in evolution and consists of at least six different genuine subunits in yeast and seven in mammals that are organized in two distinct modules (Rabl *et al*, 2009; Harner *et al*, 2011; Hoppins *et al*, 2011; Malsburg *et al*, 2011; Alkhaja *et al*, 2012; Ott *et al*, 2012; Pfanner *et al*, 2014; Guarani *et al*, 2015; Muñoz-Gómez *et al*, 2015; Huynen *et al*, 2016; Colina Tenorio *et al*, 2020; Mukherjee *et al*, 2021; Bock-Bierbaum *et al*, 2022). One subcomplex consist of Mic60, Mic19, and in mammals additionally Mic25, and forms contact sites between inner and outer mitochondrial membranes through multiple interactions with outer membrane protein complexes. Moreover, Mic60 induces membrane curvature via an amphipathic helix within its intermembrane space domain (Hessenberger *et al*, 2017; Tarasenko *et al*, 2017). The other subcomplex is composed of large oligomers of Mic10, a small inner membrane protein with an intrinsic membrane-bending activity, together with Mic12/QIL1, Mic26 and Mic27 (Barbot *et al*, 2015; Bohnert *et al*, 2015; Friedman *et al*, 2015; Guarani *et al*, 2015). Both MICOS subcomplexes are necessary for the formation of crista junctions and their physical coupling is largely mediated by Mic12/QIL1 (Guarani *et al*, 2015; Zerbes *et al*, 2016). MICOS deficiency causes the loss of crista junctions and the detachment of cristae from the inner boundary membrane, and loss of function mutations in human patients cause severe mitochondrial pathologies, including a fatal encephalopathy (John *et al*, 2005; Rabl *et al*, 2009; Mun *et al*, 2010; Harner *et al*, 2011; Hoppins *et al*, 2011; Malsburg *et al*, 2011; Guarani *et al*, 2016; Zeharia *et al*, 2016; Benincá *et al*, 2021; Peifer-Weiß *et al*, 2023).

Respiratory chain biogenesis is a highly complicated multi-step process that requires a plethora of dedicated assembly factors. Moreover, individual respiratory chain complexes associate to form supercomplexes of different stoichiometry (Enríquez, 2016; Hartley *et al*, 2019; Rathore *et al*, 2019; Vercellino & Sazanov, 2022). Whereas major assembly steps and intermediates in the biogenesis of NADH dehydrogenase (complex I) and cytochrome *c* oxidase (complex IV) have been identified and characterized (Mick *et al*, 2011; Soto *et al*, 2012; Stroud *et al*, 2016; Guerrero-Castillo *et al*, 2017; Formosa *et al*, 2018; Timón-Gómez *et al*, 2018), comparably little is known about the mechanism of cytochrome *bc*_1_ complex (complex III) assembly. In the yeast *Saccharomyces cerevisiae*, complex III is composed of ten different subunits (Smith *et al*, 2012; Ndi *et al*, 2018; Signes & Fernandez-Vizarra, 2018; Zara *et al*, 2022) and forms dimeric supercomplexes that are found associated with either one or two copies of complex IV (Wittig & Schägger, 2009; Hartley *et al*, 2019; Rathore *et al*, 2019). Complex III assembly is initiated by the translation and membrane insertion of the mitochondrial encoded subunit cytochrome *b* (Cob), which subsequently forms an early core subcomplex together with Qcr7 and Qcr8 (Zara *et al*, 2007; Gruschke *et al*, 2011; 2012). A late complex III assembly intermediate of about 500 kDa was identified that is already dimeric and contains all subunits except the Rieske Fe/S protein (Rip1) and Qcr10 (Zara *et al*, 2009; Conte *et al*, 2015; Stephan & Ott, 2020). Incorporation of these two proteins and formation of supercomplexes with complex IV constitute the final steps in complex III assembly (Cruciat *et al*, 1999; Wagener *et al*, 2011; Atkinson *et al*, 2011; Cui *et al*, 2012; Smith *et al*, 2012; Ndi *et al*, 2018; Kater *et al*, 2020; Tang *et al*, 2020).

Accumulating evidence suggests that mitochondrial membrane architecture and respiratory chain integrity are closely linked. Alterations of cristae morphology lead to defects in respiratory chain supercomplex formation and decreased respiratory capacity (Cogliati *et al*, 2013; 2016; Baker *et al*, 2019; Colina Tenorio *et al*, 2020). In cells with defective MICOS complexes, mitochondrial respiration is considerably reduced and the distribution of respiratory chain complexes in the inner mitochondrial membrane appears to be altered (Malsburg *et al*, 2011; Weber *et al*, 2013; Harner *et al*, 2014; Bohnert *et al*, 2015; Friedman *et al*, 2015; Guarani *et al*, 2015; Anand *et al*, 2020; Rampelt *et al*, 2022). However, the molecular nature of the interaction network that links respiratory chain biogenesis to cristae formation and remodeling has remained enigmatic.

Here we show that MICOS physically associates with respiratory chain (super-)complexes and distinct respiratory chain assembly intermediates. We have identified the so far uncharacterized inner mitochondrial membrane protein Mar26 as an interaction partner of both respiratory chain complexes and MICOS. Mar26 is part of a novel Rip1-containing complex III assembly intermediate and couples this subcomplex to MICOS via the Mic60-Mic19 module. Loss of Mar26 leads to decreased respiratory growth and perturbs late stages of complex III biogenesis, indicating that Mar26 directly contributes to late complex III assembly steps. Recruitment of the Rip1 assembly intermediate to MICOS at crista junctions is required for its faithful assembly. Our results show that MICOS facilitates late complex III assembly by its interaction with the novel assembly factor Mar26.

## RESULTS

### MICOS is physically connected to the respiratory chain

In a previous study we determined the interactome of the MICOS core component Mic60 (Malsburg *et al*, 2011). In addition to the five MICOS components Mic10, Mic12, Mic19, Mic26, Mic27 and subunits of the TOM complex, our analysis identified subunits of the respiratory chain as potential interaction partners of Mic60. To investigate the relationship between MICOS and the respiratory chain in more detail, we purified native protein complexes by affinity chromatography via protein A-tags on either Mic60 or Mic12 after solubilization of mitochondrial membranes with the mild detergent digitonin. Western blot analysis of the elution fractions revealed a specific co-isolation of core subunits of respiratory chain complex III (Cyt1, Qcr8, Rip1) and complex IV (Cox2, Cox9) with both Mic60_ProtA_ and Mic12_ProtA_ (Figure 1A, lanes 4-6). Furthermore, we recovered in the elution fractions the respiratory chain supercomplex-associated proteins Rcf1 and Rcf2 (Strogolova *et al*, 2012; Vukotic *et al*, 2012; Chen *et al*, 2012; Strogolova *et al*, 2019), as well as the complex IV assembly factor Shy1 (Mick *et al*, 2007). As reported previously, Tom40 was efficiently co-isolated only with Mic60_ProtA_, indicating the presence of different Mic60 pools in the inner mitochondrial membrane (Malsburg *et al*, 2011; Bohnert *et al*, 2012).

**Figure 1.**
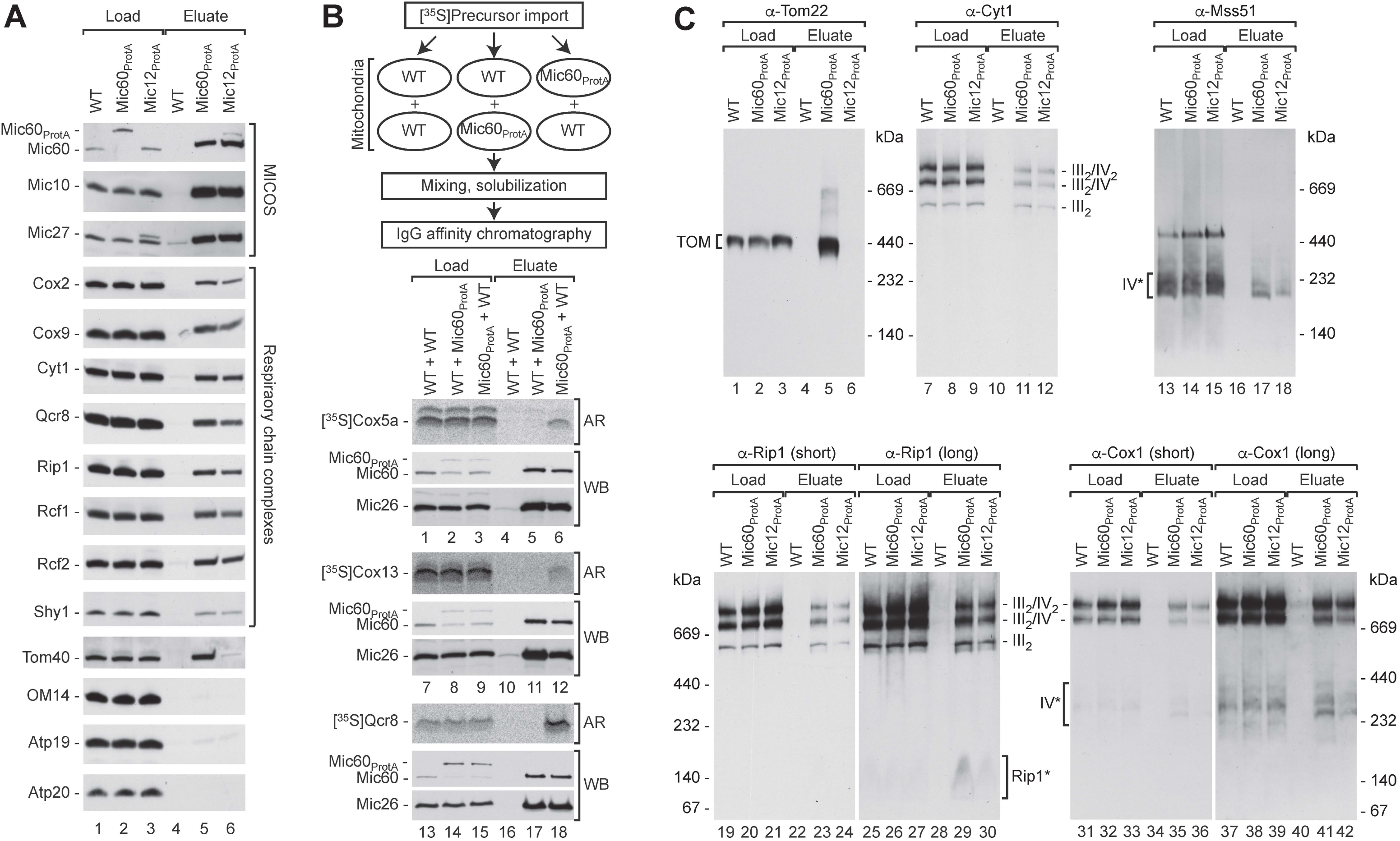
MICOS is connected to the respiratory chain. (A) Protein complexes were purified under native conditions from digitonin-solubilized wild-type (WT), Mic12_ProtA_ and Mic60_ProtA_ mitochondria by IgG affinity chromatography and subsequently analyzed by SDS-PAGE and Western blotting. Load, 1%; Eluate, 100%. Mic60, Mic10, Mic12, Mic27, MICOS subunits; Cox2, Cox9, complex IV subunits; Cyt1, Qcr8, Rip1, complex III subunits; Rcf1, Rcf2, Shy1, respiratory chain assembly factors; Tom40, subunit of the translocase of the outer membrane (TOM); OM14, outer membrane protein; Atp19, Atp20, subunits of the F_1_F_o_-ATP synthase. (B) Indicated [^35^S]labeled preproteins were imported either into wild-type (WT) or Mic60_ProtA_ mitochondria. Mitochondria were re-isolated and mixed with either wild-type or Mic60_ProtA_ mitochondria as indicated. Protein complexes were subsequently purified by IgG affinity chromatography as in (A) and analyzed by SDS-PAGE and Western blot (WB) or autoradiography (AR). Load, 1%; Eluate, 100%. Cox5a, Cox13, complex IV subunits; Qcr8, complex III subunit, Mic26, MICOS subunit. (C) Protein complexes were purified as described in (A) and analyzed by blue native (BN-)PAGE and immunoblotting. Short and long exposures of the blots probed with Rip1 and Cox1 antibodies are shown. Load, 1%; Eluate, 100%. III_2_/IV_2_, III_2_/IV, III_2_, supercomplexes formed by respiratory chain complexes III and IV; IV*, assembly intermediates of complex IV; Cyt1, Rip1, complex III subunits; Rip1*, Rip1-containing assembly intermediate of complex III; Mss51, complex IV assembly factor; Cox1, complex IV subunit; Tom22, TOM subunit (translocase of outer mitochondrial membrane).

To exclude the possibility that MICOS and the respiratory chain components interacted only after detergent-mediated lysis of mitochondria, we performed a post-lysis control experiment: The *in vitro* synthesized and radiolabeled precursors of Cox5a, Cox13 and Qcr8 were imported either into wild-type or Mic60_ProtA_ mitochondria. After re-isolation, preprotein-loaded wild-type mitochondria were mixed with untreated Mic60_ProtA_ mitochondria and vice versa (Figure 1B, flow diagram). Mitochondrial samples were then solubilized with digitonin and MICOS complexes were isolated. Radiolabeled respiratory chain subunits were only co-isolated with tagged Mic60 when they had been imported into Mic60_ProtA_ mitochondria, whereas intrinsic (unlabeled) Mic26 and Mic60 were co-isolated in all cases (Figure 1B, lanes 4-6, 10-12, and 16-18). We conclude that respiratory chain complexes associate with MICOS within the mitochondrial inner membrane and not after lysis. Blue native PAGE analysis of the import reactions confirmed that imported Cox13 and Qcr8 were assembled into respiratory chain supercomplexes (Figure S1, lanes 4-9), whereas the majority of imported Cox5a accumulated in complex IV assembly intermediates as previously reported (Figure S1, lanes 1-3) (Mick *et al*, 2007).

These data suggested that both respiratory chain supercomplexes and assembly factors interact with MICOS. We therefore analyzed the elution fractions of MICOS isolations using tagged Mic60 and Mic12 by blue native PAGE and Western blotting. In line with the SDS-PAGE analysis (Figure 1A), TOM complexes were specifically purified with tagged Mic60, but not with Mic12 as expected (Figure 1C, lanes 4-6). The elution fractions of both Mic60_ProtA_ and Mic12_ProtA_ isolations contained considerable amounts of different respiratory chain supercomplex species composed of complex III and IV, demonstrating that mature respiratory chain supercomplexes interact with MICOS (Figure 1C, lanes 10-12, 22-24, and 34-36). Distinct Cox1-containing assembly intermediates of complex IV have been identified that are associated with specialized assembly factors, like Mss51, not present in mature supercomplexes (Mick *et al*, 2011). These early assembly intermediates were also co-isolated with Mic60_ProtA_ and Mic12_ProtA_ (Figure 1C, lanes 16-18 and 40-42). Interestingly, using antibodies against Rip1 we found in the elution fractions a small Rip1-containing protein complex with an apparent molecular weight of ∼140 kDa that has not been described previously. This complex is barely detectable and thus of low abundance in total mitochondrial extracts, but strongly enriched with purified MICOS complexes (Figure 1C, lanes 22-24 and 28-30). Hence, the small Rip1-containing complex efficiently associates with MICOS and potentially represents a complex III assembly intermediate.

### The Mic60-Mic19 module mediates the connection of MICOS to the respiratory chain

The MICOS complex is composed of two distinct subcomplexes that both contribute to membrane curvature generation at crista junctions: a Mic10-Mic12-Mic26-Mic27 module and a Mic60-Mic19 module that interacts with outer mitochondrial membrane protein complexes to form membrane contacts sites (Barbot *et al*, 2015; Bohnert *et al*, 2015; Friedman *et al*, 2015; Hessenberger *et al*, 2017; Tarasenko *et al*, 2017; Bock-Bierbaum *et al*, 2022). To investigate which MICOS module is responsible for respiratory chain coupling, affinity chromatography and blue native PAGE analysis was performed with Mic60_ProtA_ *mic10*Δ mitochondria. Deletion of MIC10 leads to MICOS disruption and loss of crista junctions (Harner *et al*, 2011; Hoppins *et al*, 2011; Malsburg *et al*, 2011; Bohnert *et al*, 2015). Loss of Mic10 did not abolish the co-isolation of respiratory chain complexes III and IV with tagged Mic60 (Figure 2A, lanes 4-6, 10-12, 16-18; Figure S2). (The efficiency of co-isolation appeared even slightly enhanced in the absence of Mic10.) These data indicate that respiratory chain complexes are coupled to MICOS via the Mic60-Mic19 module and that the Mic10-containing module is not required for respiratory chain interaction of Mic60-Mic19. In line with this conclusion, the amounts of respiratory chain complexes co-purified with Mic12_ProtA_ in the absence of Mic60 or Mic10 were only slightly above the unspecific background (Figure 2B, lanes 5-8, 13-16). In Mic10-deficient mitochondria, Mic60 is still present, but not associated with the Mic12-containing MICOS subcomplex (Bohnert *et al*, 2015). We conclude that the Mic60-Mic19 module recruits respiratory chain complexes to MICOS at crista junctions.

**Figure 2.**
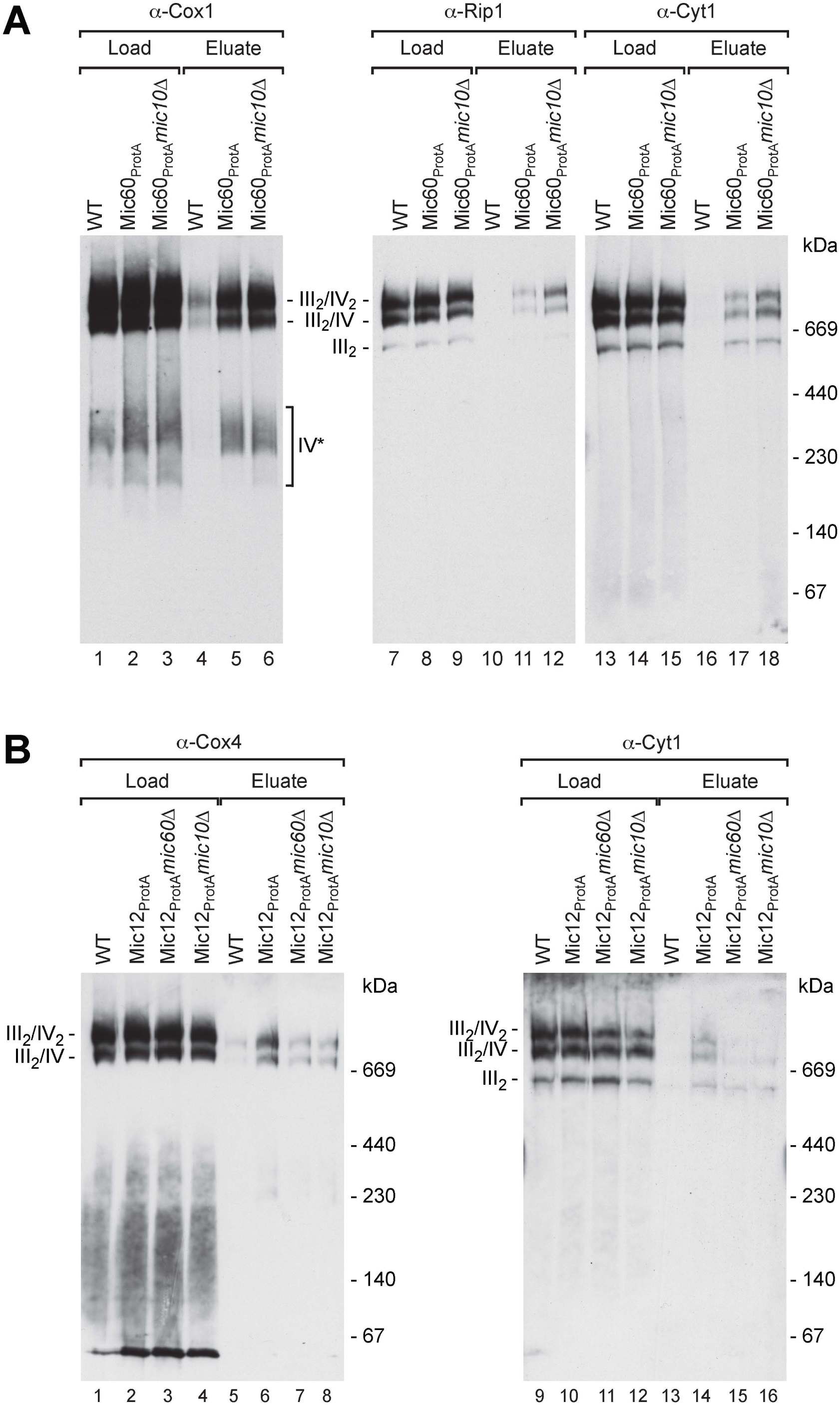
The Mic60-Mic19 module mediates MICOS connection to the respiratory chain. (A) Protein complexes were purified from digitonin-solubilized wild-type (WT), Mic60_ProtA_ and Mic60_ProtA_ *mic10*Δ mitochondria by IgG affinity chromatography and subsequently analyzed by BN-PAGE and Western blotting as in Figure 1C. Load 1%, Eluate 100%. III_2_/IV_2_, III_2_/IV, III_2_, supercomplexes of respiratory chain complexes III and IV; IV*, assembly intermediates of complex IV. (B) Protein complexes purified from wild-type, Mic12_ProtA_, Mic12_ProtA_ *mic60*Δ and Mic12_ProtA_ *mic10*Δ mitochondria were analyzed as in (A). Load 1%, Eluate 100%. Cox4, complex IV subunit.

### Mar26 is associated with MICOS and respiratory chain complexes

To further analyze the connection between respiratory chain and MICOS complexes, we used yeast cells expressing TAP-tagged Cor1 of complex III (van der Laan *et al*, 2006). Stable isotope labeling with amino acids in cell culture (SILAC) combined with high-resolution mass spectrometry (MS) (Malsburg *et al*, 2011) was used to identify respiratory chain-interacting proteins (Table S1). Besides the genuine subunits of respiratory chain complexes III and IV, we co-isolated components of the presequence translocase of the inner mitochondrial membrane and a number of metabolite carrier proteins as previously reported (van der Laan *et al*, 2006; Claypool *et al*, 2008; Dienhart & Stuart, 2008; Mehnert *et al*, 2014) (Figure 3A). Moreover, MICOS components were also significantly enriched with tagged respiratory chain complexes, independently confirming the physical association of these protein machineries (Figure 3A). Interestingly, we also found a strong enrichment of a so far uncharacterized mitochondrial protein encoded by the *S. cerevisiae* open reading frame YER182W that was named Found in mitochondrial proteome 10 (Fmp10). We will below refer to this protein as Mar26 (MICOS associated respiratory chain factor of 26 kDa). Our SILAC/MS-based analysis of the Mic60 interactome had previously identified Mar26 as a putative partner protein of MICOS (Malsburg *et al*, 2011). (For comparison, we have included a new analysis of the original Mic60_ProtA_ SILAC/MS data set performed as for Cor1_TAP_ isolations in Table S1.)

**Figure 3.**
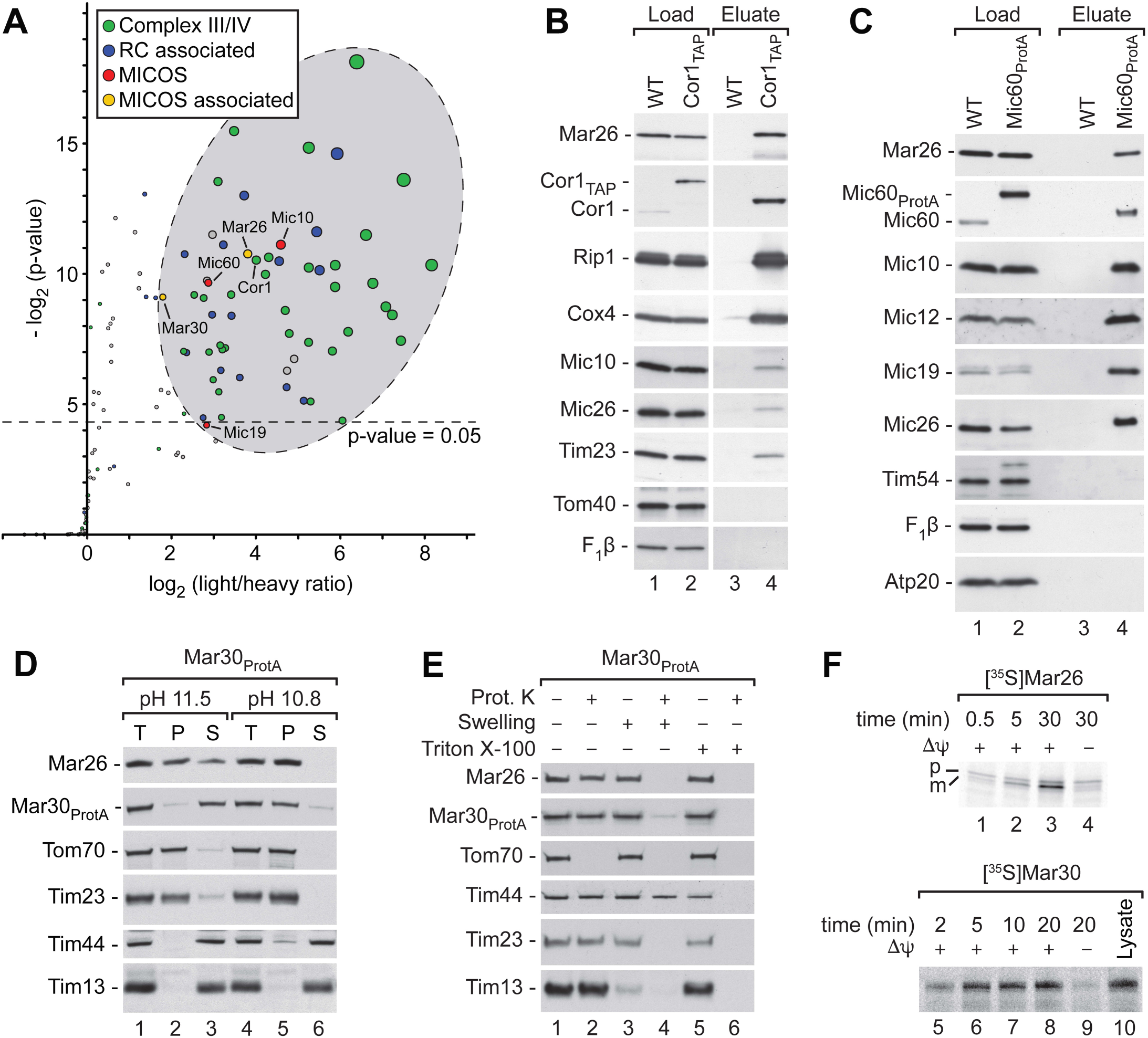
Identification and characterization of Mar26 (Yer182w) and Mar30 (Ybl095w). (A) SILAC-based MS analysis of Cor1_TAP_ interactors. Structural subunits of complex III and IV and known proteins required for complex III and IV biogenesis (green), proteins described previously to associate with respiratory chain (RC) complexes III and IV (blue), MICOS components (red) and newly identified MICOS associated respiratory chain factors (yellow) are indicated. Additional proteins labeled in grey have not been reported before to associate with the respiratory chain (Rad4, Tao3, Ynl122c). A complete list of identified proteins can be found in Table S1. Cor1, complex III subunit. (B) Protein complexes were purified from wild-type (WT) and Cor1_TAP_ mitochondria by IgG affinity chromatography and analyzed by SDS-PAGE and Western blotting. Load, 4%; Eluate, 100%. Tim23, subunit of the TIM23 presequence translocase of the inner membrane. F_1_β, β subunit of F_1_F_o_-ATP synthase. (C) Protein complexes purified from wild-type and Mic60_ProtA_ mitochondria were analyzed as in (B). Load, 4%; Eluate, 100%. Mic19, MICOS subunit; Tim54, subunit of the TIM22 carrier translocase of the inner membrane. (D) Membrane association of Mar26 und Mar30 was tested by alkaline extraction and subsequent ultracentrifugation. T, total; P, pellet; S, supernatant. Tom70, TOM component; Tim44, TIM23 component; Tim13, small Tim chaperone of the intermembrane space. (E) The submitochondrial localization of Mar26 and Mar30 was assessed by protease K accessibility. Mar30_ProtA_ mitochondria were either incubated in SEM buffer (intact mitochondria), subjected to hypoosmotic swelling or solubilized with 0.5% (v/v) Triton X-100 and subsequently treated with proteinase K (Prot. K) as indicated. (F) [^35^S]Mar26 and [^35^S]Mar30 preproteins were imported for the indicated time periods into wild-type mitochondria in the presence or absence of an inner membrane potential (Δψ). Non-imported preproteins were removed by proteinase K treatment. Samples were analyzed by SDS-PAGE and autoradiography. *In vitro* synthesized preprotein (Lysate) is shown for comparison where indicated. p, precursor; m, mature protein.

We generated an antiserum against a C-terminal peptide of Mar26. Western blot analysis of Cor1_TAP_ and Mic60_ProtA_ isolations confirmed that Mar26 is efficiently co-isolated with both respiratory chain complexes and MICOS (Figures 3B and 3C). We asked if the presence of a functional respiratory chain is necessary for the stable expression and accumulation of Mar26 in mitochondria. We generated a Mic60_ProtA_ yeast strain lacking mitochondrial DNA (*rho*−). Respiratory chain complex subunits that are encoded by nuclear genes, like Rip1, Qcr8, Cox4, or Cox9, are still detectable in isolated mitochondria of *rho*− cells (Figure S3A) but cannot assemble into functional respiratory chain complexes due to the lack of the mitochondrial encoded subunits. Therefore, their levels are considerably reduced compared to *rho*+ mitochondria. In contrast, the steady state levels of Mar26 as well as the outer membrane preprotein receptor Tom70 were comparable in *rho*+ and *rho*− mitochondria, and those of the MICOS core components Mic60 and Mic10 were only mildly reduced (Figure S3A). We conclude that accumulation of Mar26 in mitochondria does not depend on the presence of respiratory chain complexes. Affinity chromatography experiments furthermore revealed that Mar26 is still efficiently co-isolated with tagged Mic60 in the *rho*− mitochondria lacking respiratory chain complexes (Figure S3B).

### Mar26 and its paralog Mar30 are inner mitochondrial membrane proteins

Homology searches revealed that MAR26 has a paralog in *S. cerevisiae* encoded by the open reading frame YBL095W. Like Mar26, the encoded protein was found associated with mitochondrial ribosomes (and therefore named Mrx3) (Kehrein *et al*, 2015), identified as a potential MICOS-interacting protein (Malsburg *et al*, 2011; Jin *et al*, 2015), and found enriched in our Cor1_TAP_ isolations (Figure 3A). We termed the encoded protein Mar30 (MICOS associated respiratory chain factor of 30 kDa). Mar26 and Mar30 share 49.3% similarity and 16.4% identity (Figure S3C). We did not succeed in generating antibodies against Mar30 and instead fused a protein A-tag to the protein for biochemical analysis. Alkaline extraction of mitochondrial membranes showed that both Mar26 and Mar30_ProtA_ remained in the pellet fraction together with other integral membrane proteins, like Tom70 or Tim23, at pH 10.8 (Figure 3D). Mar26 remained in the pellet also at pH 11.5, whereas Mar30 was found in the supernatant fraction at pH 11.5 (Figure 3D). This behavior is in agreement with the predicted presence of one transmembrane segment in each protein and the low predicted hydrophobicity of the potential Mar30 transmembrane segment. The submitochondrial localization of both proteins was tested by a protease accessibility assay (Figure 3E). When proteinase K is added to intact mitochondria, only surface-exposed outer membrane proteins, like Tom70, are digested. Hypo-osmotic swelling opens up the outer membrane and allows access of the protease to proteins that expose soluble domains to the intermembrane space, like Tim23 or Tim13. Under these conditions, also Mar26 and Mar30_ProtA_ were proteolytically degraded. Matrix-exposed proteins, like Tim44, remained protected and were only degraded upon detergent solubilization of mitochondria (Figure 3E). We conclude that Mar26 and Mar30 are both inner membrane proteins exposed to the intermembrane space.

No presequence cleavage site is predicted for Mar30. In agreement with this prediction, the *in vitro* synthesized and radiolabeled Mar30 preprotein was imported into isolated mitochondria to a protease-protected localization in a membrane-potential-dependent manner, but not proteolytically processed. Radiolabeled Mar26 was likewise imported into mitochondria, but processed to a mature form indicative of presequence cleavage (Figure 3F). This is consistent with the finding that mature Mar26 is generated by the successive processing of 12 + 1 N-terminal residues by the presequence peptidase MPP and the intermediate cleaving peptidase Icp55, respectively (Vögtle *et al*, 2009).

### Mar26, MICOS and respiratory chain complexes III and IV form an interaction network

We performed Mar26 and Mar30 affinity purifications using Protein A-tagged variants of these proteins expressed from their native chromosomal loci. Initial SILAC-MS analysis of the Mar26 interactome revealed a strong interaction with Mar30 and confirmed its close association with MICOS and respiratory chain supercomplexes (Table S1). Further Western blot analysis showed that Mar26_ProtA_ and Mar30_ProtA_ co-purified MICOS components as well as the central TOM complex subunit Tom40 with similar efficiency (Figure 4A, lanes 4-6). MICOS components were recovered together with Mar26_ProtA_ und Mar30_ProtA_ also from *rho*− mitochondria showing that the association of both Mar26 and Mar30 with MICOS does not depend on the presence of respiratory chain complexes (Figures S3D). Using *rho*+ mitochondria, subunits of complex III, like Rip1 or Cyt1, and of complex IV, like Cox2, were considerably more abundant in the elution fractions of Mar26_ProtA_ isolations compared to Mar30_ProtA_ (Figure 4A, lanes 4-6). Accordingly, mature respiratory chain supercomplexes, as detected with antibodies against the complex III subunit Cyt1, were co-isolated with Mar26_ProtA_, but only in minor amounts with Mar30_ProtA_ (Figure 4A, lanes 10-12). Thus, Mar26 and Mar30 are both MICOS-binding proteins, but the connection to the respiratory chain is clearly stronger for Mar26.

**Figure 4.**
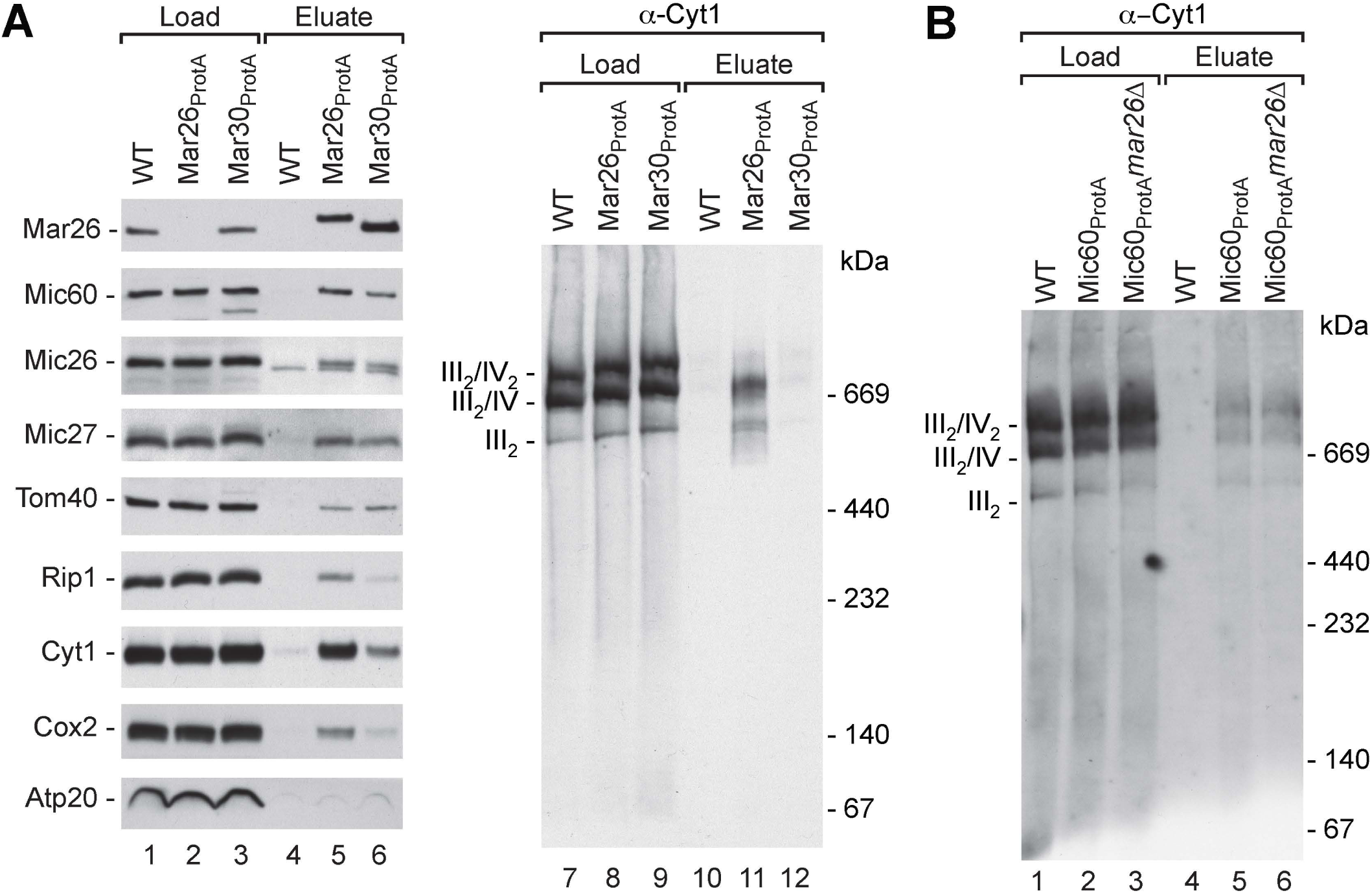
Mar26, MICOS and respiratory chain complexes III and IV form an interaction network. (A) Protein complexes of wild type (WT), Mar26_ProtA_ and Mar30_ProtA_ were purified under native conditions and analyzed by SDS-PAGE or BN-PAGE and Western blotting. Load, 1%; Eluate, 100%. III_2_/IV_2_, III_2_/IV, III_2_, supercomplexes formed by respiratory chain complexes III and IV. (B) Analysis of respiratory chain supercomplexes co-purified with Mic60_ProtA_ in the presence or absence of Mar26 as indicated by BN-PAGE and Western blotting. Load, 1%; Eluate, 100%.

We next tested whether Mar26 may be involved in the recruitment of respiratory chain supercomplexes to MICOS. However, affinity purification experiments with mitochondria from cells expressing Mic60_ProtA_ in the absence of Mar26 revealed that the recovery of mature respiratory chain supercomplexes in the elution fractions was similar for Mic60_ProtA_ and Mic60_ProtA_ *mar26*Δ mitochondria (Figure 4C, lanes 4-6). Thus, Mar26 is not required for coupling of mature respiratory chain supercomplexes to MICOS.

### Lack of Mar26 leads to alterations of complex III

What is the functional role of Mar26 in mitochondria? To find out whether Mar26 and its paralog are important for respiratory growth, we tested growth of the deletion mutants on fermentable and respiratory medium containing either glucose or ethanol and glycerol as carbon sources. Notably, *mar26*Δ and *mar30*Δ cells showed a strong growth defect specifically on respiratory medium (Figure 5A). Based on our interactome analysis (Figure 3A, Table S1), we first asked whether cristae architecture was affected in these mutants and consequently examined mutant mitochondria by electron microscopy. Mitochondrial ultrastructure, cristae morphology and the prevalence of crista junctions were indistinguishable from the corresponding wild-type indicating that MICOS function is not considerably impaired in the absence of Mar26 (Figure 5B). The steady-state levels of MICOS subunits and many other mitochondrial proteins tested were also normal in *mar26*Δ mitochondria (Figure 5C). Respiratory chain supercomplexes and Mss51-containing complex IV assembly intermediates were similarly abundant in wild-type, *mar26*Δ, and *mic60*Δ mutant mitochondria (Figure 5D, lanes 1-5 and 11-15; Figure S4A). However, longer exposures of blue native PAGE Western blots probed with antibodies against Cyt1 of complex III revealed the accumulation of a ∼500 kDa Cyt1-containing complex in the absence of Mar26 (Figure 5D, lanes 6-10). The apparent molecular weight of this protein complex resembles that of the 500 kDa late assembly intermediate of complex III that accumulates in the absence of Rip1 (Figure S4B lane 3) (Zara *et al*, 2009). In agreement with the literature (Conte *et al*, 2015), we observed that the 500 kDa complex present in *rip1*Δ, which is a dimer of partially assembled complex III (Stephan & Ott, 2020), already associates with complex IV to form (non-functional) respiratory chain supercomplexes, albeit with reduced efficiency (Figure S4C, lanes 3 and 6). Strikingly, the small ∼140 kDa complex containing Rip1 that we found associated and enriched with purified MICOS complexes (Figure 1C) was not detectable in the *mar26*Δ mutant mitochondria, indicating that Mar26 is likely part of this small Rip1 complex (Figure 5D, lanes 16-17). In line with this finding, Mar26 forms a low molecular weight complexes in mitochondria (Figure S4C). Although this complex efficiently binds to MICOS via the Mic60-Mic19 module, its formation and stability does not require Mic60, because it is present at normal levels in *mic60*Δ mitochondria (Figure 5D, lanes 19-20).

**Figure 5.**
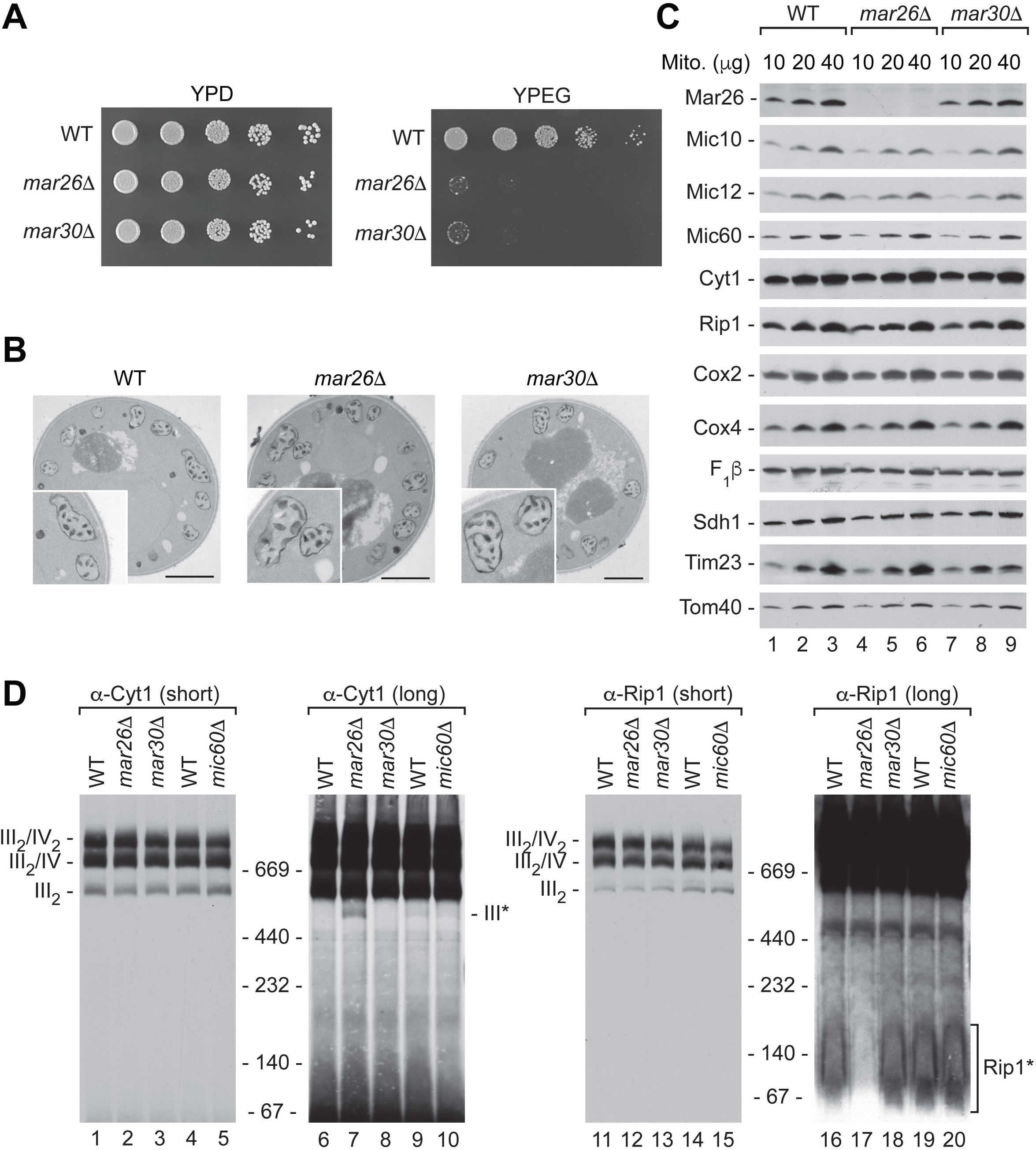
Lack of Mar26 but not of Mar30 leads to alterations of complex III. (A) WT, *mar26*Δ and *mar30*Δ yeast cells were spotted as 5-fold serial dilutions on YPD or YPEG plates and incubated at 34°C. (B) Electron microscopy analysis of wild-type (WT), *mar26*Δ and *mar30*Δ cells using diaminobenzidine staining of mitochondrial cristae. Bar, 1 μm. (C) Steady state protein levels of WT, *mar26*Δ and *mar30*Δ mitochondria were analyzed by SDS-PAGE and Western blotting. Sdh1, complex II component. (D) Respiratory chain supercomplexes in WT, *mar26*Δ, *mar30*Δ and *mic60*Δ mitochondria were analyzed by BN-PAGE and immunoblotting. III_2_/IV_2_, III_2_/IV, III_2_, supercomplexes formed by respiratory chain complexes III and IV; III*, 500 kDa late assembly intermediate of complex III; Rip1*, Rip1-containing assembly intermediate of complex III.

### Mar26 plays a regulatory role in complex III biogenesis

Our findings strongly indicate that Mar26 may play a direct role in complex III biogenesis. To test this idea we imported radiolabeled preproteins of respiratory chain complex subunits into isolated wild-type and *mar26*Δ mutant mitochondria and followed their assembly into respiratory chain supercomplexes by blue native PAGE. Incorporation of [^35^S]Qcr10 that assembles at the final stages of complex III biogenesis with the 500 kDa intermediate (Smith *et al*, 2012; Ndi *et al*, 2018) was not affected in *mar26*Δ mitochondria (Figure S5A). Similarly, import and assembly of the complex IV subunits [^35^S]Cox13 into mature respiratory chain supercomplexes (Figure S5B), [^35^S]Cox5a into Cox1-containing assembly intermediates (Figure S5C), and of the late assembly factor [^35^S]Rcf1 (Figure S5D) was comparable in wild-type and *mar26*Δ mutant mitochondria. In marked contrast, incorporation of [^35^S]Qcr8 into respiratory chain supercomplexes was strongly impaired in the absence of Mar26 (Figure 6A, lanes 4-6). Instead, radiolabeled Qcr8 accumulated in a complex of ∼500 kDa. This assembly defect resembles the behavior of Qcr8 when imported into *rip1*Δ mitochondria (Figure 6A, lanes 10-15), identifying the radiolabeled band as the 500 kDa late assembly intermediate of complex III. Surprisingly, radiolabeled, imported [^35^S]Rip1 assembled more efficiently into respiratory chain supercomplexes in the absence of Mar26 (Figure 6B, lanes 4-6). Such increased labeling of respiratory chain supercomplexes is also observed when [^35^S]Rip1 is imported into *rip1*Δ mitochondria (Figure 6B, lanes 10-12) (Wagener *et al*, 2011).

**Figure 6.**
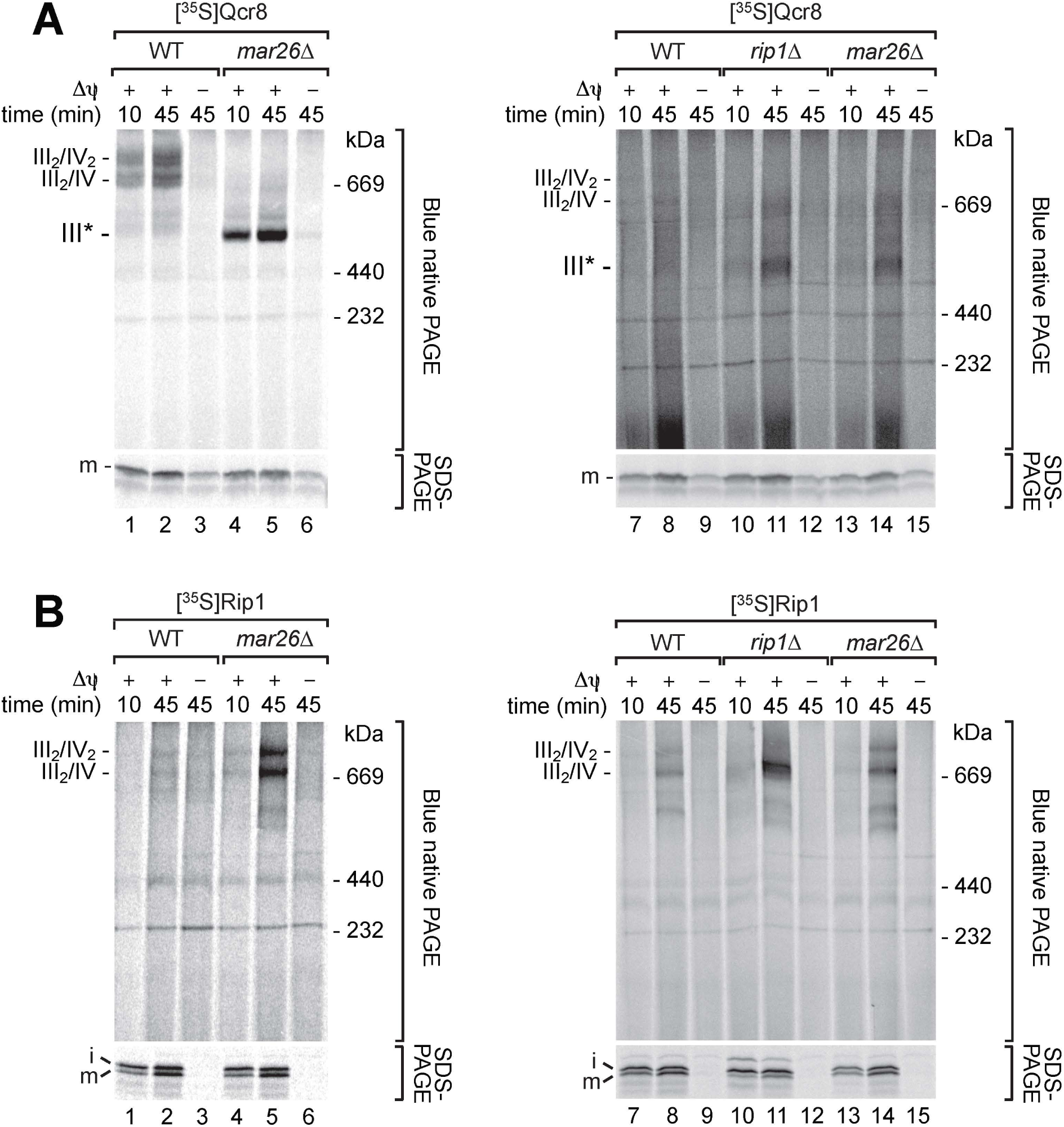
Mar26 plays a regulatory role in complex III biogenesis. (A) Radiolabeled Qcr8 preprotein was imported into mitochondria isolated from wild-type (WT) or *mar26*Δ cells grown on respiratory medium (lanes 1-6), or from WT, *mar26*Δ or *rip1*Δ cells grown on fermentative medium (lanes 7-15). Mitochondria were subsequently solubilized in digitonin-containing buffer and analyzed by SDS-PAGE or BN-PAGE and visualized by autoradiography. III_2_/IV_2_, III_2_/IV, supercomplexes of respiratory chain complexes III and IV; III*, 500 kDa late assembly intermediate of complex III; Δψ, membrane potential; m, mature protein. (B) Radiolabeled Rip1 preprotein was imported into mitochondria and analyzed as in (A). i, intermediate.

The accelerated assembly of Rip1 in *mar26*Δ and *rip1*Δ mitochondria likely results from an accumulation of pre-assembled 500 kDa intermediates *mar26*Δ (Figure 5D). Radiolabeled Rip1 takes over the role of the intrinsic protein that is unavailable (*rip1*Δ) or not sufficiently disposed (*mar26*Δ) for assembly into mature respiratory chain supercomplexes. Thus, Mar26 appears to regulate the assembly of incoming Rip1 with the 500 kDa late assembly intermediate. We propose that the Mar26-dependent small Rip1 complex of ∼140 kDa provides a constant pool of assembly-competent Rip1 for the late assembly stages of complex III biogenesis.

### Mar26 recruits a small Rip1-containing complex to MICOS for efficient assembly

We have demonstrated that Mar26 is a partner protein of MICOS and is also involved in late stages of complex III assembly, probably by modulating the incorporation of Rip1. To ask whether these two findings reflect a functional connection, we examined the co-isolation of individual MICOS and respiratory chain subunits from Mic60_ProtA_ *mar26*Δ mutant mitochondria in detail (Figure 7A). We observed that among the respiratory chain subunits interacting with MICOS, solely Rip1 was co-isolated in reduced amounts when Mar26 was absent (Figure 7A, 16-18). The recovery in the elution fractions of all other tested MICOS and respiratory chain subunits was not changed for Mic60_ProtA_ *mar26*Δ compared to Mic60_ProtA_ mitochondria. Strikingly, overexpression of Mar26 in mitochondria increased not only the amount of Mar26 co-isolated with Mic60_ProtA_ (Figure 7A, lanes 22-24), but the amount of Rip1 associated with tagged Mic60 was also dramatically elevated (Figure 7A, lanes 22-24). The co-purification of all other MICOS and respiratory chain subunits tested was not affected by increased Mar26 levels (Figure 7A), and Mar26 overexpression did not cause any defects in respiratory growth (Figure S6A-B). Since the Mar26-dependent small Rip1-containing complex of ∼140 kDa interacts with MICOS (Figure 1C, lane 29), we analyzed the elution fractions of complex isolations from *mar26*Δ or Mar26 overexpression by BN-PAGE. In line with a Mar26-dependent recruitment of Rip1 to MICOS, our BN-PAGE analysis confirmed that increased levels of Mar26 resulted in a strong increase of the amounts of the small Rip1-containing complex in the elution fractions (Figure 7B, lanes 4-6). We asked whether recruitment of the Rip1 intermediate to MICOS is functionally significant. Loss of Mic60 does not result in accumulation of the late complex III assembly intermediate observed in *mar26*Δ mitochondria, nor is formation of the Mar26-dependent Rip1 intermediate affected in *mic60*Δ mitochondria (Figure 5D), indicating that the Rip1-related function of Mar26 does not depend on MICOS. Despite this, we found that assembly of newly imported Rip1 into mitochondria lacking Mic10 or Mic60 was impaired, and Rip1 instead accumulated in a low molecular weight intermediate (Figure 7C, lanes 1-9). Moreover, assembly of Qcr10, which is incorporated into complex III after Rip1, also displayed an assembly defect in MICOS-deficient mitochondria (Figure 7C, lanes 10-18). In contrast, the early assembling subunit Qcr8 did not display a decreased assembly efficiency in MICOS mutant mitochondria (Figure S6C).

**Figure 7.**
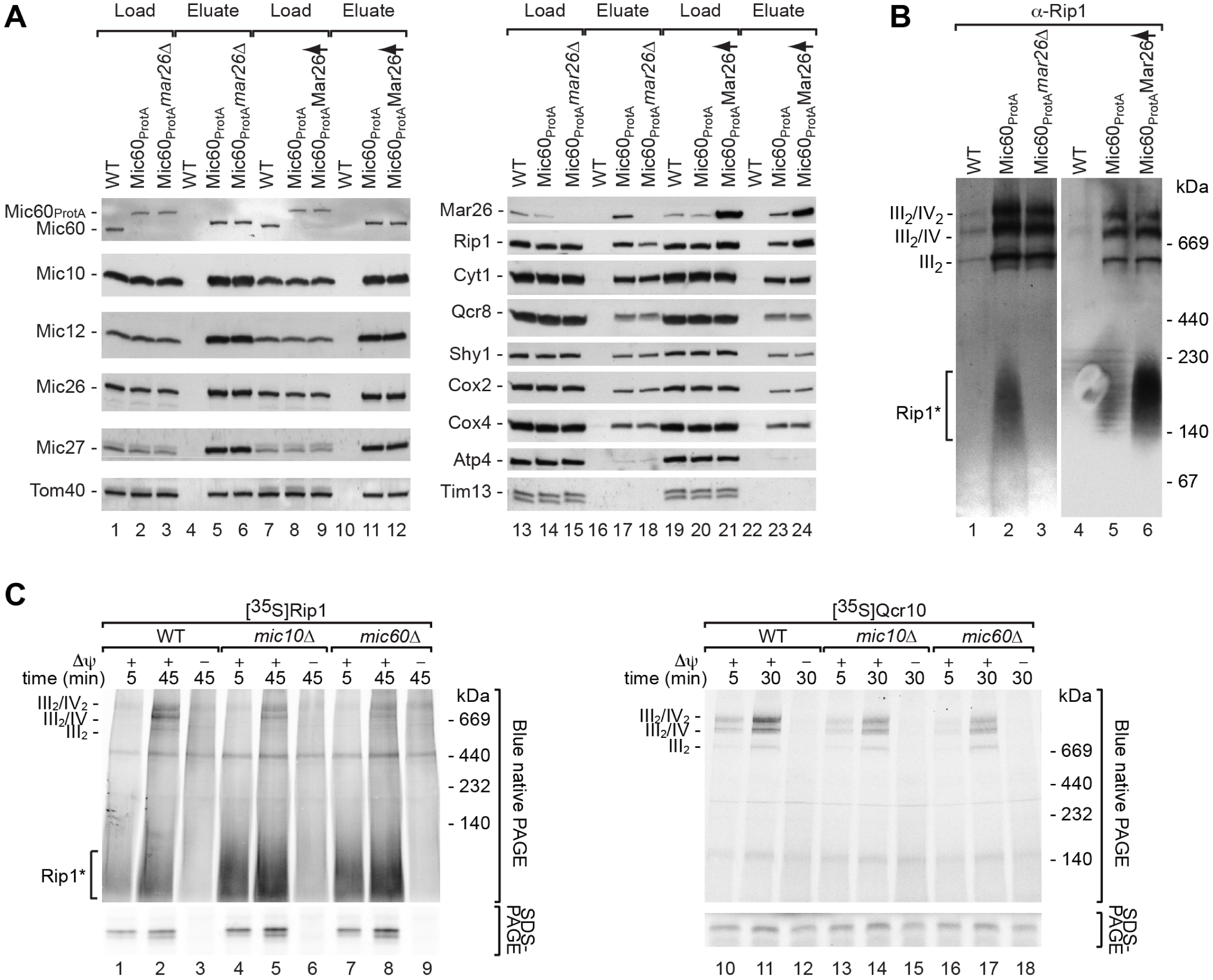
Mar26 recruits Rip1 to MICOS for facilitated assembly. (A) Protein complexes were purified from wild-type (WT), Mic60_ProtA_, Mic60_ProtA_ *mar26*Δ or Mic60_ProtA_ Mar26 overexpression (↑) mitochondria by IgG affinity chromatography. Samples were analyzed by SDS-PAGE and Western blotting. Load, 4 % (lanes 1-12) or 1 % (lanes 13-24); Eluate, 100%. Atp4, F_1_F_o_-ATP synthase subunit. (B) Protein complex isolations were performed as in (A). Eluates were analyzed by BN-PAGE and immunoblotting. III_2_/IV_2_, III_2_/IV, III_2_, supercomplexes formed by respiratory chain complexes III and IV; Rip1*, Rip1-containing assembly intermediate of complex III. (C) Radiolabeled Rip1 or Qcr10 preprotein was imported into mitochondria isolated from wild-type (WT), *mic10*Δ or *mic60*Δ cells. Mitochondria were subsequently solubilized in digitonin-containing buffer and analyzed by SDS-PAGE or BN-PAGE and visualized by autoradiography. III_2_/IV_2_, III_2_/IV, supercomplexes of respiratory chain complexes III and IV; Δψ, membrane potential.

We conclude that Mar26 interacts with a pool of unassembled Rip1 and connects it to the Mic60-Mic19 module of MICOS. In this way, Mar26 physically links late stages of complex III biogenesis to the MICOS complex at crista junctions. The interaction of the Rip1 assembly intermediate with an intact MICOS is important for its efficient assembly, demonstrating that MICOS facilitates late complex III assembly steps.

## DISCUSSION

Most respiratory chain complex subunits are nuclear encoded and inserted into the inner membrane in the boundary region. The assembly of these proteins with each other and with mitochondrial encoded subunits must be tightly coordinated in space and time (Richter-Dennerlein *et al*, 2016). Individual proteins or pre-assembled subcomplexes have to cross the permeability barrier at the crista junction to assemble into respiratory chain supercomplexes (Stoldt *et al*, 2018). Our results indicate that the MICOS complex at crista junctions plays an important role in respiratory chain biogenesis by facilitating submitochondrial protein sorting and coordination of complex assembly steps.

Recent studies revealed a functional link between MICOS and respiratory chain complexes of mitochondria, but the molecular basis of this connection has remained unclear (Harner *et al*, 2014; Bohnert *et al*, 2015; Friedman *et al*, 2015; Guarani *et al*, 2015). In this study we show that respiratory chain supercomplexes are specifically co-purified with MICOS complexes, demonstrating a physical association of these two crucial mitochondrial protein machineries. Defined assembly intermediates of respiratory chain complexes are present in mitochondria at low levels. Interestingly, some assembly intermediates such as the Mar26-dependent Rip1 intermediate appear to be enriched at MICOS complexes.

Our comprehensive MICOS and respiratory chain interactome analysis identified the so far uncharacterized protein Mar26 as a shared interaction partner. Deletion of MAR26 did not interfere with the coupling of respiratory chain supercomplexes to MICOS. Instead, we found that Mar26 plays a specific role in the biogenesis of complex III. We identified a small Rip1-containing subcomplex that likely represents an assembly intermediate and is only detectable in the presence of Mar26. The small Rip1 complex accumulates at MICOS and is connected to MICOS via Mar26. Because respiratory chain supercomplexes are still formed *in vivo* and only small amounts of partially assembled complex III accumulate in *mar26*Δ mutant mitochondria, Mar26 cannot be strictly required for complex III formation. However, our *in vitro* import and assembly assays shed light on a possible role of Mar26. In these experiments only individual subunits are imported into isolated mitochondria, whereas *in vivo* all subunits necessary for assembly of mature respiratory chain complexes are usually supplied. The subunits must bind in a defined order to avoid disrupting the assembly process. Many dedicated assembly factors are known that orchestrate the assembly of respiratory chain complexes. We propose that Mar26 plays a specific role in the late assembly steps of complex III. Our results suggest that Mar26 interacts with unassembled Rip1 and may constantly maintain a small, assembly-competent pool of this subunit. In *mar26*Δ mutant mitochondria, this pool of unassembled Rip1 is absent and a late complex III intermediate accumulates, leading to defective assembly of other subunits, like Qcr8, if supplied individually to the assembly line. Alternatively, individual subunits may be exchanged in existing complexes, for instance for repair or as part of protein complex dynamics. We propose that the Mar26-bound pool of Rip1 becomes critical under conditions where the balanced supply of subunits is disturbed - a condition that is mimicked in our *in vitro* assembly assay. In living cells, such unbalanced situations may occur stochastically with increased frequency when particularly high rates of respiratory chain complex biogenesis are required.

Our finding that respiratory chain intermediates interact with MICOS suggests that crista junctions may be a central hub for respiratory chain biogenesis. In fact, we demonstrate that beyond the role of Mar26 in Rip1 assembly, Mar26-dependent recruitment of the Rip1 intermediate to an intact MICOS is important for efficient Rip1 assembly. Upon import into MICOS-deficient mitochondria, Rip1 accumulates in an intermediate, likely the Mar26-dependent assembly intermediate, and incorporation into complex III is impaired. Delayed Rip1 assembly likely results from loss of the interaction with the Mic60 subcomplex, or from the Rip1 intermediate becoming trapped at a defunct MICOS complex in *mic10*Δ mitochondria. However, MICOS is not globally required for efficient complex III biogenesis since Qcr8 assembly is unaffected in MICOS mutants. Thus, the observed defects reflect a specific function of MICOS rather than a general disruption resulting from altered cristae architecture. In addition to direct actions of MICOS in recruiting assembly intermediates, the strong membrane curvature at crista junctions and the accumulation of specific phospholipids at these sites may also facilitate certain assembly steps (Horvath *et al*, 2015; Rampelt *et al*, 2017; Basu Ball *et al*, 2017).

We find that Mar26 is involved in the coordination of complex III assembly in space and time. Such a coordination requires not only chaperone-like proteins that facilitate the incorporation of individual subunits, but also factors that prevent the premature assembly of subunits. Our data indicate that Mar26 recruits a fraction of unassembled Rip1 into a waiting position at MICOS, where specific assembly intermediates of respiratory chain complexes accumulate and may pick up further components.

We conclude that MICOS functions as a coordinating hub in the sorting of mitochondrial inner membrane proteins between boundary and cristae membrane domains. Our study provides the first indications how mitochondrial ultrastructure and inner membrane protein biogenesis and distribution are linked to each other on the molecular level, and paves the way for further studies on the spatial and temporal organization of mitochondrial protein complex biogenesis.

## SUPPLEMENTAL INFORMATION

Supplemental information includes six figures and three tables.

## SUPPLEMENTAL FIGURE AND TABLE LEGENDS

**Figure S1.**
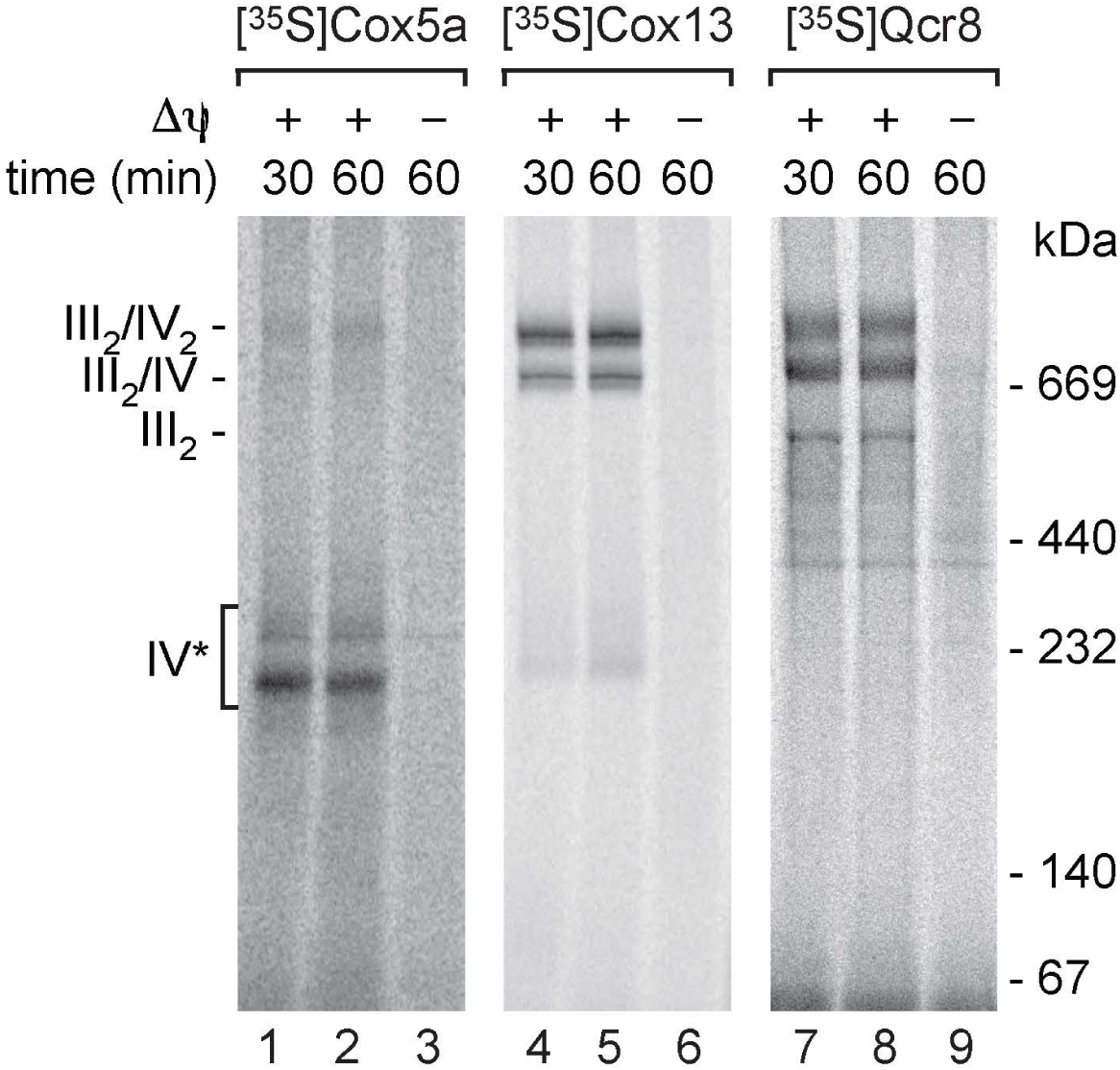
Assembly of radiolabeled respiratory chain preproteins. The indicated radiolabeled preproteins were imported into wild-type mitochondria. Upon solubilized in digitonin-containing buffer samples were analyzed by blue native PAGE and autoradiography. III_2_/IV_2_, III_2_/IV, III_2_, supercomplexes of respiratory chain complexes III and IV; IV*, complex IV and assembly intermediates thereof; Δψ, membrane potential.

**Figure S2.**
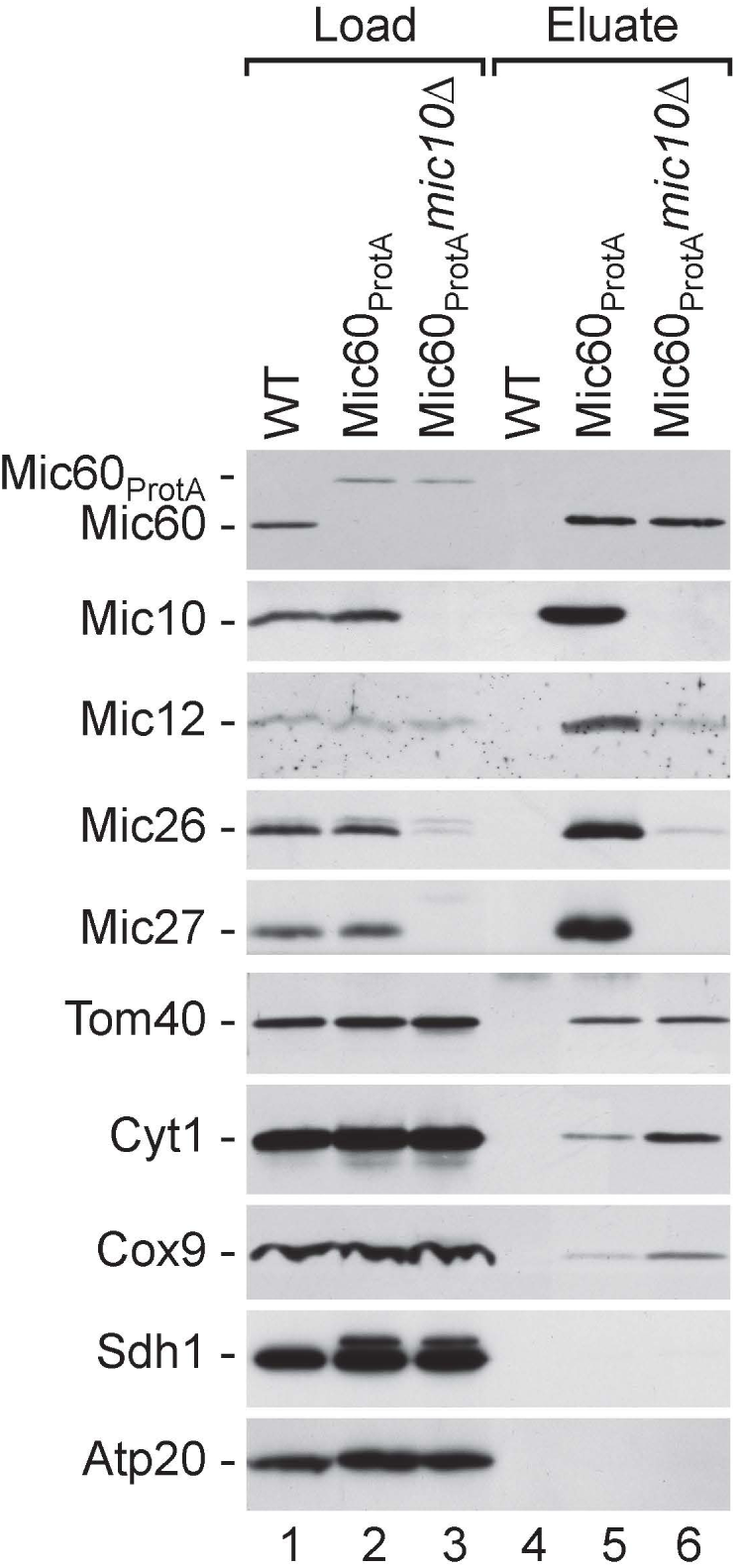
MIC10 deletion leads to dissociation of MICOS. IgG affinity purification was performed with wild-type (WT), Mic60_ProtA_ and Mic60_ProtA_ *mic10*Δ mitochondria and samples were analyzed by SDS-PAGE and immunoblotting. Load, 4%; Eluate, 100%.

**Figure S3.**
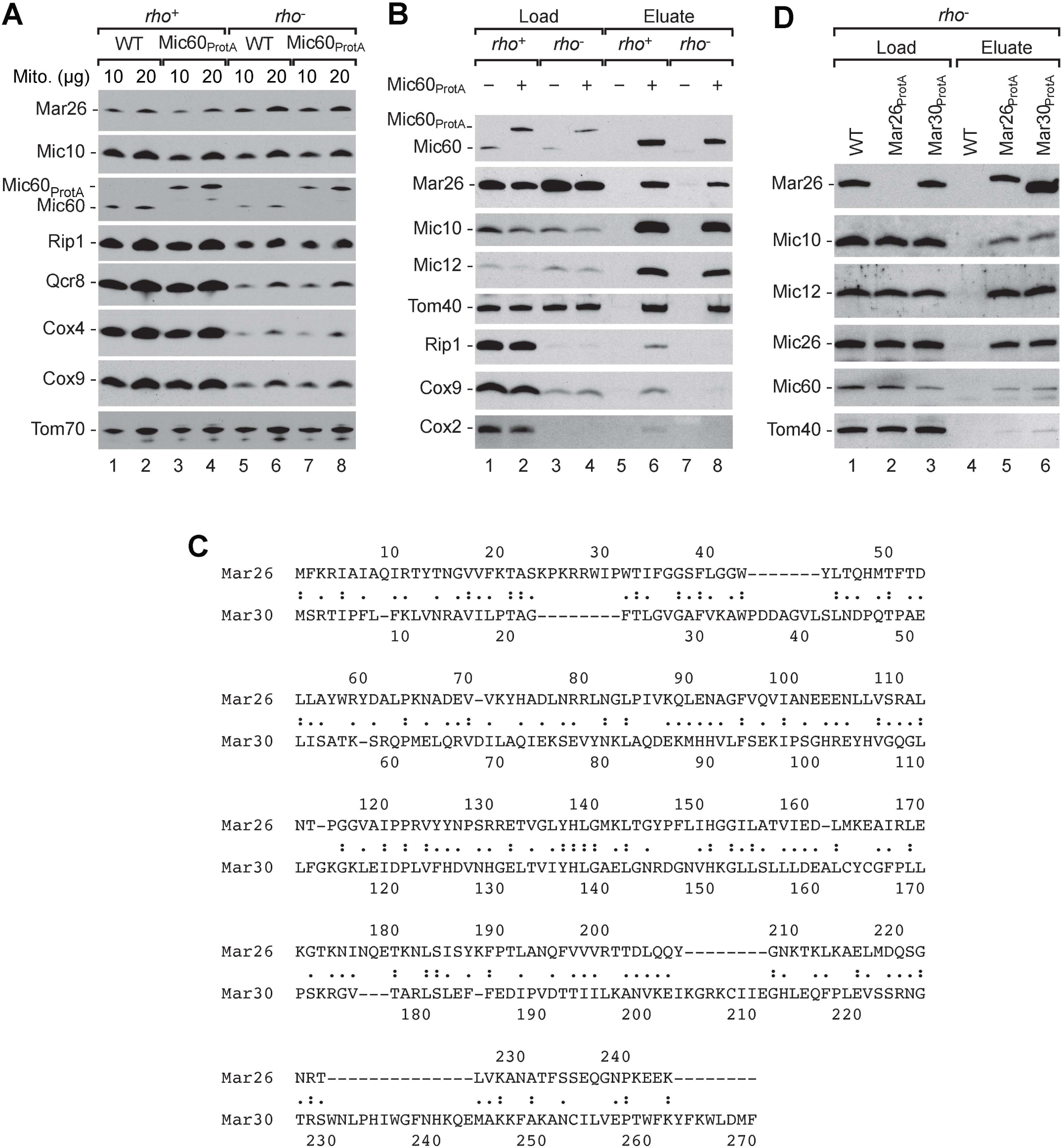
Mar26 and Mar30 are paralogs and interact with MICOS in the absence of a functional respiratory chain. (A) Steady state protein levels of *rho*+ or *rho*− mitochondria isolated from wild-type (WT) cells or cells expressing Mic60_ProtA_ were analyzed by SDS-PAGE and Western blotting with the indicated antibodies. (B) Protein complexes were purified from *rho*+ or *rho*− wild-type (WT) or Mic60_ProtA_ mitochondria by IgG affinity chromatography and analyzed by SDS-PAGE and Western blotting. Load, 1%; Eluate 100%. (C) Alignment of the amino acid sequences of Mar26 and Mar30. (D) Protein complexes were purified from wild-type (WT), Mar26_ProtA_ and Mar30_ProtA_ mitochondria isolated from *rho*− cells by IgG affinity chromatography and analyzed by SDS-PAGE and Western blotting. Load, 1%; Eluate 100%.

**Figure S4.**
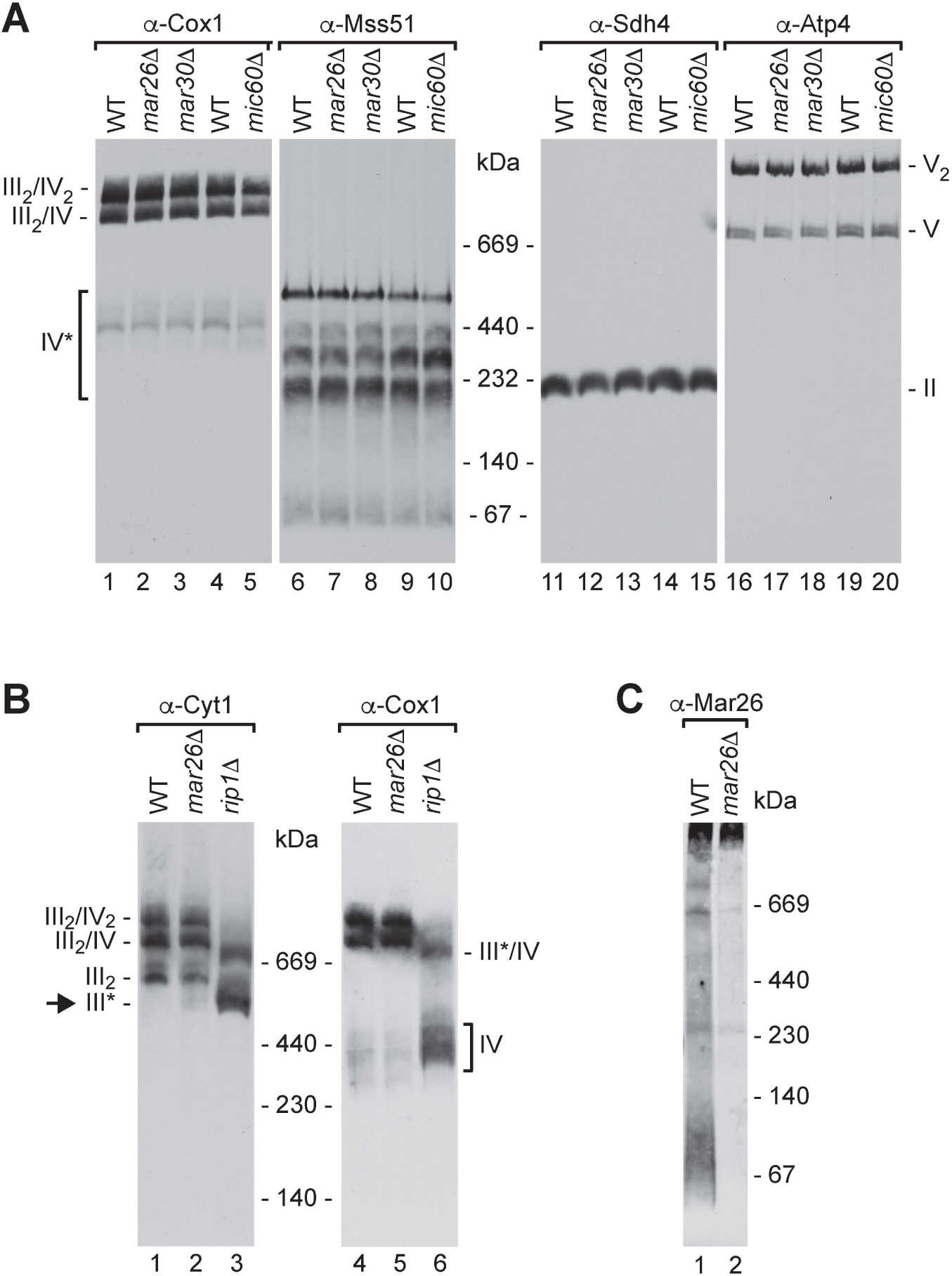
Steady state levels of mitochondrial protein complexes in *mar26*Δ mutant mitochondria. (A) WT, *mar26*Δ, and *mic60*Δ mitochondria were solubilized in digitonin buffer. Protein complexes were subsequently separated by BN-PAGE and detected by Western blotting and immunodecoration with the indicated antibodies. III_2_/IV_2_, III_2_/IV, supercomplexes of respiratory chain complexes III and IV; IV*, assembly intermediates of complex IV; II, respiratory chain complex II (SDH); V_2_, V, F_1_F_o_-ATP synthase (complex V) dimers and monomers, respectively. (B) Native protein complexes of wild-type, *mar26*Δ and *rip1*Δ mitochondria were analyzed as in (A). III_2_, complex III dimer; III*, 500 kDa late assembly intermediate of complex III; III*/IV, putative supercomplex of III* and complex IV. (C) WT and *mar26*Δ mitochondria were analyzed as in (A).

**Figure S5.**
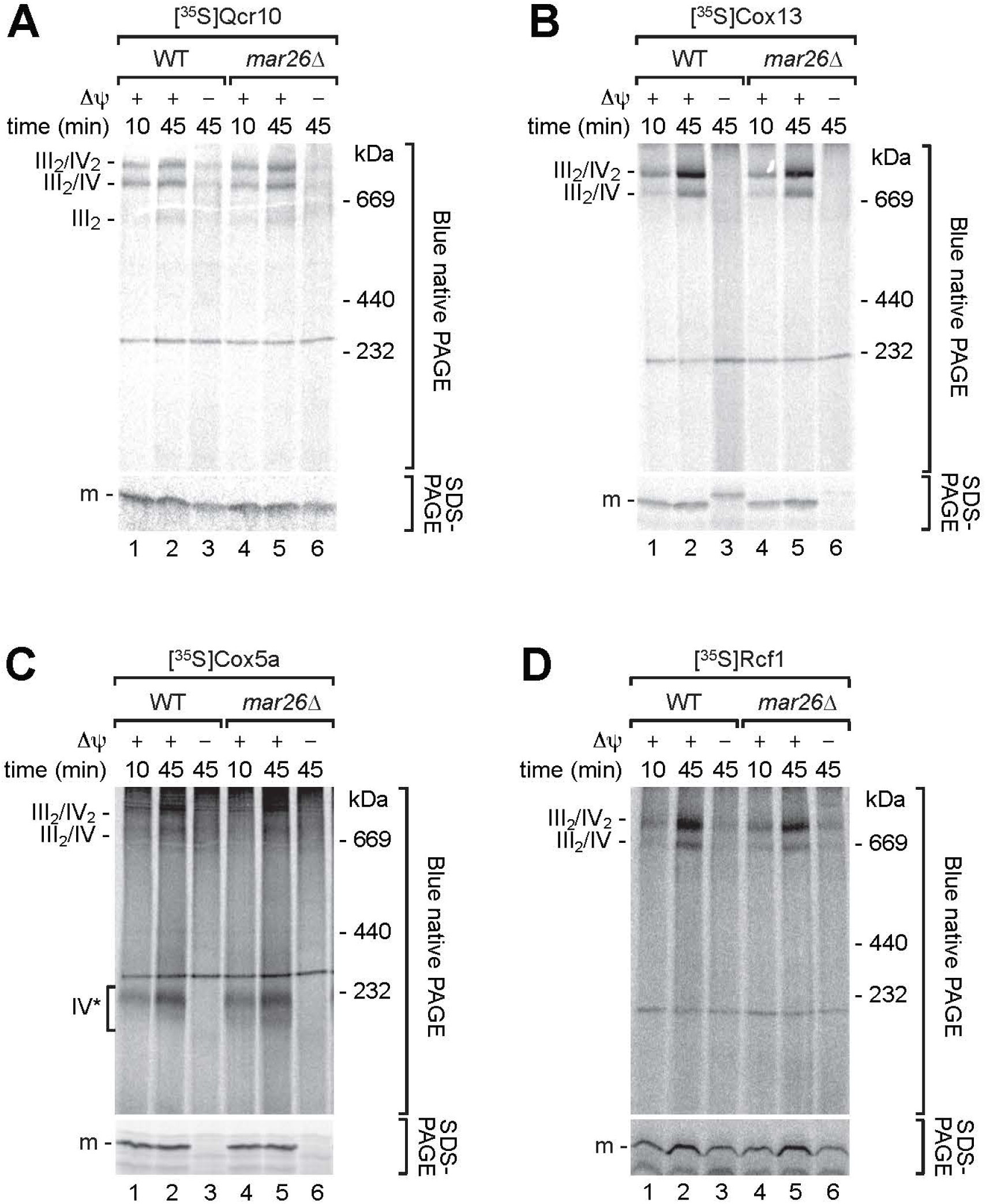
Assembly of respiratory chain complex subunits in *mar26*Δ mitochondria. Radiolabeled Qcr10 (A), Cox13 (B), Cox5a (C) or Rcf1 (D) preproteins were imported into wild-type or *mar26*Δ mitochondria. Samples were solubilized in digitonin buffer and analyzed by BN-PAGE or SDS-Page as indicated. Proteins were visualized by autoradiography. III_2_/IV_2_, III_2_/IV, III_2_, supercomplexes formed by respiratory chain complexes III and IV; IV*, assembly intermediates of complex IV; m, mature proteins; Δψ, membrane potential.

**Figures S6.**
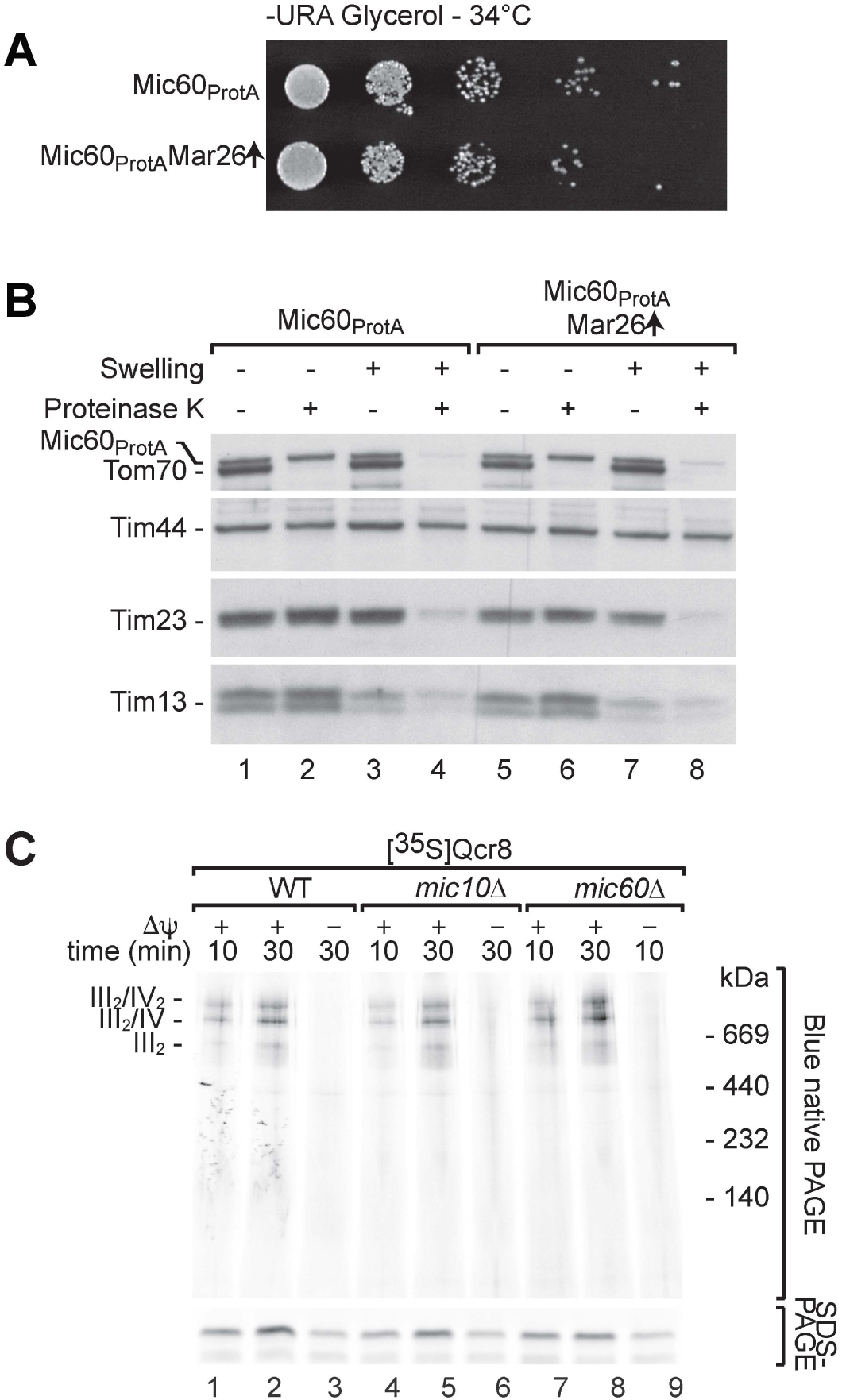
Consequences of Mar26 overexpression, and early complex III assembly in MICOS mutants. (A) Mic60_ProtA_ and Mic60_ProtA_ Mar26 overexpression (↑) yeast cells were spotted as 5-fold serial dilutions on –URA minimal medium with glycerol as carbon source and incubated at 34°C. (B) Mic60_ProtA_ and Mic60_ProtA_ Mar26 overexpression (↑) mitochondria were resuspended in hypo-osmotic buffer (Swelling) and subsequently digested with proteinase K where indicated. Samples were analyzed by SDS-PAGE and immunodecoration. (C) Radiolabeled Qcr8 was imported into mitochondria isolated from wild-type (WT), *mic10*Δ or *mic60*Δ cells. Mitochondria were subsequently solubilized in digitonin-containing buffer and analyzed by SDS-PAGE or BN-PAGE and visualized by autoradiography. III_2_/IV_2_, III_2_/IV, supercomplexes of respiratory chain complexes III and IV; Δψ, membrane potential.

**Table S1: SILAC-based quantitative MS analyses of affinity-purified Cor1, Mar26 and Mic60 complexes.**

Protein complexes were affinity-purified from differentially SILAC-labeled mitochondria using Mar26, Mic60, or Cor1 as bait (n = 3 each) and analyzed by LC/MS. Mass spectrometric raw data of all experiments were jointly processed by MaxQuant/Andromeda for protein identification and relative quantification. Light-over-heavy ratios were log2-transformed, mean log2 ratios across all three replicates of a protein complex were calculated, and the p-value for each protein was determined using a one-sided t-test. All proteins listed in this table were identified with ≥ 2 peptides (at least one of them unique) in the entire dataset, were quantified by MaxQuant in ≥ 2 replicates in at least two complexes, and exhibit a p-value ≤ 0.05 in at least one protein complex. Values shaded in grey represent log2-values of imputed ratios. For detailed information about affinity-purification, LC/MS analyses, data processing by MaxQuant, data imputation, and statistics, please refer to the methods section.

## METHODS

### Yeast growth and mitochondrial isolation

*Saccharomyces cerevisiae* strains (Table S2) used in this study are derivatives of either YPH499 (Sikorski & Hieter, 1989) (MATa *ura3-52 lys2-801 ade2-101 trp1-*Δ*63 his3-*Δ*200 leu2-*Δ*1*) or BY4741 (*MATa his3Δ0 leu2Δ0 met15Δ0 ura3Δ0*).

Yeast strains were grown either in YPG medium (1% [w/v] yeast extract, 2% [w/v] bacto-peptone, 3% [v/v] glycerol), YPEG (YPG supplemented with 3% [v/v] ethanol), YPD medium (1% [w/v] yeast extract, 2% [w/v] bacto-peptone, 2% [w/v] glucose), YPGal medium (1% [w/v] yeast extract, 2% [w/v] bacto-peptone, 2% [w/v] galactose) or selective minimal medium (0.67% [w/v] yeast nitrogen base, 0.07% [w/v] CSM amino acid mixture minus uracil, 3% [v/v] glycerol) at 30°C. Growth conditions for heavy isotope labeling of yeast cells (SILAC) and for electron microscopy analysis are described separately.

For growth tests, serial dilutions of the indicated yeast cells were spotted on agar plates containing the indicated medium and incubated at 34°C.

Crude mitochondria were isolated by differential centrifugation (Meisinger *et al*, 2006). Briefly, cells were pre-treated with DTT buffer (100 mM Tris-H_2_SO_4_ pH 9.4, 10 mM DTT) and lysed with a glass Teflon homogenizer in homogenization buffer (100 mM Tris-HCl pH 7.4, 0.6 M sorbitol, 1 mM EDTA, 0.2% bovine serum albumine, 1 mM PMSF). Cell lysates were cleared by centrifugation steps at 2000 *g* and mitochondria were pelleted at 17000 *g* and resuspended in SEM buffer (250 mM sucrose, 10 mM MOPS pH 7.2, 1 mM EDTA).

### Yeast strain construction

Strains expressing C-terminally Protein A-tagged versions of Mar26 or Mar30 were generated by PCR-based homologous recombination. The cassette used encodes the protein A moiety, a Tobacco Etch Virus (TEV) protease cleavage site and a HIS3 selection marker (Knop *et al*, 1999). Gene deletions were generated by amplifying the respective deletion cassettes from the corresponding Euroscarf strains. Cells that overexpress Mar26 were transformed with pRS426-Mar26 that contains the MAR26 wild-type open reading frame (ORF) flanked by the endogenous promotor (500 bases upstream the ORF) and terminator (300 bases downstream the ORF).

All strains were constructed by lithium acetate transformation. To generate yeast cells lacking mtDNA (*rho*-strains) the respective strains were subjected to three passages on agar plates supplemented with ethidium bromide. Primers used are listed in Table S3.

### Native analysis of protein complexes

For blue native PAGE mitochondria were solubilized in solubilization buffer (1% [w/v] digitonin, 20 mM Tris-HCl, pH 7.4, 0.1 mM EDTA, 50 mM NaCl, 10% [v/v] glycerol, and 1 mM PMSF). After solubilization non-soluble debris was removed by a clarifying spin. Loading dye was added to the supernatant and protein complexes were separated on blue native polyacrylamide gradient gels.

### Affinity purification of protein complexes

For affinity purification of native mitochondrial membrane protein complexes mitochondria expressing Protein A-tagged versions of the indicated proteins and the corresponding wild-type mitochondria were solubilized in digitonin buffer (20 mM Tris-HCl pH 7.4, 50 mM NaCl, 0.1 mM EDTA, 10% [v/v] glycerol, 1% [w/v] digitonin, 2 mM PMSF, 1 x Roche EDTA free protease inhibitor cocktail). After removing the debris by a clarifying spin (12,000 x g, 10min, 4°C) detergent extracts were incubated with IgG-coupled Sepharose beads for 2 hours. Extensive washing with washing buffer (20 mM Tris-HCl pH 7.4, 60 mM NaCl, 0.5 mM EDTA, 10% [v/v] glycerol, 0.3% [w/v] digitonin, 2 mM PMSF) removed unbound proteins from the column. Specifically bound proteins were eluted by TEV protease cleavage over night at 4°C in washing buffer. TEV elution fractions were either analyzed by SDS-PAGE or blue native PAGE.

### Protein import into isolated mitochondria

Radiolabeled precursors were generated by *in vitro* translation in the presence of [^35^S]methionine (TNT SP6 Quick Coupled or Flexi Rabbit Reticulocyte Lysate Systems) and subsequently incubated together with mitochondria (50-100 μg per import reaction) diluted in import buffer (3% [w/v] bovine serum albumin, 250 mM sucrose, 80 mM KCl, 5 mM MgCl_2_, 2 mM KH_2_PO_4_, 5 mM methionine, 10 mM MOPS-KOH pH 7.2, 4 mM ATP, 4 mM NADH, 5 mM creatine phosphate, 100 μg/ml creatine kinase) at 25°C or 30°C.(Stojanovski *et al*, 2007; Priesnitz *et al*, 2020) Import reactions were stopped by the addition of AVO mix (8 μM antimycin A, 1 μM valinomycin, 20 μM oligomycin). Mitochondria were washed with SEM buffer and analyzed by SDS-PAGE or blue native PAGE followed by autoradiography.

### Protein localization

To asses the submitochondrial localization of proteins, mitochondria were suspended in SEM buffer (250 mM sucrose, 10 mM MOPS pH 7.2, 1 mM EDTA) and subsequently diluted 1:10 in EM buffer (10 mM MOPS pH 7.2, 1 mM EDTA). After 30 min incubation on ice, proteinase K was added to a final concentration of 25 μg/ml for another 15 min. Proteinase K was inactivated by the addition of 2 mM PMSF. To determine if proteinase K digested proteins upon lysis of mitochondria, mitochondria were solubilized in SEM supplemented with 0.5 % [v/v] Triton X-100. Subsequently proteinase K was added to a final concentration of 25 μg/ml. Proteinase K was inactivated by the addition of 2 mM PMSF. Proteins were analyzed by SDS-PAGE and Western blotting.

To separate soluble and membrane-associated proteins, isolated mitochondria were incubated in 0.1 M Na_2_CO_3_ at pH 10.8 or at pH 11.5 on ice for 30 min. Subsequently membrane bound proteins were pelleted by ultracentrifugation for 30 min at 100,000 x g at 4°C. Supernatant and pellet fractions were precipitated with trichloroacetic acid and analyzed by SDS-PAGE and Western blotting.

### Stable isotope labeling with amino acids in cell culture (SILAC)

Wild-type, Mic60_ProtA_, Mar26_ProtA_ and Cor1_TAP_ yeast cells were grown in synthetic medium (0.67% [w/v] bacto-yeast nitrogen base, amino acid mix containing histidine, tryptophan, adenine, methionine, uracil, isoleucine, tyrosine, phenylalanine, leucine, valine, threonine and proline, 3% [v/v] glycerol) supplemented with 0.1 mg/ml ampicillin. Wild-type cells were labeled with 22.84 mg/l [^13^C_6_/^15^N_4_] L-arginine and 23.52 mg/l [^13^C_6_/^15^N_2_] L-lysine (Euriso-top, Gif-sur-Yvette, France) while Mic60_ProtA_, Mar26_ProtA_ and Cor1_TAP_ cells were grown in media supplemented with the corresponding [^12^C/^14^N] amino acids (18 mg/l arginine and 22.5 mg/l lysine).

### LC/MS and MS data analysis

LC/MS sample preparation and analysis of affinity-purified MICOS complexes has been described before (Malsburg *et al*, 2011), and Cor1 and Mar26 complexes were processed essentially following the same protocol. In brief, differentially SILAC-labeled protein complexes were acetone-precipitated, proteins resuspended in 60% (v/v) methanol/20 mM NH_4_HCO_3_, and digested with trypsin. Peptide mixtures of three independent replicates per protein complex were analyzed on an LTQ-Orbitrap XL (Thermo Scientific, Bremen, Germany), which was directly coupled to an UltiMate^TM^ 3000 HPLC or RSLCnano system (Thermo Scientific, Dreieich, Germany). Peptides were separated on C18 reversed-phase nano LC columns applying a 135- to 150-min linear gradient of increasing acetonitrile concentration [4 - 34% (v/v) in 0.1% (v/v) formic acid] at a flow rate of 300 nl/min (UltiMate^TM^ 3000 HPLC) or 250 nl/min (UltiMate^TM^ 3000 RSLCnano). MS survey scans ranging from *m/z* 300 to 2,000 (Cor1) or 370 to 1,700 (Mic60 and Mar26) were acquired in the orbitrap at a resolution of 60,000. Peptide ions with a charge of ≥ +2 were selected for fragmentation by collision-induced dissociation in the linear ion trap using a top6 method and applying a dynamic exclusion time of 45 sec.

Mass spectrometric raw data of Cor1, Mic60, and Mar26 complexes were processed together using MaxQuant (version 1.4.1.2) and its search engine Andromeda (Cox & Mann, 2008; Cox *et al*, 2011). For protein identification, MS spectra were correlated with the *Saccharomyces* Genome Database (version of 02/03/2011). Heavy labels were set to Arg6/Lys6 for data derived from Cor1 complexes and Arg10/Lys8 for Mic60 and Mar26 data. The database search was performed with tryptic specificity; a maximum of two missed cleavages; acetylation of protein N-termini and oxidation of methionine as variable modifications; precursor and fragment ion mass tolerances of 4.5 ppm and 0.5 Da, respectively; a false discovery rate of 1% on peptide and protein level; and at least one unique peptide with a minimum length of 7 amino acids. Relative protein quantification was based on unique peptides and at least one SILAC peptide pair (i.e. ratio count). The data analysis is described in detail in the section Quantification and statistical analysis.

### Electron microscopy

For electron microscopy cells were inoculated in minimal medium(van Dijken *et al*, 1976) containing 2% (v/v) L-lactate (pH 5.0) and 0.1% glucose. After 24 h at 30°C and 200 rpm cultures were diluted to an OD_600_ of 0.1 in minimal medium containing only 2% (v/v) L-lactate (pH 5.0) as carbon source. After 16 h cells were harvested, washed with 0.1 M sodium cacodylate buffer pH 7.2 and subsequently fixed with 3% (v/v) glutaraldehyde in 0.1 M sodium cacodylate buffer (pH 7.2) for 1 h on ice. For diaminobenzidine (DAB) staining fixed cells were incubated for 45 min in 0.1 M Tris buffer (pH 7.5) with 2 mg/ml 3:3’-diaminobenzidine and 0.06% (v/v) H_2_O_2_ at 30°C under constant aeration. For post-fixation cells were incubated in 1.5% (w/v) KMnO_4_ for 20 min at room temperature, followed by incubation in 0.5% (w/v) uranyl acetate overnight and embedding in Epon 812. For statistical analysis 100 random cell sections were imaged and the number of crista junctions in each section was counted.

## STATISTICAL ANALYSIS

For mass spectrometry analysis (Figure 3A, Table S1), MaxQuant results were further processed using an in-house developed data analysis pipeline programmed in R. In case no SILAC ratio was reported by MaxQuant, the missing value was estimated and replaced based on the distribution of proteins with the lowest 5% of MS intensity (Arg0/Lys0 label) of all proteins identified in a replicate. Following data imputation, ratios (light-over-heavy) were log_2_-transformed, the mean log_2_ ratios across all three replicates of a protein complex were calculated, and a one-sided t-test was performed to determine the p-value for each protein. Data about MaxQuant protein identification and quantification are provided in Table S1.

## DATA AVAILABILITY

This study includes no data deposited in external repositories.

## ACKNOWLEDGEMENTS

We thank Florian Wollweber, Drs. Susanne Horvath, Nikolaus Pfanner and Nils Wiedemann for discussion, and Thorina Boenke and Anita Kram for materials and technical assistance, respectively. Work included in this study has also been performed in partial fulfillment of the requirements for the doctoral theses of R.M.Z., C.S.M. and C.D.P. at the University of Freiburg. This project was supported by the Deutsche Forschungsgemeinschaft (DFG, German Research Foundation) FOR 2848 – project ID 401510699 (to H.R. and M.v.d.L.), SFB 894 - project ID 157660137 and SFB 746 - project ID 26068018 (to M.v.d.L.), the Excellence Initiative of the German Federal and State Governments (BIOSS – EXC-294 to M.v.d.L. and B.W., and CIBSS – EXC-2189 – project ID 390939984 to H.R.) and the European Research Council (ERC) Consolidator Grant 648235 (to B.W.).

## AUTHOR CONTRIBUTIONS

R.M.Z., M.B., H.R., and M.v.d.L. conceptualized and guided the project, R.M.Z., L.C.-T., M.B., K.v.d.M., C.S.M., I.P., R.K., and S.O. performed the experiments and analyzed the data together with C.D.P., I.v.d.K., B.W., H.R. and M.v.d.L. Preparation of figures and tables was done by R.M.Z., M.B., C.D.P., S.O., H.R. and M.v.d.L. R.M.Z., H.R. and M.v.d.L. wrote the manuscript with assistance from all other authors.

---

**Table S2.** *Saccharomyces* cerevisiae strains used in this study.

| STRAIN | SOURCE | IDENTIFIER |
| --- | --- | --- |
| WT (YPH499 – MATa; *ade2-101*; *his3-Δ200*; *leu2-Δ1*; *ura3-52*; *trp1-Δ63*; *lys2-801*) | Sikorski and Hieter, 1989<sup>103</sup> | 1501 |
| WT (BY4741 – MATa; *his3-Δ1*; *leu2-Δ0*; *met15Δ0*; *ura3-Δ0*) | Euroscarf | 1354 |
| Mic60<sub>ProtA</sub> (YPH499 *mic60::MIC60<sub>ProA</sub>-HIS3MX6*) | von der Malsburg et al., 2011<sup>30</sup> | 2035 |
| Mic12<sub>ProtA</sub> (YPH499 *mic12::MIC12<sub>ProA</sub>-HIS3MX6*) | von der Malsburg et al., 2011<sup>30</sup> | 3083 |
| Mic60<sub>ProtA</sub> *mic10Δ* (YPH499 *mic60::MIC60<sub>ProA</sub>-HIS3MX6*; *mic10::kanMX4*) | von der Malsburg et al., 2011<sup>30</sup> | 3049 |
| Mic12<sub>ProtA</sub> *mic10Δ* (YPH499 *mic12::MIC12<sub>ProA</sub>-HIS3MX6*; *mic10::kanMX4*) | This study | 4042 |
| Mic12<sub>ProtA</sub> *mic60Δ* (YPH499 *mic12::MIC12<sub>ProA</sub>-HIS3MX6*; *mic60::kanMX4*) | This study | 3093 |
| Mic60<sub>ProtA</sub> *mar26Δ* (YPH499 *mic60::MIC60<sub>ProA</sub>-HIS3MX6*; *mar26::kanMX4*) | This study | 3044 |
| Mar26<sub>ProtA</sub> (YPH499 *mar26::MAR26<sub>ProA</sub>-HIS3MX6*) | This study | 4047 |
| Mar30<sub>ProtA</sub> (YPH499 *mar30::MAR30<sub>ProA</sub>-HIS3MX6*; *arg4Δ*) | This study | 3579 |
| Cor1<sub>TAP</sub> (YPH499 *cor1::COR1<sub>TAP</sub>-HIS3MX6*) | van der Laan et al., 2006<sup>94</sup> | 2510 |
| WT rho- (YPH499 rho-) | This study | 3777 |
| Mic60<sub>ProtA</sub> rho- (YPH499 *mic60::MIC60<sub>ProA</sub>-HIS3MX6*; rho-) | This study | 3778 |
| *mar26Δ* (YPH499 *mar26::kanMX4*) | This study | 3043 |
| *mar30Δ* (YPH499 *mar30::kanMX4*) | This study | 3575 |
| *mic60Δ* (YPH499 *mic60::kanMX4*) | von der Malsburg et al., 2011<sup>30</sup> | 3092 |
| *mic10Δ* (YPH499 *mic10::kanMX4*) | von der Malsburg et al., 2011<sup>30</sup> | 3047 |
| WT pRS426 (YPH499 pRS426) | Bohnert et al., 2015<sup>46</sup> | 3243 |
| Mic60<sub>ProtA</sub> pRS426 (YPH499 *mic60::MIC60<sub>ProA</sub>-HIS3MX6*; pRS426) | Bohnert et al., 2015<sup>46</sup> | 3244 |
| Mic60<sub>ProtA</sub> Mar26 up (YPH499 *mic60::MIC60<sub>ProA</sub>-HIS3MX6*; pRS426-MAR26) | This study | 3587 |
| Mar26<sub>ProtA</sub> rho- (YPH499 *mar26::MAR26<sub>ProA</sub>-HIS3MX6*; rho-) | This study | 3782 |
| Mar30<sub>ProtA</sub> rho- (YPH499 ; *mar30::MAR30<sub>ProA</sub>-HIS3MX6*; *arg4Δ*; rho-) | This study | 3783 |
| *mar26Δ* (BY4741 *mar26::kanMX4*) | Euroscarf | 2835 |
| *mar30Δ* (BY4741 *mar30::kanMX4*) | Euroscarf | 2833 |
| *rip1Δ* (BY4741 *rip1::kanMX4*) | Euroscarf | 1932 |

**Table S3.**
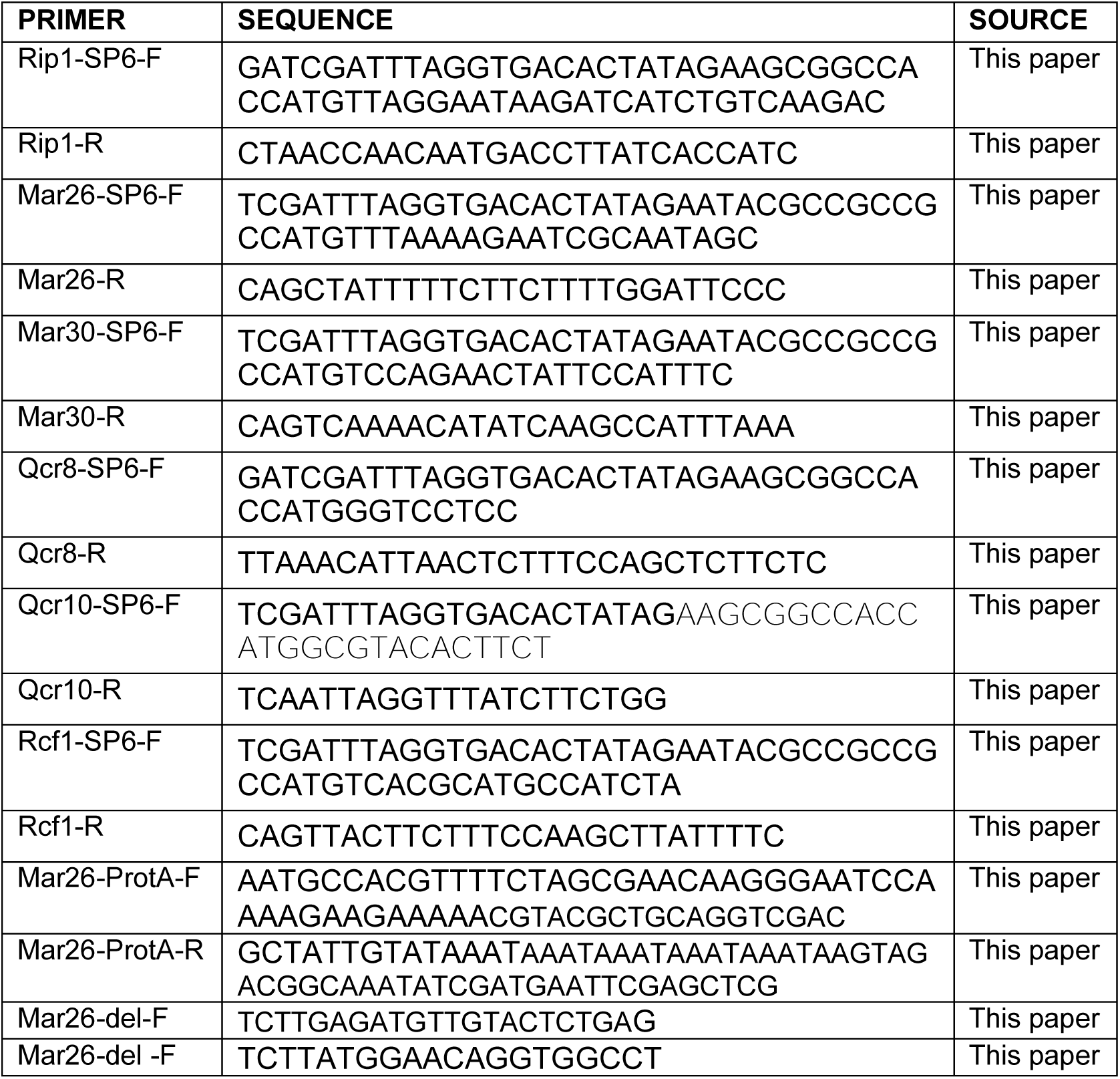
Oligonucleotides used in this study.

