## Supplementary material for "Coordination of cytochrome *bc*_1_ complex assembly at MICOS": Table S1

**Supplemental Table SX: Results of SILAC-based quantitative MS analyses of affinity-purified Cor1, Mar26 and Mic60 complexes.** Protein complexes were affinity-purified from differentially SILAC-labeled mitochondria using Mar26, Mic60, or Cor1 as bait (n = 3 each) and analyzed by LC/MS. Mass spectrometric raw data of all experiments were jointly processed by MaxQuant/Andromeda (Cox and Mann, Nat. Biotechnol. 26, 2008; Cox et al., J. Proteome Res. 10, 2011) for protein identification and relative quantification. Light-over-heavy ratios were $log_2$-transformed, mean $log_2$ ratios across all three replicates of a protein complex were calculated, and the p-value for each protein was determined using a one-sided t-test. All proteins listed in this table were identified with ≥ 2 peptides (at least one of them unique) in the entire dataset, were quantified by MaxQuant in ≥ 2 replicates in at least two complexes, and exhibit a p-value ≤ 0.05 in at least one protein complex. Values shaded in grey represent $log_2$-values of imputed ratios. For detailed information about affinity-purification, LC/MS analyses, data processing by MaxQuant, data imputation, and statistics, please refer to the methods section.

| | | | | | | | | | | | Mar26 - Replicate 1 | | | | | | Mar26 - Replicate 2 | | | | | | | |
|---|---|---|---|---|---|---|---|---|---|---|---|---|---|---|---|---|---|---|---|---|---|---|---|---|
| Systematic Names | Gene Names | Molecular Weight (kDa) | Sequence Coverage (%) | Posterior Error Probability | Mean log2 Ratio Mar26/WT | t-Test p-Value Mar26 | Mean log2 Ratio Mic60/WT | t-Test p-Value Mic60 | Mean log2 Ratio Cor1/WT | t-Test p-Value Cor1 | Peptides | Unique Peptides | Ratio Mar26/WT | log2 Ratio Mar26/WT | Ratio Variability [%] | Ratio Count | Peptides | Unique Peptides | Ratio Mar26/WT | log2 Ratio Mar26/WT | Ratio Variability [%] | Ratio Count | Peptides | Unique Peptides |
| YER182W | MAR26 | 27,70 | 67,60 | 7,28E-226 | 6,594 | 0,001020 | 4,076 | 0,002524 | 3,807 | 0,000573 | 18 | 18 | 100,866 | 6,656 | 142,18 | 20 | 14 | 14 | 66,221 | 6,049 | 154,91 | 13 | 18 | 18 |
| YBL045C | COR1 | 50,23 | 82,70 | 0 | 6,227 | 0,001028 | 4,099 | 0,002879 | 4,009 | 0,000676 | 24 | 24 | 103,968 | 6,700 | 112,62 | 23 | 16 | 16 | 52,787 | 5,722 | 59,30 | 18 | 25 | 25 |
| YPR191W | QCR2 | 40,48 | 95,70 | 0 | 5,932 | 0,001307 | 4,485 | 0,012663 | 4,223 | 0,000988 | 25 | 25 | 86,972 | 6,442 | 128,73 | 26 | 21 | 21 | 41,964 | 5,391 | 93,14 | 27 | 25 | 25 |
| YJL166W | QCR8 | 10,97 | 89,40 | 6,34E-51 | 5,666 | 0,001581 | 4,032 | 0,005861 | 5,876 | 0,001380 | 4 | 4 | 52,200 | 5,706 | 86,19 | 2 | 4 | 4 | 34,159 | 5,094 | 210,68 | 4 | 5 | 5 |
| YMR145C | NDE1 | 62,77 | 73,80 | 0 | 5,644 | 0,002300 | 4,883 | 0,000183 | 3,229 | 0,000450 | 29 | 28 | 73,405 | 6,198 | 82,50 | 34 | 23 | 23 | 29,979 | 4,906 | 139,31 | 26 | 30 | 29 |
| YCL057C-A | MIC10 | 10,41 | 88,70 | 1,75E-114 | 5,641 | 0,008024 | 4,879 | 0,001383 | 4,589 | 0,000449 | 5 | 5 | 88,176 | 6,462 | 66,96 | 5 | 5 | 5 | 18,363 | 4,199 | 71,22 | 5 | 6 | 6 |
| YLR038C | COX12 | 9,79 | 74,70 | 3,83E-66 | 5,542 | 0,001584 | 5,488 | 0,001111 | 7,435 | 0,005766 | 1 | 1 | 51,010 | 5,673 | 36,51 | 3 | 1 | 1 | 30,834 | 4,946 | 53,04 | 3 | 1 | 1 |
| YBL095W | MAR30 | 30,55 | 73,00 | 1,78E-113 | 5,533 | 0,002344 | 3,634 | 0,000504 | 1,794 | 0,001805 | 10 | 10 | 70,756 | 6,145 | 124,87 | 11 | 7 | 7 | 49,075 | 5,617 | 135,42 | 10 | 15 | 15 |
| YOR065W | CYT1 | 34,05 | 43,00 | 0 | 5,523 | 0,001263 | 4,770 | 0,008920 | 6,389 | 0,000003 | 7 | 7 | 49,865 | 5,640 | 74,31 | 12 | 6 | 6 | 61,226 | 5,936 | 39,14 | 6 | 7 | 7 |
| YKR016W | MIC60 | 61,08 | 78,30 | 0 | 5,271 | 0,003146 | 5,502 | 0,001509 | 2,873 | 0,001234 | 26 | 26 | 46,668 | 5,544 | 120,34 | 23 | 20 | 20 | 21,802 | 4,446 | 51,19 | 24 | 24 | 24 |
| YGL187C | COX4 | 17,14 | 73,50 | 2,56E-265 | 5,148 | 0,000983 | 3,938 | 0,014479 | 6,608 | 0,000347 | 7 | 7 | 48,636 | 5,604 | 75,28 | 8 | 6 | 6 | 30,774 | 4,944 | 74,03 | 7 | 7 | 7 |
| YGL191W | COX13 | 15,02 | 80,60 | 1,64E-134 | 5,070 | 0,000271 | 3,228 | 0,003150 | 5,879 | 0,000774 | 8 | 8 | 33,843 | 5,081 | 118,38 | 7 | 1 | 1 | 38,568 | 5,269 | n. def. | 1 | 5 | 5 |
| YBL030C;YBR085W | PET9; AAC3 | 34,43 | 75,80 | 0 | 5,056 | 0,001038 | 4,158 | 0,003443 | 5,508 | 0,000885 | 16 | 15 | 38,782 | 5,277 | 122,73 | 22 | 14 | 13 | 24,166 | 4,595 | 53,50 | 19 | 18 | 17 |
| YHR051W | COX6 | 17,34 | 58,80 | 4,49E-236 | 4,828 | 0,006535 | 3,464 | 0,001858 | 8,164 | 0,000771 | 4 | 4 | 61,478 | 5,942 | 76,51 | 4 | 4 | 4 | 19,668 | 4,298 | 71,86 | 5 | 7 | 7 |
| YPL004C | LSP1 | 38,07 | 58,40 | 1,04E-187 | 4,809 | 0,001943 | 0,976 | 0,218101 | -0,418 | 0,992252 | 13 | 11 | 40,336 | 5,334 | 59,25 | 11 | 11 | 9 | 19,599 | 4,293 | 83,94 | 9 | 13 | 11 |
| YFR011C | MIC19 | 18,86 | 64,10 | 1,97E-222 | 4,738 | 0,001325 | 5,530 | 0,000726 | 3,101 | 0,007722 | 9 | 9 | 37,238 | 5,219 | 67,43 | 7 | 7 | 7 | 21,403 | 4,420 | 87,39 | 7 | 10 | 10 |
| Q0250 | COX2 | 28,57 | 44,20 | 7,15E-118 | 4,716 | 0,011539 | 3,673 | 0,003517 | 6,184 | 0,004547 | 4 | 4 | 65,108 | 6,025 | 122,02 | 6 | 4 | 4 | 11,352 | 3,505 | 132,52 | 4 | 4 | 4 |
| YDL174C | DLD1 | 65,29 | 60,30 | 1,26E-246 | 4,690 | 0,000874 | 3,722 | 0,010523 | 2,269 | 0,000899 | 21 | 21 | 32,139 | 5,006 | 144,27 | 23 | 15 | 15 | 26,606 | 4,734 | 138,88 | 16 | 16 | 16 |
| YJR077C | MIR1 | 32,81 | 73,60 | 1,60E-262 | 4,661 | 0,004472 | 4,215 | 0,001401 | 1,290 | 0,054428 | 16 | 16 | 41,973 | 5,391 | 101,59 | 17 | 12 | 12 | 26,588 | 4,733 | 93,17 | 13 | 12 | 12 |
| YDR529C | QCR7 | 14,57 | 97,60 | 2,69E-299 | 4,579 | 0,005456 | 4,296 | 0,002924 | 6,770 | 0,001254 | 5 | 5 | 13,782 | 3,785 | 243,84 | 10 | 8 | 8 | 43,723 | 5,450 | 137,25 | 9 | 9 | 9 |
| YEL024W | RIP1 | 23,37 | 63,30 | 1,88E-293 | 4,542 | 0,000778 | 4,843 | 0,003939 | 4,693 | 0,002578 | 12 | 12 | 25,561 | 4,676 | 118,35 | 13 | 13 | 13 | 18,207 | 4,186 | 74,15 | 16 | 12 | 12 |
| YGR086C | PIL1 | 38,35 | 69,60 | 1,80E-176 | 4,488 | 0,002259 | 0,886 | 0,200101 | -0,467 | 0,995816 | 9 | 7 | 27,131 | 4,762 | 105,27 | 9 | 10 | 8 | 14,763 | 3,884 | 51,53 | 9 | 12 | 10 |
| YHR001W-A | QCR10 | 8,59 | 55,80 | 1,55E-50 | 4,435 | 0,000396 | 4,605 | 0,005074 | 1,612 | 0,125158 | 4 | 4 | 24,433 | 4,611 | 174,57 | 4 | 2 | 2 | 22,656 | 4,502 | 86,12 | 2 | 3 | 3 |
| YDL198C | GGC1 | 33,22 | 51,70 | 5,24E-90 | 4,404 | 0,000955 | 3,421 | 0,000113 | 2,421 | 0,058027 | 9 | 9 | 20,523 | 4,359 | 80,02 | 7 | 4 | 4 | 27,044 | 4,757 | 48,94 | 3 | 8 | 8 |
| YOL027C | MDM38 | 65,00 | 56,00 | 0 | 4,402 | 0,004848 | 1,618 | 0,092329 | 3,102 | 0,000084 | 22 | 22 | 28,604 | 4,838 | 93,51 | 24 | 12 | 12 | 11,543 | 3,529 | 52,89 | 14 | 14 | 14 |
| YER141W | COX15 | 54,66 | 24,90 | 3,56E-51 | 4,385 | 0,000516 | 4,000 | 0,001455 | 1,615 | 0,169885 | 8 | 8 | 19,727 | 4,302 | 200,53 | 6 | 3 | 3 | 18,288 | 4,193 | 86,45 | 3 | 3 | 3 |
| YNL100W | MIC27 | 26,96 | 79,90 | 1,40E-199 | 4,302 | 0,004319 | 5,121 | 0,002619 | 1,693 | 0,075017 | 9 | 9 | 33,821 | 5,080 | 203,00 | 8 | 5 | 5 | 13,309 | 3,734 | 35,43 | 5 | 8 | 8 |
| YDL085W | NDE2 | 61,66 | 67,00 | 0 | 4,253 | 0,002658 | 3,784 | 0,002929 | 2,963 | 0,002887 | 22 | 21 | 20,963 | 4,390 | 146,87 | 29 | 15 | 15 | 26,177 | 4,710 | 176,19 | 14 | 20 | 19 |
| YMR203W | TOM40 | 42,04 | 56,80 | 9,99E-144 | 4,246 | 0,002969 | 4,198 | 0,038238 | 2,194 | 0,206864 | 5 | 5 | 18,547 | 4,213 | 73,24 | 5 | 4 | 4 | 12,940 | 3,694 | 90,08 | 4 | 5 | 5 |
| YDR119W-A | COX26 | 7,45 | 30,30 | 1,04E-15 | 4,218 | 0,003256 | 5,222 | 0,011131 | 7,232 | 0,002902 | 1 | 1 | 25,133 | 4,652 | n. def. | 1 | 1 | 1 | 22,014 | 4,460 | n. def. | 1 | 1 | 1 |
| YBR262C | MIC12 | 12,39 | 71,70 | 7,68E-147 | 4,150 | 0,021328 | 4,087 | 0,000004 | -0,173 | 0,573018 | 8 | 8 | 32,289 | 5,013 | 122,99 | 9 | 8 | 8 | 5,200 | 2,379 | 99,60 | 8 | 7 | 7 |
| YGR174C | CBP4 | 20,22 | 58,80 | 1,15E-94 | 4,149 | 0,009883 | 4,704 | 0,003359 | 4,301 | 0,000628 | 5 | 5 | 40,235 | 5,330 | 68,88 | 4 | 2 | 2 | 12,404 | 3,633 | 63,07 | 2 | 5 | 5 |
| YJR095W | SFC1 | 35,34 | 56,20 | 5,01E-108 | 4,145 | 0,006559 | 3,587 | 0,004673 | 1,625 | 0,001842 | 12 | 12 | 22,950 | 4,520 | 76,60 | 10 | 5 | 5 | 9,147 | 3,193 | 52,52 | 6 | 7 | 7 |
| YGR235C | MIC26 | 26,91 | 58,40 | 1,45E-148 | 4,090 | 0,007622 | 3,375 | 0,000216 | 1,583 | 0,023028 | 9 | 9 | 27,783 | 4,796 | 116,25 | 11 | 4 | 4 | 20,759 | 4,376 | 42,92 | 4 | 6 | 6 |
| YPR125W | YLH47 | 52,17 | 35,90 | 2,26E-49 | 4,056 | 0,008591 | 2,278 | 0,094927 | 2,311 | 0,000577 | 9 | 9 | 24,356 | 4,606 | 80,59 | 7 | 5 | 5 | 7,881 | 2,978 | 39,79 | 5 | 5 | 5 |
| YPL173W | MRPL40 | 33,75 | 45,10 | 3,00E-80 | 3,962 | 0,006214 | 0,094 | 0,395959 | 2,740 | 0,103017 | 9 | 9 | 18,531 | 4,212 | 55,38 | 10 | 6 | 6 | 8,547 | 3,095 | 21,39 | 7 | 6 | 6 |
| YMR056C | AAC1 | 34,12 | 44,00 | 7,03E-95 | 3,941 | 0,000559 | 4,486 | 0,000328 | 5,131 | 0,028385 | 4 | 3 | 16,915 | 4,080 | 120,28 | 3 | 7 | 6 | 16,744 | 4,066 | 108,32 | 5 | 4 | 3 |
| YLL041C | SDH2 | 30,23 | 35,70 | 2,44E-73 | 3,925 | 0,005677 | 3,799 | 0,013163 | 1,977 | 0,042673 | 4 | 4 | 27,244 | 4,768 | 24,76 | 4 | 3 | 3 | 11,506 | 3,524 | 36,03 | 3 | 4 | 4 |
| YBR024W | SCO2 | 34,89 | 44,20 | 1,95E-36 | 3,776 | 0,000422 | 2,491 | 0,011469 | 1,857 | 0,047768 | 6 | 6 | 13,137 | 3,716 | 63,55 | 4 | 4 | 4 | 12,324 | 3,623 | 62,59 | 4 | 4 | 4 |
| YDR316W | OMS1 | 55,59 | 42,00 | 4,08E-90 | 3,775 | 0,001711 | 1,772 | 0,037449 | 2,972 | 0,000344 | 11 | 11 | 17,628 | 4,140 | 111,64 | 8 | 4 | 4 | 10,376 | 3,375 | 17,55 | 2 | 6 | 6 |
| YPL118W | MRP51 | 39,45 | 45,30 | 5,52E-71 | 3,742 | 0,016399 | 2,838 | 0,000706 | 5,252 | 0,000034 | 11 | 11 | 33,668 | 5,073 | 74,85 | 12 | 6 | 6 | 6,638 | 2,731 | 59,95 | 7 | 8 | 8 |
| YDR041W | RSM10 | 23,42 | 52,70 | 1,90E-84 | 3,742 | 0,008138 | 2,420 | 0,097793 | 0,905 | 0,152884 | 6 | 6 | 25,907 | 4,695 | 145,22 | 7 | 4 | 4 | 8,744 | 3,128 | 34,30 | 5 | 5 | 5 |
| YGR076C | MRPL25 | 18,59 | 25,50 | 7,18E-14 | 3,722 | 0,007557 | 1,767 | 0,128248 | 1,290 | 0,163722 | 4 | 4 | 21,929 | 4,455 | 54,66 | 5 | 2 | 2 | 7,289 | 2,866 | 10,24 | 3 | 2 | 2 |
| YGR231C | PHB2 | 34,41 | 49,00 | 1,54E-110 | 3,699 | 0,009923 | 1,475 | 0,136354 | 0,538 | 0,173946 | 10 | 10 | 14,469 | 3,855 | 41,37 | 14 | 8 | 8 | 6,566 | 2,715 | 14,94 | 11 | 6 | 6 |
| YPL098C | MGR2 | 12,25 | 34,50 | 9,77E-27 | 3,680 | 0,005920 | 1,802 | 0,248817 | 3,619 | 0,015354 | 2 | 2 | 20,024 | 4,324 | 136,46 | 2 | 3 | 3 | 13,743 | 3,781 | 54,23 | 3 | 2 | 2 |
| YGR165W | MRPS35 | 39,58 | 30,10 | 4,35E-41 | 3,653 | 0,014152 | 2,050 | 0,016037 | 0,128 | 0,174855 | 6 | 6 | 26,615 | 4,734 | 190,17 | 6 | 4 | 4 | 12,703 | 3,667 | 168,21 | 5 | 2 | 2 |
| YPR010C-A | MIN8 | 7,90 | 37,50 | 1,96E-48 | 3,647 | 0,014096 | 4,242 | 0,020608 | 1,468 | 0,014360 | 1 | 1 | 29,682 | 4,891 | n. def. | 1 | 2 | 2 | 7,509 | 2,909 | 148,75 | 2 | 2 | 2 |
| YIL087C | AIM19 | 16,92 | 52,90 | 3,15E-78 | 3,610 | 0,005566 | 2,873 | 0,029981 | 4,550 | 0,000695 | 5 | 5 | 14,416 | 3,850 | 99,16 | 7 | 3 | 3 | 17,425 | 4,123 | 57,52 | 3 | 5 | 5 |
| YBR255C-A | RCF3 | 13,74 | 28,30 | 1,87E-42 | 3,587 | 0,014265 | 1,769 | 0,055799 | 2,750 | 0,044656 | 2 | 2 | 26,765 | 4,742 | n. def. | 1 | 0 | 0 | NA | 3,396 | n. def. | 0 | 2 | 2 |
| YMR241W | YHM2 | 34,18 | 43,30 | 2,53E-124 | 3,585 | 0,005610 | 3,862 | 0,010524 | 3,423 | 0,002967 | 5 | 5 | 11,236 | 3,490 | 215,47 | 5 | 5 | 5 | 19,568 | 4,290 | 67,62 | 7 | 6 | 6 |
| YPL063W | TIM50 | 55,10 | 49,60 | 5,68E-120 | 3,583 | 0,017001 | 2,192 | 0,039270 | 3,169 | 0,012668 | 9 | 9 | 7,693 | 2,944 | 85,56 | 9 | 6 | 6 | 7,297 | 2,867 | 40,35 | 5 | 2 | 2 |
| YNL131W | TOM22 | 16,79 | 47,40 | 1,66E-104 | 3,569 | 0,034903 | 4,406 | 0,005000 | 2,986 | 0,027403 | 2 | 2 | 27,657 | 4,790 | 77,00 | 3 | 2 | 2 | 19,984 | 4,321 | 36,24 | 2 | 2 | 2 |
| YGR132C | PHB1 | 31,43 | 76,30 | 2,12E-196 | 3,566 | 0,009853 | 1,318 | 0,144476 | 1,228 | 0,001432 | 12 | 12 | 13,981 | 3,805 | 29,24 | 18 | 11 | 11 | 6,025 | 2,591 | 21,96 | 17 | 10 | 10 |
| YJR085C | TMH11 | 11,25 | 63,80 | 3,41E-63 | 3,543 | 0,010105 | 2,902 | 0,012553 | 3,503 | 0,008256 | 4 | 4 | 10,440 | 3,384 | 72,28 | 5 | 4 | 4 | 22,596 | 4,498 | 61,15 | 3 | 2 | 2 |
| YLR348C | DIC1 | 32,99 | 41,90 | 3,62E-56 | 3,492 | 0,003388 | 3,489 | 0,009447 | 2,350 | 0,064712 | 3 | 3 | 12,736 | 3,671 | 29,86 | 3 | 3 | 3 | 14,698 | 3,878 | n. def. | 1 | 3 | 3 |
| YGR183C | QCR9 | 7,48 | 42,40 | 4,81E-25 | 3,488 | 0,019205 | 1,417 | 0,114756 | 5,812 | 0,007586 | 1 | 1 | 23,290 | 4,542 | n. def. | 1 | 0 | 0 | NA | 2,152 | n. def. | 0 | 1 | 1 |
| YNR017W | TIM23 | 23,24 | 51,40 | 3,49E-54 | 3,471 | 0,006786 | 2,989 | 0,015582 | 0,639 | 0,162928 | 4 | 4 | 6,305 | 2,656 | 85,26 | 3 | 3 | 3 | 14,106 | 3,818 | 3,05 | 2 | 3 | 3 |
| YMR110C | HFD1 | 59,98 | 36,50 | 2,55E-125 | 3,451 | 0,005165 | 1,420 | 0,066584 | -0,211 | 0,996089 | 12 | 12 | 17,532 | 4,132 | 80,60 | 17 | 8 | 8 | 9,685 | 3,276 | 85,06 | 8 | 7 | 7 |
| YML120C | NDI1 | 57,25 | 81,50 | 0 | 3,450 | 0,002956 | 2,114 | 0,063088 | 5,440 | 0,000318 | 19 | 19 | 15,743 | 3,977 | 41,33 | 28 | 17 | 17 | 9,559 | 3,257 | 34,43 | 18 | 12 | 12 |
| YER154W | OXA1 | 44,82 | 25,60 | 6,45E-117 | 3,429 | 0,003035 | 2,583 | 0,019424 | 2,890 | 0,007793 | 6 | 6 | 14,653 | 3,873 | 57,88 | 6 | 1 | 1 | 11,065 | 3,468 | n. def. | 1 | 4 | 4 |
| YER048W-A | ISD11 | 11,27 | 77,70 | 3,18E-42 | 3,408 | 0,002958 | 2,439 | 0,103671 | -0,108 | 0,961090 | 5 | 5 | 13,260 | 3,729 | 120,57 | 5 | 2 | 2 | 7,392 | 2,886 | 7,90 | 2 | 2 | 2 |
| YKL192C | ACP1 | 13,94 | 35,20 | 3,35E-61 | 3,400 | 0,000829 | 2,746 | 0,023346 | -0,035 | 0,739614 | 2 | 2 | 12,761 | 3,674 | 6,31 | 2 | 1 | 1 | 9,339 | 3,223 | n. def. | 1 | 1 | 1 |
| YML042W | CAT2 | 77,24 | 66,70 | 0 | 3,382 | 0,005705 | 1,046 | 0,205274 | -0,240 | 0,760529 | 28 | 28 | 11,036 | 3,464 | 53,52 | 55 | 28 | 28 | 6,560 | 2,714 | 61,96 | 43 | 24 | 24 |
| YKR009C | FOX2 | 98,70 | 61,30 | 0 | 3,371 | 0,002466 | 1,279 | 0,167941 | -0,372 | 0,979780 | 26 | 26 | 11,304 | 3,499 | 50,00 | 36 | 20 | 20 | 7,521 | 2,911 | 20,26 | 25 | 25 | 25 |
| YJL104W | PAM16 | 16,22 | 31,50 | 4,66E-12 | 3,360 | 0,000400 | 2,074 | 0,018015 | 0,254 | 0,422255 | 2 | 2 | 9,537 | 3,254 | 15,56 | 3 | 0 | 0 | NA | 3,276 | n. def. | 0 | 1 | 1 |
| YIL136W | OM45 | 44,58 | 64,40 | 0 | 3,338 | 0,000560 | 2,928 | 0,046547 | 1,584 | 0,010584 | 20 | 20 | 9,172 | 3,197 | 49,00 | 25 | 8 | 8 | 11,786 | 3,559 | 45,59 | 7 | 15 | 15 |
| YLR077W | FMP25 | 66,69 | 56,10 | 2,59E-203 | 3,338 | 0,005756 | 1,467 | 0,138837 | 3,414 | 0,001693 | 12 | 12 | 16,306 | 4,027 | 202,36 | 13 | 10 | 10 | 9,058 | 3,179 | 74,46 | 8 | 9 | 9 |
| YDR237W | MRPL7 | 33,10 | 20,90 | 3,95E-35 | 3,330 | 0,002027 | 3,313 | 0,002995 | 1,074 | 0,195498 | 4 | 4 | 12,966 | 3,697 | 31,04 | 5 | 3 | 3 | 7,781 | 2,960 | 9,61 | 3 | 1 | 1 |
| YPL134C | ODC1 | 34,21 | 46,80 | 9,24E-82 | 3,293 | 0,010585 | 2,397 | 0,067319 | 2,973 | 0,017419 | 6 | 6 | 6,401 | 2,678 | 86,77 | 6 | 5 | 5 | 19,091 | 4,255 | 97,64 | 3 | 6 | 6 |
| YIL157C | COA1 | 22,04 | 26,40 | 5,90E-18 | 3,287 | 0,021143 | 0,705 | 0,314282 | 2,387 | 0,052397 | 2 | 2 | 3,907 | 1,966 | 274,19 | 2 | 1 | 1 | 11,746 | 3,554 | n. def. | 1 | 0 | 0 |
| YBR037C | SCO1 | 33,17 | 19,70 | 1,45E-32 | 3,274 | 0,010048 | 2,329 | 0,049803 | 2,288 | 0,007646 | 3 | 3 | 8,566 | 3,099 | 69,15 | 2 | 3 | 3 | 17,917 | 4,163 | 59,18 | 2 | 2 | 2 |
| YBR263W | SHM1 | 53,69 | 74,10 | 0 | 3,265 | 0,005026 | 0,748 | 0,303438 | -0,014 | 0,716157 | 17 | 17 | 14,972 | 3,904 | 29,03 | 22 | 14 | 14 | 8,490 | 3,086 | 22,48 | 17 | 15 | 15 |
| YOR222W | ODC2 | 34,01 | 21,20 | 1,58E-23 | 3,258 | 0,001440 | 1,782 | 0,116906 | 0,487 | 0,251553 | 0 | 0 | NA | 2,912 | n. def. | 0 | 2 | 2 | 10,410 | 3,380 | 138,99 | 2 | 2 | 2 |
| YKL141W | SDH3 | 22,07 | 22,20 | 6,59E-36 | 3,233 | 0,041132 | 3,995 | 0,040178 | 0,107 | 0,474419 | 3 | 3 | 17,956 | 4,166 | 4,90 | 2 | 1 | 1 | 19,380 | 4,277 | n. def. | 1 | 0 | 0 |
| YCL017C | NFS1 | 54,47 | 51,90 | 0 | 3,204 | 0,003666 | 2,301 | 0,013588 | -0,111 | 0,957539 | 9 | 9 | 11,936 | 3,577 | 55,88 | 10 | 6 | 6 | 6,345 | 2,666 | 23,55 | 7 | 7 | 7 |
| YFL016C | MDJ1 | 55,56 | 37,00 | 1,42E-82 | 3,166 | 0,002000 | 1,434 | 0,206983 | 0,149 | 0,031102 | 15 | 15 | 11,593 | 3,535 | 97,37 | 17 | 7 | 7 | 7,182 | 2,844 | 37,12 | 7 | 8 | 8 |
| YKR094C;YIL148W;YLR167W | RPL40B; RPL40A | 14,55 | 36,70 | 1,95E-82 | 3,143 | 0,009257 | 2,579 | 0,061719 | 0,751 | 0,265602 | 1 | 1 | 7,280 | 2,864 | n. def. | 1 | 0 | 0 | NA | 2,571 | n. def. | 0 | 1 | 1 |
| YOR211C | MGM1 | 99,18 | 22,40 | 4,33E-61 | 3,128 | 0,012907 | 2,113 | 0,019116 | 0,768 | 0,127802 | 7 | 7 | 12,274 | 3,617 | 69,18 | 7 | 4 | 4 | 4,297 | 2,103 | 155,81 | 4 | 4 | 4 |
| YGR033C | TIM21 | 27,21 | 34,30 | 1,37E-43 | 3,128 | 0,061222 | 3,507 | 0,010229 | 4,726 | 0,019875 | 2 | 2 | 5,738 | 2,521 | 76,55 | 2 | 2 | 2 | 2,650 | 1,406 | n. def. | 1 | 2 | 2 |
| YML030W | RCF1 | 18,48 | 62,30 | 8,68E-111 | 3,109 | 0,007886 | 2,676 | 0,021365 | 3,122 | 0,022439 | 3 | 3 | 9,953 | 3,315 | 78,33 | 4 | 3 | 3 | 12,705 | 3,667 | 37,23 | 3 | 3 | 3 |
| YER050C | RSM18 | 15,84 | 37,00 | 7,24E-24 | 3,054 | 0,007549 | 2,519 | 0,034921 | 3,483 | 0,000022 | 3 | 3 | 9,646 | 3,270 | 109,42 | 3 | 1 | 1 | 4,980 | 2,316 | n. def. | 1 | 0 | 0 |
| YIL070C | MAM33 | 30,13 | 39,50 | 5,17E-237 | 3,020 | 0,000684 | 2,626 | 0,006280 | 5,925 | 0,000040 | 5 | 5 | 9,131 | 3,191 | 46,37 | 5 | 1 | 1 | 7,011 | 2,810 | n. def. | 1 | 3 | 3 |
| YKL148C | SDH1 | 70,23 | 39,20 | 2,74E-160 | 3,012 | 0,002298 | 4,141 | 0,015139 | 0,856 | 0,320382 | 13 | 11 | 8,576 | 3,100 | 132,31 | 10 | 9 | 7 | 6,152 | 2,621 | 133,48 | 5 | 9 | 8 |
| YBR120C | CBP6 | 18,68 | 27,20 | 6,76E-43 | 2,986 | 0,010151 | 2,489 | 0,010020 | 2,393 | 0,051297 | 3 | 3 | 14,048 | 3,812 | 29,30 | 2 | 0 | 0 | NA | 2,353 | n. def. | 0 | 2 | 2 |
| YDR231C | COX20 | 23,76 | 43,90 | 1,23E-67 | 2,982 | 0,013625 | 2,469 | 0,109002 | 2,711 | 0,007546 | 4 | 4 | 12,496 | 3,643 | 19,80 | 4 | 2 | 2 | 9,894 | 3,307 | n. def. | 1 | 1 | 1 |
| YPR024W | YME1 | 81,77 | 26,20 | 1,58E-118 | 2,968 | 0,013543 | 3,049 | 0,044417 | 1,866 | 0,120694 | 9 | 9 | 14,292 | 3,837 | 80,69 | 6 | 4 | 4 | 7,765 | 2,957 | 58,89 | 4 | 7 | 7 |
| YDL067C | COX9 | 6,96 | 39,00 | 3,73E-34 | 2,952 | 0,000917 | 4,752 | 0,003146 | 3,275 | 0,006988 | 0 | 0 | NA | 3,042 | n. def. | 0 | 0 | 0 | NA | 2,702 | n. def. | 0 | 1 | 1 |
| YGR082W | TOM20 | 20,32 | 46,40 | 1,31E-59 | 2,918 | 0,010128 | 3,798 | 0,046857 | 2,363 | 0,017147 | 1 | 1 | 12,311 | 3,622 | n. def. | 1 | 0 | 0 | NA | 2,164 | n. def. | 0 | 1 | 1 |
| YJR100C | AIM25 | 37,48 | 19,00 | 2,66E-20 | 2,890 | 0,006022 | 1,320 | 0,123153 | 1,752 | 0,021379 | 3 | 3 | 7,784 | 2,960 | 29,77 | 3 | 1 | 1 | 4,937 | 2,304 | n. def. | 1 | 0 | 0 |
| YLR203C | MSS51 | 50,88 | 30,50 | 4,43E-42 | 2,865 | 0,013640 | 2,731 | 0,023918 | 2,662 | 0,032242 | 5 | 5 | 7,942 | 2,990 | 55,96 | 4 | 2 | 2 | 3,925 | 1,973 | 23,53 | 2 | 2 | 2 |
| YNL052W | COX5A | 17,14 | 70,60 | 1,19E-203 | 2,784 | 0,019594 | 3,572 | 0,004240 | 5,252 | 0,000825 | 6 | 5 | 13,038 | 3,705 | 113,85 | 6 | 5 | 4 | 7,472 | 2,901 | 149,80 | 4 | 5 | 4 |
| YDR070C | FMP16 | 10,85 | 47,30 | 4,86E-26 | 2,737 | 0,010322 | 1,874 | 0,001581 | 0,678 | 0,099455 | 4 | 4 | 11,335 | 3,503 | 83,14 | 4 | 1 | 1 | 5,866 | 2,552 | n. def. | 1 | 2 | 2 |
| YPL215W | CBP3 | 39,08 | 36,10 | 4,27E-38 | 2,732 | 0,029535 | 3,064 | 0,041730 | 2,538 | 0,001704 | 4 | 4 | 16,900 | 4,079 | 42,16 | 4 | 3 | 3 | 3,390 | 1,761 | 207,58 | 3 | 2 | 2 |
| YGR222W | PET54 | 34,64 | 39,20 | 3,60E-42 | 2,729 | 0,000047 | 1,891 | 0,034782 | 0,016 | 0,407575 | 3 | 3 | 6,866 | 2,779 | 63,09 | 3 | 3 | 3 | 6,577 | 2,717 | 59,05 | 3 | 3 | 3 |
| YHR008C | SOD2 | 25,77 | 30,00 | 4,01E-46 | 2,687 | 0,010294 | 1,721 | 0,059495 | 1,498 | 0,082000 | 3 | 3 | 11,078 | 3,470 | 49,51 | 3 | 3 | 3 | 4,848 | 2,277 | 21,36 | 3 | 3 | 3 |
| YER014W | HEM14 | 59,70 | 28,20 | 1,10E-44 | 2,635 | 0,002166 | 1,008 | 0,253117 | 3,189 | 0,064151 | 6 | 6 | 7,849 | 2,973 | 67,87 | 5 | 0 | 0 | NA | 2,392 | n. def. | 0 | 6 | 6 |
| YIL051C | MMF1 | 15,91 | 58,60 | 1,47E-43 | 2,610 | 0,012552 | 1,621 | 0,082675 | 2,122 | 0,063364 | 1 | 1 | 10,388 | 3,377 | 14,10 | 2 | 1 | 1 | 5,769 | 2,528 | n. def. | 1 | 3 | 3 |
| YJL208C | NUC1 | 37,21 | 53,50 | 2,68E-62 | 2,608 | 0,002431 | 1,273 | 0,180863 | 1,045 | 0,142798 | 9 | 9 | 4,778 | 2,256 | 20,17 | 11 | 5 | 5 | 6,487 | 2,697 | 14,32 | 6 | 4 | 4 |
| YJR121W | ATP2 | 54,79 | 83,40 | 0 | 2,589 | 0,005887 | 2,181 | 0,065505 | 0,671 | 0,000221 | 13 | 13 | 8,560 | 3,098 | 40,98 | 15 | 15 | 15 | 4,340 | 2,118 | 21,35 | 19 | 14 | 14 |
| YIL111W | COX5B | 17,20 | 50,30 | 8,14E-117 | 2,565 | 0,007237 | 1,407 | 0,153732 | 3,205 | 0,007298 | 1 | 0 | NA | 1,952 | n. def. | 0 | 2 | 1 | 6,826 | 2,771 | n. def. | 1 | 2 | 1 |
| YLR355C | ILV5 | 44,37 | 57,70 | 2,63E-169 | 2,549 | 0,003784 | 2,002 | 0,061537 | 3,846 | 0,082122 | 9 | 9 | 7,890 | 2,980 | 18,23 | 9 | 6 | 6 | 4,702 | 2,233 | 23,38 | 7 | 8 | 8 |
| YBL099W | ATP1 | 58,61 | 59,10 | 0 | 2,520 | 0,006098 | 2,189 | 0,071196 | 0,553 | 0,003662 | 14 | 14 | 7,561 | 2,919 | 31,15 | 16 | 14 | 14 | 3,939 | 1,978 | 18,25 | 15 | 18 | 18 |
| YPR001W | CIT3 | 53,81 | 47,70 | 3,88E-107 | 2,509 | 0,002466 | 1,450 | 0,154442 | -0,109 | 0,919697 | 15 | 15 | 6,304 | 2,656 | 55,15 | 18 | 11 | 11 | 4,460 | 2,157 | 15,67 | 14 | 9 | 9 |
| YGR112W | SHY1 | 45,06 | 29,30 | 1,31E-62 | 2,502 | 0,052909 | 0,272 | 0,449734 | 2,246 | 0,024567 | 4 | 4 | 17,611 | 4,138 | 124,18 | 3 | 1 | 1 | 2,137 | 1,095 | n. def. | 1 | 2 | 2 |
| YPL078C | ATP4 | 26,93 | 48,80 | 4,87E-70 | 2,499 | 0,007738 | 1,965 | 0,071880 | 0,610 | 0,003197 | 3 | 3 | 8,648 | 3,112 | 7,69 | 3 | 3 | 3 | 4,202 | 2,071 | 4,02 | 3 | 0 | 0 |
| YLR253W | MCP2 | 65,92 | 14,90 | 9,99E-33 | 2,478 | 0,010541 | 2,342 | 0,056723 | 3,389 | 0,007759 | 3 | 3 | 6,330 | 2,662 | 120,28 | 3 | 4 | 4 | 3,419 | 1,773 | n. def. | 1 | 3 | 3 |
| YPL099C | AIM43 | 21,84 | 40,70 | 3,18E-72 | 2,467 | 0,017999 | 2,349 | 0,003560 | 1,320 | 0,192716 | 5 | 5 | 9,948 | 3,314 | 79,07 | 3 | 1 | 1 | 5,419 | 2,438 | 13,98 | 2 | 2 | 2 |
| YMR089C | YTA12 | 93,28 | 22,10 | 1,09E-65 | 2,459 | 0,003799 | 1,327 | 0,138576 | 0,569 | 0,393015 | 12 | 12 | 7,363 | 2,880 | 129,38 | 11 | 4 | 4 | 4,493 | 2,168 | 22,92 | 4 | 5 | 5 |
| YDR377W | ATP17 | 11,31 | 29,70 | 4,97E-12 | 2,432 | 0,013516 | 2,230 | 0,056364 | 1,248 | 0,000966 | 1 | 1 | 3,088 | 1,627 | n. def. | 1 | 0 | 0 | NA | 2,719 | n. def. | 0 | 1 | 1 |
| YJR019C | TES1 | 40,26 | 22,10 | 3,50E-33 | 2,406 | 0,019377 | 1,199 | 0,203544 | 0,039 | 0,186522 | 0 | 0 | NA | 1,874 | n. def. | 0 | 2 | 2 | 3,900 | 1,963 | 3,78 | 2 | 0 | 0 |
| YLR259C | HSP60 | 60,75 | 63,30 | 0 | 2,398 | 0,000558 | 0,946 | 0,236129 | 0,358 | 0,000922 | 25 | 25 | 5,473 | 2,452 | 53,60 | 29 | 24 | 24 | 5,666 | 2,502 | 32,28 | 27 | 22 | 22 |
| YOR142W | LSC1 | 35,03 | 33,70 | 2,63E-81 | 2,365 | 0,008914 | 1,812 | 0,092730 | 3,193 | 0,000041 | 3 | 3 | 7,808 | 2,965 | 18,21 | 3 | 2 | 2 | 4,781 | 2,257 | 18,94 | 2 | 4 | 4 |
| YNL055C | POR1 | 30,43 | 67,80 | 1,57E-219 | 2,361 | 0,014551 | 1,316 | 0,108453 | 2,709 | 0,049132 | 5 | 5 | 8,661 | 3,115 | 62,32 | 6 | 8 | 8 | 3,240 | 1,696 | 14,48 | 9 | 8 | 8 |
| YIL077C | RCI37 | 36,95 | 33,80 | 2,89E-60 | 2,336 | 0,010941 | 0,987 | 0,090895 | 2,123 | 0,024062 | 7 | 7 | 8,203 | 3,036 | 64,31 | 5 | 1 | 1 | 3,829 | 1,937 | n. def. | 1 | 1 | 1 |
| YBR039W | ATP3 | 34,35 | 54,00 | 2,06E-77 | 2,327 | 0,004980 | 1,875 | 0,044039 | 3,091 | 0,004008 | 4 | 4 | 6,931 | 2,793 | 36,73 | 3 | 2 | 2 | 4,174 | 2,061 | 18,08 | 2 | 2 | 2 |
| YDL181W | INH1 | 9,87 | 50,60 | 9,37E-65 | 2,320 | 0,049818 | 1,489 | 0,043150 | 0,574 | 0,006570 | 0 | 0 | NA | 3,366 | n. def. | 0 | 0 | 0 | NA | 2,829 | n. def. | 0 | 0 | 0 |
| YMR256C | COX7 | 6,93 | 83,30 | 7,10E-30 | 2,306 | 0,035605 | 3,488 | 0,057549 | 7,078 | 0,002340 | 1 | 1 | 2,215 | 1,147 | n. def. | 1 | 1 | 1 | 5,177 | 2,372 | n. def. | 1 | 0 | 0 |
| YKL085W | MDH1 | 35,65 | 85,00 | 1,52E-215 | 2,302 | 0,000914 | 1,629 | 0,102356 | 2,496 | 0,009403 | 5 | 5 | 5,634 | 2,494 | 20,57 | 5 | 5 | 5 | 4,490 | 2,167 | 10,41 | 6 | 5 | 5 |
| YPL132W | COX11 | 34,04 | 21,30 | 1,16E-16 | 2,299 | 0,021816 | 2,402 | 0,035192 | 3,154 | 0,006497 | 2 | 2 | 9,686 | 3,276 | 1,57 | 2 | 1 | 1 | 3,902 | 1,964 | n. def. | 1 | 1 | 1 |
| YML078W | CPR3 | 19,92 | 42,90 | 5,16E-54 | 2,201 | 0,003688 | 2,482 | 0,003787 | 1,463 | 0,018706 | 2 | 2 | 4,550 | 2,186 | 38,36 | 2 | 2 | 2 | 3,680 | 1,880 | 82,18 | 2 | 5 | 5 |
| Q0045;Q0070;Q0060;Q005 | COX1; AI5_ALPH | 58,80 | 14,40 | 1,84E-90 | 2,182 | 0,038309 | 3,830 | 0,035971 | 7,501 | 0,000080 | 1 | 1 | 10,206 | 3,351 | n. def. | 1 | 0 | 0 | NA | 2,056 | n. def. | 0 | 0 | 0 |
| YOR020C | HSP10 | 11,37 | 53,80 | 1,45E-21 | 2,155 | 0,037418 | 1,428 | 0,128825 | 2,431 | 0,132065 | 0 | 0 | NA | 3,337 | n. def. | 0 | 1 | 1 | 3,786 | 1,921 | n. def. | 1 | 1 | 1 |
| YPL271W | ATP15 | 6,74 | 90,30 | 1,25E-65 | 2,139 | 0,005054 | 1,712 | 0,067888 | 2,717 | 0,018024 | 2 | 2 | 5,273 | 2,399 | 51,01 | 2 | 2 | 2 | 3,269 | 1,709 | 18,06 | 2 | 1 | 1 |
| YKL195W | MIA40 | 44,54 | 17,90 | 1,09E-29 | 2,101 | 0,046316 | 2,004 | 0,233109 | 0,499 | 0,004146 | 3 | 3 | 3,318 | 1,730 | 83,15 | 3 | 1 | 1 | 10,810 | 3,434 | n. def. | 1 | 1 | 1 |
| YBL064C | PRX1 | 29,50 | 54,80 | 7,79E-75 | 2,090 | 0,001046 | 1,697 | 0,101883 | 0,460 | 0,395127 | 8 | 8 | 4,389 | 2,134 | 20,75 | 9 | 8 | 8 | 3,750 | 1,907 | 16,82 | 9 | 7 | 7 |
| YOR271C | FSF1 | 35,41 | 32,70 | 3,45E-35 | 2,070 | 0,006073 | 0,389 | 0,416163 | 1,872 | 0,037255 | 3 | 3 | 4,016 | 2,006 | 134,54 | 2 | 1 | 1 | 3,266 | 1,708 | n. def. | 1 | 2 | 2 |
| YGL008C;YPL036W | PMA1; PMA2 | 99,62 | 19,90 | 2,96E-95 | 2,054 | 0,018575 | 0,939 | 0,192911 | 2,817 | 0,097644 | 10 | 10 | 3,129 | 1,646 | 22,32 | 13 | 6 | 6 | 3,134 | 1,648 | 11,83 | 6 | 7 | 7 |
| YLR304C;YJL200C | ACO1; ACO2 | 85,37 | 47,80 | 3,97E-259 | 2,050 | 0,004569 | 1,638 | 0,074871 | 2,075 | 0,111089 | 11 | 11 | 5,428 | 2,440 | 28,31 | 12 | 10 | 10 | 3,498 | 1,806 | 19,14 | 10 | 10 | 10 |
| YNR018W | RCF2 | 25,34 | 39,70 | 6,40E-68 | 2,040 | 0,045687 | 2,076 | 0,039596 | 4,788 | 0,004765 | 3 | 3 | NA | 3,326 | n. def. | 0 | 0 | 0 | NA | 1,119 | n. def. | 0 | 2 | 2 |
| YDR298C | ATP5 | 22,81 | 52,80 | 1,10E-86 | 2,007 | 0,010014 | 1,962 | 0,051630 | 2,758 | 0,047851 | 3 | 3 | 5,525 | 2,466 | 20,33 | 3 | 2 | 2 | 2,780 | 1,475 | 18,24 | 2 | 2 | 2 |
| YKL016C | ATP7 | 19,81 | 68,40 | 7,06E-56 | 1,969 | 0,001421 | 1,879 | 0,049953 | 1,578 | 0,149307 | 2 | 2 | 4,320 | 2,111 | 46,40 | 2 | 4 | 4 | 3,395 | 1,763 | 22,83 | 4 | 5 | 5 |
| YLR395C | COX8 | 8,91 | 15,40 | 1,32E-17 | 1,954 | 0,063313 | 5,091 | 0,002445 | 5,247 | 0,006018 | 0 | 0 | NA | 3,473 | n. def. | 0 | 0 | 0 | NA | 0,973 | n. def. | 0 | 0 | 0 |
| YNR001C | CIT1 | 53,36 | 60,10 | 4,51E-142 | 1,917 | 0,031911 | 1,298 | 0,137098 | 0,075 | 0,218939 | 14 | 13 | 2,266 | 1,180 | 22,08 | 15 | 11 | 11 | 3,197 | 1,677 | 14,00 | 12 | 14 | 13 |
| YCR010C | ADY2 | 30,73 | 13,10 | 1,80E-32 | 1,895 | 0,001505 | 1,278 | 0,159864 | 1,153 | 0,155154 | 2 | 2 | 3,473 | 1,796 | 4,12 | 2 | 2 | 2 | 3,446 | 1,785 | 9,11 | 2 | 2 | 2 |
| YNL208W | YNL208W | 20,15 | 44,70 | 8,96E-45 | 1,857 | 0,029654 | 2,337 | 0,086394 | 0,240 | 0,358552 | 3 | 3 | 3,930 | 1,974 | 24,37 | 3 | 0 | 0 | NA | 0,984 | n. def. | 0 | 2 | 2 |
| YPR002W | PDH1 | 57,68 | 15,50 | 9,93E-22 | 1,841 | 0,032864 | 0,750 | 0,162923 | 1,394 | 0,038042 | 3 | 3 | 2,228 | 1,156 | 28,93 | 3 | 0 | 0 | NA | 2,807 | n. def. | 0 | 1 | 1 |
| YDL004W | ATP16 | 17,02 | 45,60 | 6,23E-66 | 1,839 | 0,002842 | 1,654 | 0,083435 | 0,609 | 0,002125 | 1 | 1 | 3,125 | 1,644 | 55,12 | 2 | 1 | 1 | 4,312 | 2,108 | n. def. | 1 | 1 | 1 |
| YIL155C | GUT2 | 72,39 | 32,20 | 4,36E-105 | 1,825 | 0,000663 | 0,900 | 0,135687 | 1,815 | 0,060625 | 13 | 13 | 3,243 | 1,697 | 29,52 | 15 | 9 | 9 | 3,624 | 1,857 | 29,73 | 10 | 8 | 8 |
| YJR048W | CYC1 | 12,18 | 73,40 | 7,46E-169 | 1,824 | 0,007689 | 1,423 | 0,170974 | -0,192 | 0,995455 | 5 | 4 | 3,290 | 1,718 | 21,68 | 5 | 1 | 1 | 2,813 | 1,492 | n. def. | 1 | 4 | 3 |
| YOR176W | HEM15 | 44,60 | 65,40 | 7,92E-151 | 1,784 | 0,002600 | 1,054 | 0,209357 | -0,058 | 0,903350 | 17 | 17 | 3,971 | 1,989 | 23,31 | 24 | 12 | 12 | 2,920 | 1,546 | 19,60 | 18 | 16 | 16 |
| YKL133C | RCI50 | 54,49 | 26,30 | 4,46E-41 | 1,767 | 0,005650 | 1,391 | 0,102994 | 2,487 | 0,038369 | 3 | 3 | 4,241 | 2,084 | 141,21 | 2 | 2 | 2 | 3,451 | 1,787 | 168,70 | 2 | 3 | 3 |
| YKL065W-A | DPC7 | 8,53 | 52,10 | 4,05E-29 | 1,752 | 0,007355 | 2,128 | 0,126384 | 0,923 | 0,211154 | 2 | 2 | 2,866 | 1,519 | 50,13 | 7 | 2 | 2 | 4,537 | 2,182 | 15,23 | 3 | 3 | 3 |
| YNL104C;YOR108W | LEU4; LEU9 | 68,41 | 25,50 | 2,69E-51 | 1,735 | 0,003860 | 1,305 | 0,151849 | 2,722 | 0,083019 | 3 | 3 | 4,065 | 2,023 | 21,62 | 3 | 2 | 2 | 2,829 | 1,500 | 44,30 | 2 | 2 | 2 |
| YKL150W | MCR1 | 34,14 | 65,90 | 1,41E-134 | 1,677 | 0,001216 | 0,858 | 0,167543 | -0,076 | 0,929805 | 8 | 8 | 3,574 | 1,838 | 28,28 | 9 | 7 | 7 | 2,952 | 1,561 | 18,00 | 7 | 10 | 10 |
| YDR148C | KGD2 | 50,43 | 51,60 | 2,23E-200 | 1,676 | 0,015880 | 0,482 | 0,342087 | -0,029 | 0,586368 | 19 | 19 | 3,447 | 1,785 | 25,97 | 29 | 15 | 15 | 2,144 | 1,100 | 36,45 | 21 | 17 | 17 |
| YGL068W | MNP1 | 20,65 | 50,00 | 6,43E-53 | 1,661 | 0,023999 | 1,806 | 0,069425 | -0,080 | 0,918890 | 3 | 3 | 2,627 | 1,393 | 18,06 | 4 | 4 | 4 | 2,270 | 1,183 | 24,64 | 6 | 1 | 1 |
| YIL125W | KGD1 | 114,41 | 59,00 | 0 | 1,632 | 0,017717 | 0,362 | 0,382116 | 0,157 | 0,022474 | 39 | 39 | 3,241 | 1,696 | 27,63 | 58 | 37 | 37 | 2,080 | 1,056 | 11,03 | 52 | 34 | 34 |
| YNL037C | IDH1 | 39,32 | 41,40 | 7,41E-58 | 1,606 | 0,001400 | 2,158 | 0,038850 | 3,419 | 0,067794 | 2 | 2 | 3,054 | 1,611 | 10,42 | 3 | 1 | 1 | 2,744 | 1,456 | n. def. | 1 | 1 | 1 |
| YFR049W | YMR31 | 13,69 | 56,10 | 3,67E-45 | 1,595 | 0,012048 | 0,283 | 0,403813 | -0,417 | 0,849996 | 5 | 5 | 3,218 | 1,686 | 106,62 | 6 | 5 | 5 | 2,173 | 1,119 | 25,81 | 6 | 3 | 3 |
| YOR136W | IDH2 | 39,74 | 22,50 | 5,09E-64 | 1,582 | 0,001017 | 1,020 | 0,104493 | 2,909 | 0,019258 | 2 | 2 | 3,168 | 1,663 | 1,04 | 2 | 2 | 2 | 2,712 | 1,439 | 1,80 | 2 | 2 | 2 |
| YNL121C | TOM70 | 70,12 | 33,10 | 5,48E-148 | 1,474 | 0,009362 | 2,601 | 0,028075 | -0,255 | 0,716686 | 3 | 3 | 3,325 | 1,734 | 22,46 | 3 | 0 | 0 | NA | 1,070 | n. def. | 0 | 1 | 1 |
| YER004W | FMP52 | 25,09 | 64,10 | 1,20E-104 | 1,385 | 0,004936 | 1,678 | 0,120494 | -0,058 | 0,946304 | 6 | 6 | 2,406 | 1,267 | 14,74 | 6 | 5 | 5 | 3,164 | 1,662 | 36,32 | 5 | 7 | 7 |
| YJR045C;YEL030W | SSC1; ECM10 | 70,63 | 60,40 | 0 | 1,369 | 0,005316 | 1,005 | 0,212829 | 1,135 | 0,000322 | 27 | 27 | 2,231 | 1,158 | 30,09 | 33 | 23 | 23 | 2,478 | 1,309 | 35,61 | 28 | 21 | 21 |
| YFL018C | LPD1 | 54,01 | 67,50 | 0 | 1,349 | 0,026209 | 0,065 | 0,469231 | 0,122 | 0,114859 | 28 | 28 | 2,649 | 1,405 | 21,83 | 38 | 27 | 27 | 1,701 | 0,766 | 16,48 | 38 | 19 | 19 |
| YBL015W | ACH1 | 58,71 | 32,30 | 1,75E-57 | 1,349 | 0,007768 | 1,367 | 0,004793 | 1,808 | 0,027589 | 7 | 7 | 3,218 | 1,686 | 103,27 | 7 | 1 | 1 | 2,208 | 1,143 | n. def. | 1 | 2 | 2 |
| YDR322C-A | TIM11 | 10,88 | 54,20 | 4,13E-39 | 1,339 | 0,003687 | 1,680 | 0,126302 | 0,797 | 0,045555 | 1 | 1 | 2,750 | 1,460 | n. def. | 1 | 1 | 1 | 2,732 | 1,450 | n. def. | 1 | 0 | 0 |
| YOR374W | ALD4 | 56,72 | 79,20 | 0 | 1,312 | 0,003402 | 1,020 | 0,174607 | 0,492 | 0,000761 | 28 | 27 | 2,799 | 1,485 | 29,38 | 37 | 26 | 25 | 2,160 | 1,111 | 20,21 | 32 | 30 | 28 |
| YPR058W | YMC1 | 33,38 | 21,80 | 3,93E-39 | 1,292 | 0,261781 | 1,499 | 0,212353 | 3,724 | 0,000121 | 3 | 3 | 6,752 | 2,755 | 186,59 | 2 | 1 | 1 | 0,238 | -2,070 | n. def. | 1 | 0 | 0 |
| YBR026C | ETR1 | 42,07 | 38,90 | 3,83E-43 | 1,164 | 0,005406 | 1,822 | 0,094727 | 1,442 | 0,189951 | 7 | 7 | 2,062 | 1,044 | 17,71 | 7 | 2 | 2 | 2,653 | 1,408 | 48,48 | 2 | 5 | 5 |
| YJR016C | ILV3 | 62,86 | 42,20 | 1,15E-107 | 1,124 | 0,003514 | 1,044 | 0,203396 | 3,848 | 0,013477 | 13 | 13 | 1,968 | 0,977 | 18,53 | 16 | 3 | 3 | 2,464 | 1,301 | 15,33 | 4 | 12 | 12 |
| YGR244C | LSC2 | 46,90 | 50,80 | 6,12E-118 | 1,083 | 0,098483 | 0,728 | 0,020319 | 0,339 | 0,439441 | 13 | 13 | 1,814 | 0,859 | 25,49 | 14 | 3 | 3 | 4,469 | 2,160 | 13,01 | 4 | 12 | 12 |
| YOL109W | ZEO1 | 12,59 | 28,30 | 3,39E-28 | 0,991 | 0,005605 | 1,172 | 0,284751 | 0,275 | 0,023140 | 1 | 1 | 2,188 | 1,130 | n. def. | 1 | 1 | 1 | 1,721 | 0,783 | n. def. | 1 | 1 | 1 |
| YOL077W-A | ATP19 | 7,53 | 72,10 | 1,37E-23 | 0,976 | 0,023074 | 1,124 | 0,197700 | 0,325 | 0,398959 | 3 | 3 | 1,663 | 0,733 | 19,03 | 3 | 2 | 2 | 1,724 | 0,785 | 56,10 | 2 | 2 | 2 |
| YJR144W | MGM101 | 30,09 | 50,90 | 8,95E-96 | 0,893 | 0,052006 | -0,314 | 0,606705 | 3,133 | 0,037396 | 13 | 13 | 1,252 | 0,324 | 28,23 | 18 | 12 | 12 | 1,934 | 0,951 | 49,08 | 15 | 12 | 12 |
| YIL065C | FIS1 | 17,73 | 38,10 | 3,54E-27 | 0,625 | 0,002807 | 0,939 | 0,237822 | 2,437 | 0,087820 | 6 | 6 | 1,556 | 0,638 | 124,42 | 6 | 4 | 4 | 1,451 | 0,537 | 109,26 | 4 | 4 | 4 |
| YNL071W | LAT1 | 51,82 | 49,60 | 1,24E-155 | 0,474 | 0,193371 | -0,309 | 0,602603 | 0,390 | 0,036482 | 13 | 13 | 1,309 | 0,389 | 27,24 | 16 | 12 | 12 | 0,854 | -0,227 | 31,66 | 14 | 8 | 8 |
| YMR108W | ILV2 | 74,94 | 48,00 | 2,35E-190 | 0,460 | 0,020726 | 0,112 | 0,468045 | 1,089 | 0,118707 | 19 | 19 | 1,398 | 0,484 | 24,87 | 30 | 5 | 5 | 1,532 | 0,615 | 20,97 | 5 | 18 | 18 |

| Mar26 - Replicate 3 | | | | Mic60 - Replicate 1 | | | | | | Mic60 - Replicate 2 | | | | | | Mic60 - Replicate 3 | | | | | | Cor1 - Replicate 1 | | | |
|---|---|---|---|---|---|---|---|---|---|---|---|---|---|---|---|---|---|---|---|---|---|---|---|---|---|
| Ratio Mar26/WT | log2 Ratio Mar26/WT | Ratio Variability [%] | Ratio Count | Peptides | Unique Peptides | Ratio Mic60/WT | log2 Ratio Mic60/WT | Ratio Variability [%] | Ratio Count | Peptides | Unique Peptides | Ratio Mic60/WT | log2 Ratio Mic60/WT | Ratio Variability [%] | Ratio Count | Peptides | Unique Peptides | Ratio Mic60/WT | log2 Ratio Mic60/WT | Ratio Variability [%] | Ratio Count | Peptides | Unique Peptides | Ratio Cor1/WT | log2 Ratio Cor1/WT |
| 135,000 | 7,077 | 139,89 | 17 | 2 | 2 | 11,640 | 3,541 | 17,43 | 2 | 3 | 3 | 23,273 | 4,541 | n. def. | 1 | 12 | 12 | 17,703 | 4,146 | 205,80 | 7 | 13 | 13 | 11,717 | 3,551 |
| 76,546 | 6,258 | 123,29 | 25 | 23 | 23 | 11,785 | 3,559 | 24,56 | 27 | 23 | 23 | 17,108 | 4,097 | 47,20 | 26 | 24 | 24 | 24,951 | 4,641 | 157,40 | 20 | 50 | 50 | 19,649 | 4,296 |
| 62,305 | 5,961 | 131,31 | 29 | 24 | 24 | 11,182 | 3,483 | 24,11 | 31 | 28 | 28 | 16,818 | 4,072 | 71,82 | 34 | 31 | 31 | 59,719 | 5,900 | 127,97 | 31 | 61 | 61 | 23,742 | 4,569 |
| 73,432 | 6,198 | 329,05 | 4 | 3 | 3 | 16,337 | 4,030 | 21,55 | 2 | 2 | 2 | 27,789 | 4,796 | 25,66 | 2 | 3 | 3 | 9,651 | 3,271 | n. def. | 1 | 7 | 7 | 67,700 | 6,081 |
| 56,808 | 5,828 | 97,51 | 31 | 29 | 29 | 33,578 | 5,069 | 63,22 | 32 | 32 | 31 | 27,711 | 4,792 | 88,44 | 22 | 39 | 39 | 27,598 | 4,786 | 130,08 | 31 | 8 | 8 | 10,564 | 3,401 |
| 76,693 | 6,261 | 79,43 | 4 | 7 | 7 | 41,785 | 5,385 | 100,73 | 7 | 7 | 7 | 23,351 | 4,545 | 137,73 | 8 | 9 | 9 | 26,126 | 4,707 | 217,67 | 7 | 2 | 2 | 20,334 | 4,346 |
| 64,243 | 6,005 | 45,61 | 2 | 1 | 1 | 37,187 | 5,217 | 9,38 | 3 | 1 | 1 | 37,813 | 5,241 | 33,30 | 3 | 1 | 1 | 64,267 | 6,006 | 30,41 | 3 | 7 | 7 | 62,174 | 5,958 |
| 28,564 | 4,836 | 126,75 | 19 | 7 | 7 | 14,553 | 3,863 | 52,63 | 7 | 2 | 2 | 11,272 | 3,495 | 5,72 | 2 | 3 | 3 | 11,668 | 3,544 | 103,19 | 2 | 1 | 1 | 3,078 | 1,622 |
| 31,867 | 4,994 | 151,48 | 8 | 6 | 6 | 15,500 | 3,954 | 42,55 | 8 | 6 | 6 | 19,844 | 4,311 | 50,65 | 7 | 9 | 9 | 66,024 | 6,045 | 174,53 | 9 | 16 | 16 | 81,960 | 6,357 |
| 56,590 | 5,822 | 112,40 | 23 | 34 | 34 | 58,514 | 5,871 | 149,52 | 38 | 35 | 35 | 53,248 | 5,735 | 152,57 | 38 | 49 | 49 | 29,885 | 4,901 | 142,47 | 55 | 2 | 2 | 6,020 | 2,590 |
| 29,763 | 4,895 | 49,85 | 8 | 6 | 6 | 14,039 | 3,811 | 48,61 | 8 | 5 | 5 | 7,060 | 2,820 | 94,30 | 5 | 4 | 4 | 36,323 | 5,183 | 110,55 | 4 | 10 | 10 | 99,721 | 6,640 |
| 29,057 | 4,861 | 67,13 | 5 | 7 | 7 | 12,393 | 3,631 | 89,76 | 10 | 6 | 6 | 9,869 | 3,303 | 107,07 | 8 | 5 | 5 | 6,724 | 2,749 | 158,91 | 8 | 12 | 12 | 80,756 | 6,335 |
| 39,300 | 5,296 | 157,17 | 25 | 16 | 14 | 11,148 | 3,479 | 63,69 | 27 | 17 | 14 | 20,740 | 4,374 | 123,18 | 21 | 23 | 20 | 24,588 | 4,620 | 147,52 | 21 | 26 | 24 | 36,614 | 5,194 |
| 18,938 | 4,243 | 87,05 | 7 | 5 | 5 | 13,722 | 3,778 | 30,93 | 6 | 6 | 6 | 11,740 | 3,553 | 111,82 | 6 | 10 | 10 | 8,347 | 3,061 | 125,29 | 6 | 13 | 13 | 319,857 | 8,321 |
| 27,857 | 4,800 | 79,34 | 11 | 9 | 7 | 0,744 | -0,426 | 16,47 | 7 | 8 | 6 | 1,334 | 0,415 | 25,37 | 6 | 6 | 5 | 7,670 | 2,939 | 43,11 | 5 | 9 | 8 | 0,774 | -0,369 |
| 23,866 | 4,577 | 66,43 | 7 | 10 | 10 | 58,082 | 5,860 | 56,80 | 9 | 10 | 10 | 35,200 | 5,138 | 118,12 | 9 | 9 | 9 | 48,237 | 5,592 | 154,39 | 7 | 1 | 1 | 5,073 | 2,343 |
| 24,556 | 4,618 | 86,54 | 4 | 4 | 4 | 12,023 | 3,588 | 63,77 | 7 | 5 | 5 | 9,091 | 3,184 | 67,62 | 5 | 4 | 4 | 18,987 | 4,247 | 89,56 | 3 | 11 | 11 | 54,298 | 5,763 |
| 20,118 | 4,330 | 100,61 | 18 | 13 | 13 | 15,834 | 3,985 | 77,67 | 16 | 8 | 8 | 22,850 | 4,514 | 83,50 | 10 | 17 | 17 | 6,357 | 2,668 | 173,65 | 13 | 0 | 0 | NA | 2,088 |
| 14,511 | 3,859 | 43,86 | 13 | 13 | 13 | 16,060 | 4,005 | 36,82 | 15 | 13 | 13 | 15,762 | 3,978 | 71,23 | 12 | 15 | 15 | 25,320 | 4,662 | 134,15 | 7 | 0 | 0 | NA | 0,681 |
| 22,676 | 4,503 | 136,17 | 12 | 9 | 9 | 13,223 | 3,725 | 19,38 | 13 | 10 | 10 | 19,628 | 4,295 | 50,59 | 14 | 13 | 13 | 29,202 | 4,868 | 164,97 | 16 | 16 | 16 | 92,567 | 6,532 |
| 27,147 | 4,763 | 133,06 | 12 | 9 | 9 | 18,710 | 4,226 | 50,42 | 12 | 4 | 4 | 24,699 | 4,626 | 36,37 | 7 | 8 | 8 | 51,125 | 5,676 | 162,15 | 8 | 15 | 15 | 18,725 | 4,227 |
| 28,236 | 4,819 | 69,36 | 11 | 5 | 3 | 0,748 | -0,418 | 44,65 | 6 | 4 | 2 | 1,550 | 0,633 | 30,20 | 3 | 4 | 3 | 5,437 | 2,443 | 22,57 | 3 | 7 | 6 | 0,758 | -0,400 |
| 18,298 | 4,194 | 73,71 | 5 | 5 | 5 | 14,539 | 3,862 | 44,60 | 6 | 5 | 5 | 22,411 | 4,486 | 57,78 | 5 | 5 | 5 | 44,263 | 5,468 | 122,22 | 4 | 0 | 0 | NA | 3,624 |
| 17,080 | 4,094 | 53,65 | 6 | 6 | 6 | 10,676 | 3,416 | 34,16 | 5 | 9 | 9 | 10,086 | 3,334 | 94,07 | 7 | 7 | 7 | 11,411 | 3,512 | 125,52 | 3 | 0 | 0 | NA | 4,070 |
| 28,638 | 4,840 | 85,05 | 12 | 0 | 0 | NA | 0,203 | n. def. | 0 | 0 | 0 | NA | 1,633 | n. def. | 0 | 0 | 0 | NA | 3,018 | n. def. | 0 | 10 | 10 | 9,062 | 3,180 |
| 25,278 | 4,660 | n. def. | 1 | 5 | 5 | 18,972 | 4,246 | 132,47 | 5 | 4 | 4 | 18,185 | 4,185 | 88,60 | 2 | 4 | 4 | 11,866 | 3,569 | 140,41 | 2 | 0 | 0 | NA | 1,086 |
| 17,049 | 4,092 | 215,88 | 6 | 12 | 12 | 56,928 | 5,831 | 133,71 | 10 | 15 | 15 | 31,082 | 4,958 | 127,77 | 13 | 20 | 20 | 23,810 | 4,573 | 134,43 | 15 | 0 | 0 | NA | 3,145 |
| 12,625 | 3,658 | 194,66 | 24 | 11 | 11 | 20,478 | 4,356 | 104,37 | 10 | 14 | 13 | 12,049 | 3,591 | 74,91 | 12 | 20 | 20 | 10,597 | 3,406 | 136,06 | 13 | 3 | 3 | 10,469 | 3,388 |
| 28,460 | 4,831 | 195,06 | 4 | 8 | 8 | 47,148 | 5,559 | 141,70 | 4 | 7 | 7 | 39,355 | 5,298 | 95,97 | 4 | 12 | 12 | 3,334 | 1,737 | 206,27 | 5 | 0 | 0 | NA | 4,414 |
| 11,657 | 3,543 | n. def. | 1 | 2 | 2 | 17,605 | 4,138 | 14,65 | 3 | 2 | 2 | 27,165 | 4,764 | 25,24 | 3 | 2 | 2 | 108,814 | 6,766 | 38,87 | 3 | 3 | 3 | 135,459 | 7,082 |
| 33,307 | 5,058 | 132,04 | 7 | 9 | 9 | 17,231 | 4,107 | 83,13 | 9 | 11 | 11 | 16,781 | 4,069 | 128,69 | 12 | 6 | 6 | 16,973 | 4,085 | 129,03 | 7 | 0 | 0 | NA | -1,730 |
| 11,192 | 3,484 | 78,99 | 5 | 3 | 3 | 26,457 | 4,726 | 30,13 | 2 | 3 | 3 | 41,210 | 5,365 | 37,57 | 2 | 6 | 6 | 16,255 | 4,023 | 240,61 | 2 | 7 | 7 | 23,281 | 4,541 |
| 26,389 | 4,722 | 85,40 | 7 | 7 | 7 | 10,327 | 3,368 | 13,90 | 7 | 4 | 4 | 19,308 | 4,271 | 27,42 | 4 | 7 | 7 | 8,708 | 3,122 | n. def. | 1 | 2 | 2 | 3,358 | 1,747 |
| 8,559 | 3,097 | 101,78 | 7 | 9 | 9 | 9,867 | 3,303 | 67,24 | 11 | 12 | 12 | 11,437 | 3,516 | 154,81 | 10 | 11 | 11 | 9,902 | 3,308 | 166,38 | 11 | 0 | 0 | NA | 0,928 |
| 23,949 | 4,582 | 79,69 | 4 | 0 | 0 | NA | -0,021 | n. def. | 0 | 0 | 0 | NA | 3,092 | n. def. | 0 | 0 | 0 | NA | 3,763 | n. def. | 0 | 2 | 2 | 4,452 | 2,154 |
| 23,879 | 4,578 | 102,39 | 6 | 1 | 1 | 0,698 | -0,519 | n. def. | 1 | 1 | 1 | 1,431 | 0,517 | n. def. | 1 | 1 | 1 | 1,218 | 0,285 | n. def. | 1 | 0 | 0 | NA | 2,959 |
| 12,796 | 3,678 | n. def. | 1 | 5 | 3 | 19,111 | 4,256 | 24,01 | 4 | 7 | 4 | 24,361 | 4,606 | 58,96 | 4 | 6 | 3 | 24,181 | 4,596 | 245,44 | 3 | 8 | 6 | 5,977 | 2,579 |
| 11,172 | 3,482 | 39,16 | 4 | 6 | 6 | 7,460 | 2,899 | 42,60 | 8 | 6 | 6 | 11,213 | 3,487 | 59,55 | 7 | 8 | 8 | 32,216 | 5,010 | 348,31 | 7 | 0 | 0 | NA | 0,776 |
| 15,879 | 3,989 | 76,84 | 3 | 1 | 1 | 5,663 | 2,502 | n. def. | 1 | 1 | 1 | 3,534 | 1,821 | n. def. | 1 | 2 | 2 | NA | 3,151 | n. def. | 0 | 0 | 0 | NA | 1,009 |
| 14,022 | 3,810 | 134,43 | 4 | 0 | 0 | NA | 1,225 | n. def. | 0 | 0 | 0 | NA | 2,799 | n. def. | 0 | 0 | 0 | NA | 1,291 | n. def. | 0 | 4 | 4 | 7,041 | 2,816 |
| 10,718 | 3,422 | 59,26 | 6 | 0 | 0 | NA | 2,928 | n. def. | 0 | 0 | 0 | NA | 2,960 | n. def. | 0 | 0 | 0 | NA | 2,625 | n. def. | 0 | 1 | 1 | 36,196 | 5,178 |
| 10,566 | 3,401 | 144,16 | 3 | 0 | 0 | NA | 0,757 | n. def. | 0 | 0 | 0 | NA | 1,603 | n. def. | 0 | 0 | 0 | NA | 4,900 | n. def. | 0 | 1 | 1 | 1,068 | 0,094 |
| 14,385 | 3,846 | 37,40 | 2 | 1 | 1 | 1,116 | 0,158 | n. def. | 1 | 1 | 1 | 2,318 | 1,213 | n. def. | 1 | 0 | 0 | NA | 3,931 | n. def. | 0 | 0 | 0 | NA | 2,201 |
| 23,065 | 4,528 | 22,10 | 5 | 4 | 4 | 0,913 | -0,132 | 22,64 | 4 | 2 | 2 | 2,451 | 1,293 | 35,92 | 2 | 5 | 5 | 9,602 | 3,263 | 85,09 | 4 | 1 | 1 | 1,386 | 0,471 |
| 7,648 | 2,935 | 25,38 | 2 | 1 | 1 | 0,174 | -2,526 | n. def. | 1 | 1 | 1 | NA | 4,586 | n. def. | 0 | 1 | 1 | 10,176 | 3,347 | n. def. | 1 | 1 | 1 | 27,060 | 4,758 |
| 5,892 | 2,559 | 176,48 | 2 | 0 | 0 | NA | 2,385 | n. def. | 0 | 0 | 0 | NA | 1,299 | n. def. | 0 | 0 | 0 | NA | 2,467 | n. def. | 0 | 1 | 1 | 1,057 | 0,079 |
| 8,828 | 3,142 | 49,22 | 2 | 3 | 3 | 8,174 | 3,031 | 46,24 | 2 | 2 | 2 | 13,187 | 3,721 | 40,16 | 3 | 3 | 3 | 62,913 | 5,975 | 57,38 | 2 | 0 | 0 | NA | 1,930 |
| 7,253 | 2,859 | 91,81 | 8 | 5 | 5 | 9,597 | 3,263 | 147,89 | 2 | 6 | 6 | 2,727 | 1,447 | 122,19 | 7 | 6 | 6 | 15,033 | 3,910 | 95,91 | 5 | 3 | 3 | 20,562 | 4,362 |
| 6,161 | 2,623 | n. def. | 1 | 0 | 0 | NA | 2,425 | n. def. | 0 | 0 | 0 | NA | 0,476 | n. def. | 0 | 1 | 1 | 5,303 | 2,407 | n. def. | 1 | 1 | 1 | 4,385 | 2,133 |
| 7,857 | 2,974 | 118,98 | 6 | 4 | 4 | 19,234 | 4,266 | 22,40 | 5 | 4 | 4 | 23,950 | 4,582 | 86,17 | 4 | 6 | 6 | 6,672 | 2,738 | 10,09 | 4 | 1 | 1 | 15,246 | 3,930 |
| 30,673 | 4,939 | 163,37 | 2 | 4 | 4 | 3,970 | 1,989 | 114,12 | 4 | 3 | 3 | 2,259 | 1,176 | 16,81 | 3 | 3 | 3 | 10,644 | 3,412 | 54,26 | 2 | 6 | 6 | 12,509 | 3,645 |
| 3,022 | 1,596 | 346,18 | 2 | 3 | 3 | 38,794 | 5,278 | 47,16 | 3 | 1 | 1 | 14,179 | 3,826 | 143,89 | 2 | 2 | 2 | 17,309 | 4,113 | 34,44 | 3 | 0 | 0 | NA | 3,076 |
| 19,721 | 4,302 | 32,61 | 12 | 4 | 4 | 0,856 | -0,225 | 8,75 | 4 | 6 | 6 | 2,322 | 1,216 | 25,53 | 6 | 1 | 1 | 7,802 | 2,964 | n. def. | 1 | 2 | 2 | 2,422 | 1,276 |
| 6,716 | 2,748 | n. def. | 1 | 2 | 2 | 12,991 | 3,699 | n. def. | 1 | 1 | 1 | 7,620 | 2,930 | n. def. | 1 | 2 | 2 | 4,218 | 2,077 | 76,51 | 3 | 0 | 0 | NA | 4,263 |
| 7,601 | 2,926 | 106,66 | 3 | 4 | 4 | 21,790 | 4,446 | 22,33 | 4 | 3 | 3 | 7,248 | 2,858 | n. def. | 1 | 4 | 4 | 8,958 | 3,163 | 129,60 | 2 | 0 | 0 | NA | 0,658 |
| 13,632 | 3,769 | n. def. | 1 | 1 | 1 | 4,551 | 2,186 | n. def. | 1 | 0 | 0 | NA | -0,240 | n. def. | 0 | 0 | 0 | NA | 2,303 | n. def. | 0 | 3 | 3 | 29,254 | 4,871 |
| 15,317 | 3,937 | 33,48 | 2 | 2 | 2 | 4,922 | 2,299 | 38,67 | 3 | 2 | 2 | 6,121 | 2,614 | 142,25 | 2 | 2 | 2 | 16,619 | 4,055 | 81,21 | 2 | 2 | 2 | 1,387 | 0,472 |
| 7,704 | 2,946 | 108,37 | 5 | 1 | 1 | 1,930 | 0,948 | n. def. | 1 | 0 | 0 | NA | 2,570 | n. def. | 0 | 1 | 1 | 1,674 | 0,743 | n. def. | 1 | 8 | 8 | 0,844 | -0,245 |
| 8,672 | 3,116 | 47,37 | 13 | 10 | 10 | 1,743 | 0,801 | 8,24 | 10 | 5 | 5 | 3,693 | 1,885 | 23,98 | 5 | 11 | 11 | 12,596 | 3,655 | 57,06 | 10 | 36 | 36 | 37,574 | 5,232 |
| 7,705 | 2,946 | 30,95 | 5 | 0 | 0 | NA | 3,215 | n. def. | 0 | 0 | 0 | NA | 1,542 | n. def. | 0 | 0 | 0 | NA | 2,992 | n. def. | 0 | 4 | 4 | 9,284 | 3,215 |
| 12,202 | 3,609 | 18,91 | 2 | 0 | 0 | NA | -0,160 | n. def. | 0 | 1 | 1 | 9,691 | 3,277 | n. def. | 1 | 1 | 1 | 18,386 | 4,201 | n. def. | 1 | 2 | 2 | 0,918 | -0,123 |
| 9,878 | 3,304 | n. def. | 1 | 1 | 1 | 5,203 | 2,379 | n. def. | 1 | 1 | 1 | 15,417 | 3,946 | n. def. | 1 | 1 | 1 | 3,764 | 1,912 | 135,72 | 2 | 5 | 5 | 0,958 | -0,061 |
| 15,648 | 3,968 | 63,22 | 36 | 18 | 18 | 0,720 | -0,474 | 9,98 | 21 | 13 | 13 | 1,564 | 0,645 | 20,25 | 16 | 12 | 12 | 7,825 | 2,968 | 47,86 | 14 | 2 | 2 | 1,100 | 0,138 |
| 13,031 | 3,704 | 65,24 | 27 | 19 | 19 | 0,840 | -0,252 | 15,01 | 22 | 14 | 14 | 1,842 | 0,882 | 56,01 | 18 | 12 | 12 | 9,236 | 3,207 | 46,84 | 15 | 6 | 6 | 0,817 | -0,292 |
| 11,710 | 3,550 | n. def. | 1 | 0 | 0 | NA | 2,752 | n. def. | 0 | 0 | 0 | NA | 2,118 | n. def. | 0 | 0 | 0 | NA | 1,352 | n. def. | 0 | 1 | 1 | 2,620 | 1,390 |
| 9,569 | 3,258 | 51,57 | 16 | 10 | 10 | 3,108 | 1,636 | 28,70 | 12 | 11 | 11 | 5,062 | 2,340 | 50,98 | 15 | 20 | 20 | 28,025 | 4,809 | 120,13 | 21 | 0 | 0 | NA | 1,144 |
| 6,995 | 2,806 | 148,95 | 5 | 0 | 0 | NA | -0,242 | n. def. | 0 | 0 | 0 | NA | 1,445 | n. def. | 0 | 0 | 0 | NA | 3,199 | n. def. | 0 | 17 | 17 | 13,702 | 3,776 |
| 10,079 | 3,333 | n. def. | 1 | 0 | 0 | NA | 3,394 | n. def. | 0 | 0 | 0 | NA | 2,832 | n. def. | 0 | 0 | 0 | NA | 3,714 | n. def. | 0 | 0 | 0 | NA | 3,052 |
| 7,711 | 2,947 | 36,46 | 2 | 6 | 6 | 14,041 | 3,812 | 33,66 | 5 | 6 | 6 | 7,309 | 2,870 | 129,41 | 5 | 5 | 5 | 1,425 | 0,511 | 256,41 | 2 | 1 | 1 | NA | 2,349 |
| NA | 4,340 | n. def. | 0 | 2 | 2 | 1,572 | 0,653 | 140,50 | 2 | 1 | 1 | 0,372 | -1,427 | n. def. | 1 | 0 | 0 | NA | 2,890 | n. def. | 0 | 0 | 0 | NA | 2,410 |
| 5,894 | 2,559 | n. def. | 1 | 0 | 0 | NA | 0,749 | n. def. | 0 | 0 | 0 | NA | 2,953 | n. def. | 0 | 0 | 0 | NA | 3,285 | n. def. | 0 | 1 | 1 | 3,502 | 1,808 |
| 6,986 | 2,805 | 24,06 | 20 | 9 | 9 | 0,448 | -1,158 | 13,40 | 9 | 7 | 7 | 1,262 | 0,336 | 23,61 | 7 | 2 | 2 | 8,373 | 3,066 | 52,64 | 3 | 22 | 22 | 0,985 | -0,021 |
| 11,168 | 3,481 | n. def. | 1 | 1 | 1 | NA | -0,075 | n. def. | 0 | 0 | 0 | NA | 1,835 | n. def. | 0 | 0 | 0 | NA | 3,585 | n. def. | 0 | 0 | 0 | NA | 1,597 |
| NA | 1,255 | n. def. | 0 | 1 | 1 | 3,573 | 1,837 | n. def. | 1 | 0 | 0 | NA | 4,138 | n. def. | 0 | 2 | 2 | 64,371 | 6,008 | n. def. | 1 | 0 | 0 | NA | 1,186 |
| 10,336 | 3,370 | 48,61 | 5 | 1 | 1 | 2,883 | 1,528 | n. def. | 1 | 1 | 1 | 6,603 | 2,723 | n. def. | 1 | 4 | 4 | 6,288 | 2,653 | 74,30 | 3 | 15 | 15 | 0,905 | -0,144 |
| 8,681 | 3,118 | 113,13 | 10 | 0 | 0 | NA | -1,299 | n. def. | 0 | 1 | 1 | 4,777 | 2,256 | n. def. | 1 | 2 | 2 | 10,149 | 3,343 | 46,03 | 2 | 2 | 2 | 1,170 | 0,226 |
| 15,927 | 3,993 | n. def. | 1 | 4 | 4 | 1,672 | 0,742 | 56,36 | 4 | 4 | 4 | 7,004 | 2,808 | 90,82 | 4 | 5 | 5 | 18,220 | 4,187 | 79,91 | 4 | 1 | 1 | 2,193 | 1,133 |
| 12,674 | 3,664 | 55,73 | 5 | 0 | 0 | NA | 2,607 | n. def. | 0 | 0 | 0 | NA | 1,266 | n. def. | 0 | 0 | 0 | NA | 2,467 | n. def. | 0 | 1 | 1 | 1,173 | 0,230 |
| 43,956 | 5,458 | n. def. | 1 | 1 | 1 | 10,826 | 3,436 | n. def. | 1 | 1 | 1 | 21,455 | 4,423 | n. def. | 1 | 0 | 0 | NA | 2,662 | n. def. | 0 | 4 | 4 | 92,980 | 6,539 |
| NA | 2,346 | n. def. | 0 | 3 | 3 | 3,047 | 1,607 | 31,33 | 2 | 1 | 1 | 7,250 | 2,858 | n. def. | 1 | 2 | 2 | 11,811 | 3,562 | n. def. | 1 | 10 | 10 | 3,594 | 1,845 |
| NA | 3,577 | n. def. | 0 | 0 | 0 | NA | 1,569 | n. def. | 0 | 0 | 0 | NA | 2,096 | n. def. | 0 | 0 | 0 | NA | 3,891 | n. def. | 0 | 3 | 3 | 10,853 | 3,440 |
| 8,333 | 3,059 | 26,48 | 3 | 1 | 1 | 4,222 | 2,078 | n. def. | 1 | 0 | 0 | NA | 2,701 | n. def. | 0 | 2 | 2 | 8,568 | 3,099 | 122,80 | 2 | 14 | 14 | 59,102 | 5,885 |
| 9,944 | 3,314 | 96,16 | 6 | 13 | 11 | 8,013 | 3,002 | 78,72 | 14 | 9 | 6 | 14,903 | 3,898 | 67,58 | 7 | 9 | 7 | 45,939 | 5,522 | 30,79 | 5 | 0 | 0 | NA | 0,098 |
| 6,938 | 2,795 | 23,72 | 2 | 0 | 0 | NA | 2,171 | n. def. | 0 | 1 | 1 | NA | 2,094 | n. def. | 0 | 2 | 2 | NA | 3,204 | n. def. | 0 | 1 | 1 | NA | 0,811 |
| NA | 1,996 | n. def. | 0 | 3 | 3 | 1,317 | 0,397 | 152,13 | 3 | 1 | 1 | 3,720 | 1,895 | n. def. | 1 | 3 | 3 | 34,626 | 5,114 | n. def. | 1 | 0 | 0 | NA | 3,292 |
| 4,316 | 2,110 | 119,81 | 2 | 2 | 2 | 5,714 | 2,515 | 56,66 | 2 | 3 | 3 | 3,231 | 1,692 | n. def. | 1 | 4 | 4 | 30,697 | 4,940 | n. def. | 1 | 0 | 0 | NA | 1,751 |
| 8,642 | 3,111 | n. def. | 1 | 3 | 3 | 18,904 | 4,241 | 24,84 | 3 | 2 | 2 | 45,000 | 5,492 | n. def. | 1 | 3 | 3 | 23,013 | 4,524 | n. def. | 1 | 7 | 7 | 6,324 | 2,661 |
| 7,828 | 2,969 | n. def. | 1 | 3 | 3 | 42,497 | 5,409 | 52,23 | 3 | 2 | 2 | 25,150 | 4,652 | 14,52 | 2 | 5 | 5 | 2,516 | 1,331 | 165,22 | 5 | 0 | 0 | NA | 2,496 |
| NA | 3,405 | n. def. | 0 | 0 | 0 | NA | -0,306 | n. def. | 0 | 0 | 0 | NA | 2,075 | n. def. | 0 | 0 | 0 | NA | 2,193 | n. def. | 0 | 1 | 1 | 3,672 | 1,876 |
| 12,402 | 3,632 | 48,68 | 2 | 1 | 1 | NA | 1,491 | n. def. | 0 | 3 | 3 | 10,318 | 3,367 | 110,45 | 3 | 6 | 6 | 10,080 | 3,333 | 129,30 | 3 | 0 | 0 | NA | 1,712 |
| 3,357 | 1,747 | 202,51 | 5 | 7 | 6 | 14,713 | 3,879 | 32,18 | 9 | 9 | 7 | 15,206 | 3,927 | 72,97 | 8 | 12 | 9 | 7,519 | 2,911 | 158,81 | 10 | 14 | 11 | 35,739 | 5,159 |
| 4,458 | 2,156 | 24,52 | 2 | 2 | 2 | 4,227 | 2,080 | 5,06 | 2 | 2 | 2 | 3,515 | 1,814 | 2,16 | 2 | 2 | 2 | 3,315 | 1,729 | 49,42 | 2 | 0 | 0 | NA | 1,280 |
| 5,116 | 2,355 | 15,08 | 2 | 0 | 0 | NA | 4,922 | n. def. | 0 | 0 | 0 | NA | 1,831 | n. def. | 0 | 1 | 1 | NA | 2,438 | n. def. | 0 | 2 | 2 | 5,333 | 2,415 |
| 6,455 | 2,690 | 36,58 | 2 | 0 | 0 | NA | 0,867 | n. def. | 0 | 0 | 0 | NA | 2,190 | n. def. | 0 | 0 | 0 | NA | 2,616 | n. def. | 0 | 2 | 2 | 1,037 | 0,052 |
| 4,970 | 2,313 | 8,89 | 3 | 2 | 2 | 1,941 | 0,957 | 6,33 | 2 | 3 | 3 | 2,274 | 1,185 | 11,82 | 3 | 1 | 1 | NA | 3,022 | n. def. | 0 | 0 | 0 | NA | 0,535 |
| 5,821 | 2,541 | 98,88 | 6 | 0 | 0 | NA | -1,218 | n. def. | 0 | 0 | 0 | NA | 1,115 | n. def. | 0 | 0 | 0 | NA | 3,127 | n. def. | 0 | 0 | 0 | NA | 5,719 |
| 3,794 | 1,924 | 22,04 | 3 | 1 | 1 | 1,618 | 0,695 | n. def. | 1 | 1 | 1 | 2,069 | 1,049 | n. def. | 1 | 2 | 2 | 8,696 | 3,120 | n. def. | 1 | 0 | 0 | NA | 3,659 |
| 7,306 | 2,869 | 20,35 | 4 | 1 | 1 | 0,634 | -0,656 | n. def. | 1 | 1 | 1 | 2,594 | 1,375 | n. def. | 1 | 2 | 2 | 8,572 | 3,100 | 127,67 | 2 | 1 | 1 | NA | 1,225 |
| 5,865 | 2,552 | 24,87 | 17 | 21 | 21 | 1,911 | 0,935 | 11,30 | 25 | 23 | 23 | 3,323 | 1,733 | 22,34 | 27 | 25 | 25 | 14,681 | 3,876 | 49,07 | 29 | 3 | 3 | 1,593 | 0,671 |
| 7,847 | 2,972 | n. def. | 1 | 1 | 0 | NA | 0,596 | n. def. | 0 | 3 | 1 | NA | 0,160 | n. def. | 0 | 3 | 0 | NA | 3,463 | n. def. | 0 | 6 | 3 | 6,114 | 2,612 |
| 5,405 | 2,434 | 37,43 | 9 | 12 | 12 | 1,823 | 0,866 | 24,32 | 14 | 7 | 7 | 3,147 | 1,654 | 46,06 | 9 | 7 | 7 | 11,201 | 3,486 | 32,89 | 5 | 0 | 0 | NA | 0,281 |
| 6,333 | 2,663 | 29,17 | 21 | 25 | 25 | 1,861 | 0,896 | 13,67 | 29 | 23 | 23 | 3,208 | 1,682 | 32,01 | 28 | 19 | 19 | 15,878 | 3,989 | 37,16 | 23 | 1 | 1 | 1,519 | 0,603 |
| 6,563 | 2,714 | 79,82 | 10 | 8 | 7 | 0,889 | -0,170 | 12,91 | 7 | 7 | 7 | 2,060 | 1,043 | 27,25 | 7 | 4 | 4 | 11,138 | 3,477 | 105,65 | 5 | 3 | 3 | 0,897 | -0,157 |
| NA | 2,273 | n. def. | 0 | 0 | 0 | NA | -3,341 | n. def. | 0 | 0 | 0 | NA | 3,127 | n. def. | 0 | 2 | 2 | 2,043 | 1,031 | n. def. | 1 | 1 | 1 | NA | 1,212 |
| NA | 2,314 | n. def. | 0 | 4 | 4 | 1,685 | 0,753 | 13,12 | 4 | 3 | 3 | 2,966 | 1,569 | 10,94 | 3 | 4 | 4 | 11,915 | 3,575 | 20,76 | 5 | 1 | 1 | 1,426 | 0,512 |
| 7,996 | 2,999 | 150,21 | 2 | 1 | 1 | 9,131 | 3,191 | n. def. | 1 | 0 | 0 | NA | 0,614 | n. def. | 0 | 1 | 1 | 9,329 | 3,222 | n. def. | 1 | 0 | 0 | NA | 2,748 |
| 3,135 | 1,648 | 32,50 | 2 | 1 | 1 | 6,708 | 2,746 | n. def. | 1 | 1 | 1 | 4,332 | 2,115 | n. def. | 1 | 1 | 1 | 4,554 | 2,187 | n. def. | 1 | 0 | 0 | NA | -0,458 |
| 5,029 | 2,330 | 58,39 | 2 | 1 | 1 | 0,999 | -0,002 | n. def. | 1 | 1 | 1 | 1,928 | 0,947 | n. def. | 1 | 0 | 0 | NA | 3,035 | n. def. | 0 | 0 | 0 | NA | 2,236 |
| 7,731 | 2,951 | n. def. | 1 | 2 | 2 | 2,156 | 1,108 | 22,32 | 2 | 2 | 2 | 3,377 | 1,756 | 3,98 | 2 | 1 | 1 | 14,189 | 3,827 | 60,06 | 2 | 3 | 3 | 2,550 | 1,351 |
| NA | 3,380 | n. def. | 0 | 3 | 3 | 0,666 | -0,587 | 27,76 | 4 | 2 | 2 | 1,780 | 0,832 | 25,30 | 3 | 1 | 1 | 10,205 | 3,351 | n. def. | 1 | 3 | 3 | 1,077 | 0,107 |
| 4,726 | 2,241 | 58,50 | 24 | 26 | 26 | 0,710 | -0,495 | 23,69 | 28 | 25 | 25 | 1,215 | 0,281 | 16,19 | 27 | 21 | 21 | 8,296 | 3,052 | 48,02 | 23 | 10 | 10 | 1,309 | 0,388 |
| 3,660 | 1,872 | 16,46 | 4 | 3 | 3 | 1,423 | 0,509 | 15,02 | 3 | 3 | 3 | 2,562 | 1,357 | 5,08 | 3 | 4 | 4 | 11,874 | 3,570 | 36,26 | 4 | 0 | 0 | NA | 3,197 |
| 4,832 | 2,273 | 17,03 | 9 | 10 | 10 | 1,214 | 0,280 | 18,95 | 11 | 9 | 9 | 1,894 | 0,921 | 30,27 | 10 | 8 | 8 | 6,710 | 2,746 | 91,72 | 8 | 0 | 0 | NA | 3,850 |
| NA | 2,035 | n. def. | 0 | 0 | 0 | NA | 0,036 | n. def. | 0 | 0 | 0 | NA | 1,670 | n. def. | 0 | 2 | 2 | 2,384 | 1,253 | n. def. | 1 | 1 | 1 | NA | 2,139 |
| 4,363 | 2,125 | 6,72 | 2 | 2 | 2 | 1,824 | 0,867 | 25,04 | 2 | 2 | 2 | 3,542 | 1,825 | 45,25 | 2 | 3 | 3 | 7,631 | 2,932 | 129,39 | 2 | 0 | 0 | NA | 2,534 |
| NA | 0,765 | n. def. | 0 | 1 | 1 | 1,542 | 0,624 | n. def. | 1 | 1 | 1 | 3,050 | 1,609 | n. def. | 1 | 4 | 4 | 4,700 | 2,233 | 26,61 | 4 | 1 | 1 | 1,485 | 0,570 |
| NA | 3,398 | n. def. | 0 | 1 | 1 | 3,893 | 1,961 | 99,60 | 2 | 2 | 2 | 5,399 | 2,433 | 60,42 | 2 | 2 | 2 | 67,155 | 6,069 | 221,80 | 4 | 5 | 5 | 131,643 | 7,040 |
| 4,746 | 2,247 | 13,09 | 5 | 10 | 10 | 1,168 | 0,224 | 13,63 | 10 | 8 | 8 | 2,675 | 1,419 | 32,70 | 8 | 11 | 11 | 9,472 | 3,244 | 74,08 | 13 | 0 | 0 | NA | 2,275 |
| 3,152 | 1,656 | n. def. | 1 | 0 | 0 | NA | 2,935 | n. def. | 0 | 0 | 0 | NA | 1,065 | n. def. | 0 | 0 | 0 | NA | 3,207 | n. def. | 0 | 2 | 2 | 5,383 | 2,428 |
| 5,807 | 2,538 | 34,75 | 5 | 2 | 2 | 4,446 | 2,153 | 0,53 | 2 | 3 | 3 | 5,281 | 2,401 | 4,67 | 3 | 4 | 4 | 7,422 | 2,892 | 30,95 | 6 | 0 | 0 | NA | 0,928 |
| NA | 1,140 | n. def. | 0 | 1 | 0 | NA | 1,710 | n. def. | 0 | 2 | 2 | 39,532 | 5,305 | n. def. | 1 | 2 | 2 | 22,230 | 4,474 | 40,71 | 2 | 3 | 2 | 169,690 | 7,407 |
| 2,312 | 1,209 | n. def. | 1 | 1 | 1 | 1,069 | 0,097 | n. def. | 1 | 1 | 1 | 2,021 | 1,015 | n. def. | 1 | 0 | 0 | NA | 3,171 | n. def. | 0 | 0 | 0 | NA | 5,595 |
| 4,962 | 2,311 | n. def. | 1 | 3 | 3 | 1,541 | 0,624 | 7,62 | 3 | 2 | 2 | 2,788 | 1,479 | 1,21 | 2 | 3 | 3 | 8,179 | 3,032 | 19,20 | 3 | 0 | 0 | NA | 2,743 |
| 2,201 | 1,138 | n. def. | 1 | 0 | 0 | NA | -1,981 | n. def. | 0 | 1 | 1 | 4,611 | 2,205 | n. def. | 1 | 1 | 1 | 55,310 | 5,789 | n. def. | 1 | 1 | 1 | 1,482 | 0,567 |
| 4,690 | 2,230 | 27,46 | 7 | 3 | 3 | 1,446 | 0,532 | 16,23 | 3 | 2 | 2 | 2,091 | 1,065 | 14,89 | 2 | 2 | 2 | 11,267 | 3,494 | 83,60 | 3 | 0 | 0 | NA | 2,642 |
| 5,646 | 2,497 | n. def. | 1 | 0 | 0 | NA | -0,497 | n. def. | 0 | 0 | 0 | NA | 3,523 | n. def. | 0 | 2 | 2 | 0,275 | -1,860 | n. def. | 1 | 0 | 0 | NA | 1,307 |
| 7,307 | 2,869 | 14,53 | 8 | 4 | 4 | 0,764 | -0,389 | 20,99 | 4 | 5 | 5 | 1,596 | 0,674 | 20,01 | 5 | 4 | 4 | 5,782 | 2,532 | 197,39 | 3 | 0 | 0 | NA | 1,405 |
| 3,738 | 1,902 | 19,17 | 12 | 12 | 12 | 1,502 | 0,586 | 11,66 | 15 | 10 | 10 | 2,495 | 1,319 | 24,69 | 13 | 13 | 13 | 8,041 | 3,007 | 100,75 | 13 | 1 | 1 | 1,033 | 0,047 |
| NA | 1,674 | n. def. | 0 | 1 | 1 | NA | 1,690 | n. def. | 0 | 2 | 2 | 9,797 | 3,292 | n. def. | 1 | 1 | 1 | 2,370 | 1,245 | n. def. | 1 | 12 | 12 | 22,942 | 4,520 |
| 4,231 | 2,081 | 25,55 | 2 | 5 | 5 | 1,760 | 0,815 | 6,56 | 5 | 3 | 3 | 3,698 | 1,887 | 1,84 | 3 | 3 | 3 | 9,083 | 3,183 | 83,63 | 4 | 0 | 0 | NA | 2,810 |
| 4,088 | 2,032 | 48,81 | 5 | 4 | 4 | 1,938 | 0,955 | 23,11 | 4 | 4 | 4 | 2,963 | 1,567 | 30,14 | 4 | 4 | 4 | 8,672 | 3,116 | 108,18 | 4 | 0 | 0 | NA | -0,037 |
| NA | 1,416 | n. def. | 0 | 1 | 1 | 24,726 | 4,628 | n. def. | 1 | 1 | 1 | 28,860 | 4,851 | n. def. | 1 | 2 | 2 | 55,475 | 5,794 | n. def. | 1 | 2 | 2 | 40,106 | 5,326 |
| 7,431 | 2,893 | 28,48 | 13 | 11 | 11 | 1,026 | 0,037 | 15,36 | 12 | 11 | 11 | 1,854 | 0,891 | 20,81 | 11 | 9 | 9 | 7,817 | 2,967 | 61,69 | 9 | 3 | 3 | 0,965 | -0,052 |
| 4,296 | 2,103 | 12,78 | 2 | 1 | 1 | 0,829 | -0,271 | n. def. | 1 | 1 | 1 | 2,042 | 1,030 | n. def. | 1 | 1 | 1 | NA | 3,075 | n. def. | 0 | 0 | 0 | NA | -0,036 |
| 6,111 | 2,611 | 4,59 | 2 | 1 | 1 | 1,503 | 0,588 | n. def. | 1 | 2 | 2 | 3,980 | 1,993 | 13,18 | 2 | 4 | 4 | 21,549 | 4,430 | 63,93 | 3 | 0 | 0 | NA | 1,340 |
| 2,949 | 1,560 | n. def. | 1 | 1 | 1 | 0,967 | -0,049 | n. def. | 1 | 1 | 1 | 1,337 | 0,419 | n. def. | 1 | 1 | 1 | 3,681 | 1,880 | n. def. | 1 | 0 | 0 | NA | 1,424 |
| 3,395 | 1,764 | n. def. | 1 | 1 | 1 | 1,901 | 0,927 | n. def. | 1 | 0 | 0 | NA | 0,829 | n. def. | 0 | 2 | 2 | 9,230 | 3,206 | 20,58 | 3 | 0 | 0 | NA | 0,686 |
| 3,787 | 1,921 | 18,27 | 8 | 5 | 5 | 0,923 | -0,116 | 5,09 | 5 | 3 | 3 | 1,816 | 0,861 | 27,51 | 3 | 2 | 2 | 3,879 | 1,956 | 161,14 | 2 | 0 | 0 | NA | 1,415 |
| 4,801 | 2,263 | 13,93 | 4 | 3 | 2 | 0,775 | -0,367 | 2,45 | 2 | 3 | 2 | 2,090 | 1,063 | 10,32 | 2 | 5 | 4 | 11,887 | 3,571 | 58,03 | 5 | 9 | 8 | 0,871 | -0,199 |
| 3,527 | 1,818 | 36,25 | 22 | 9 | 9 | 0,682 | -0,553 | 22,32 | 11 | 7 | 7 | 1,630 | 0,705 | 18,80 | 9 | 7 | 7 | 8,060 | 3,011 | 46,75 | 9 | 4 | 4 | 0,922 | -0,117 |
| 2,693 | 1,429 | 85,21 | 2 | 0 | 0 | NA | 0,002 | n. def. | 0 | 0 | 0 | NA | 1,581 | n. def. | 0 | 0 | 0 | NA | 2,589 | n. def. | 0 | 0 | 0 | NA | 1,368 |
| 2,941 | 1,556 | 31,37 | 7 | 1 | 1 | 1,401 | 0,487 | n. def. | 1 | 1 | 1 | 2,171 | 1,118 | n. def. | 1 | 1 | 1 | 27,484 | 4,781 | n. def. | 1 | 0 | 0 | NA | 0,532 |
| 3,208 | 1,682 | 2,22 | 2 | 1 | 1 | 1,110 | 0,150 | n. def. | 1 | 1 | 1 | 1,487 | 0,573 | n. def. | 1 | 2 | 2 | 9,133 | 3,191 | n. def. | 1 | 0 | 0 | NA | 5,251 |
| 3,099 | 1,632 | 21,26 | 11 | 3 | 3 | 1,072 | 0,101 | 2,18 | 3 | 1 | 1 | 1,193 | 0,255 | n. def. | 1 | 0 | 0 | NA | 2,217 | n. def. | 0 | 7 | 7 | 0,911 | -0,135 |
| 4,419 | 2,144 | 38,29 | 20 | 12 | 12 | 0,479 | -1,061 | 30,40 | 13 | 11 | 11 | 1,063 | 0,088 | 43,88 | 11 | 11 | 11 | 5,351 | 2,420 | 75,67 | 12 | 1 | 1 | 0,952 | -0,071 |
| 5,299 | 2,406 | 0,85 | 2 | 0 | 0 | NA | 1,848 | n. def. | 0 | 1 | 1 | 1,395 | 0,480 | n. def. | 1 | 2 | 2 | 8,519 | 3,091 | 12,59 | 3 | 1 | 1 | 0,972 | -0,041 |
| 4,422 | 2,145 | 28,81 | 48 | 27 | 27 | 0,404 | -1,307 | 13,50 | 36 | 25 | 25 | 1,056 | 0,078 | 26,85 | 34 | 19 | 19 | 4,973 | 2,314 | 41,75 | 24 | 7 | 7 | 1,156 | 0,209 |
| 3,367 | 1,751 | n. def. | 1 | 0 | 0 | NA | 3,232 | n. def. | 0 | 1 | 1 | 2,027 | 1,019 | n. def. | 1 | 1 | 1 | 4,669 | 2,223 | n. def. | 1 | 0 | 0 | NA | 1,051 |
| 3,940 | 1,978 | 74,39 | 4 | 4 | 4 | 0,430 | -1,218 | 17,60 | 3 | 3 | 3 | 0,891 | -0,166 | 41,78 | 3 | 2 | 2 | 4,702 | 2,233 | 143,83 | 2 | 1 | 1 | 0,962 | -0,055 |
| 3,121 | 1,642 | 8,92 | 2 | 2 | 2 | 1,162 | 0,217 | 19,28 | 2 | 2 | 2 | 1,683 | 0,751 | 23,61 | 2 | 2 | 2 | 4,264 | 2,092 | n. def. | 1 | 0 | 0 | NA | 3,231 |
| 3,068 | 1,617 | n. def. | 1 | 3 | 3 | 3,491 | 1,804 | 12,93 | 3 | 1 | 1 | 4,358 | 2,124 | n. def. | 1 | 5 | 5 | 14,669 | 3,875 | 146,81 | 4 | 0 | 0 | NA | -0,789 |
| 2,341 | 1,227 | 18,54 | 7 | 5 | 5 | 1,068 | 0,095 | 9,01 | 6 | 4 | 4 | 2,570 | 1,362 | 21,86 | 4 | 5 | 5 | 11,945 | 3,578 | 94,16 | 5 | 7 | 7 | 0,987 | -0,019 |
| 3,115 | 1,639 | 41,03 | 24 | 23 | 23 | 0,712 | -0,490 | 32,76 | 25 | 21 | 21 | 1,484 | 0,569 | 65,03 | 24 | 17 | 17 | 7,649 | 2,935 | 67,23 | 19 | 7 | 7 | 2,163 | 1,113 |
| 3,672 | 1,877 | 31,00 | 24 | 24 | 24 | 0,444 | -1,170 | 28,06 | 31 | 19 | 19 | 0,963 | -0,054 | 24,65 | 25 | 7 | 7 | 2,677 | 1,420 | 25,14 | 11 | 6 | 6 | 1,066 | 0,092 |
| 2,325 | 1,218 | 18,03 | 2 | 2 | 2 | 2,267 | 1,181 | 6,86 | 2 | 1 | 1 | 2,447 | 1,291 | n. def. | 1 | 5 | 5 | 3,093 | 1,629 | 140,10 | 4 | 0 | 0 | NA | 1,939 |
| NA | 1,108 | n. def. | 0 | 2 | 2 | 1,242 | 0,313 | 5,24 | 2 | 2 | 2 | 1,957 | 0,969 | 1,21 | 2 | 2 | 2 | 13,517 | 3,757 | 9,74 | 2 | 0 | 0 | NA | 1,261 |
| 2,530 | 1,339 | 33,70 | 37 | 28 | 27 | 0,876 | -0,192 | 41,70 | 43 | 31 | 30 | 1,531 | 0,614 | 77,53 | 46 | 23 | 22 | 6,225 | 2,638 | 31,45 | 31 | 14 | 14 | 1,443 | 0,529 |
| NA | 3,190 | n. def. | 0 | 0 | 0 | NA | -1,350 | n. def. | 0 | 0 | 0 | NA | 3,775 | n. def. | 0 | 0 | 0 | NA | 2,071 | n. def. | 0 | 1 | 1 | 13,742 | 3,781 |
| 2,056 | 1,040 | 10,58 | 5 | 0 | 0 | NA | 0,112 | n. def. | 0 | 0 | 0 | NA | 3,315 | n. def. | 0 | 0 | 0 | NA | 2,040 | n. def. | 0 | 0 | 0 | NA | 4,017 |
| 2,135 | 1,094 | 38,36 | 16 | 2 | 2 | 0,769 | -0,380 | 15,79 | 2 | 3 | 3 | 1,449 | 0,535 | 14,86 | 3 | 1 | 1 | 7,873 | 2,977 | n. def. | 1 | 0 | 0 | NA | 3,881 |
| 1,173 | 0,230 | 25,99 | 16 | 1 | 1 | 1,344 | 0,426 | n. def. | 1 | 1 | 1 | 1,866 | 0,900 | n. def. | 1 | 2 | 2 | 1,813 | 0,859 | 213,31 | 2 | 0 | 0 | NA | -2,347 |
| 2,086 | 1,061 | n. def. | 1 | 0 | 0 | NA | -2,011 | n. def. | 0 | 0 | 0 | NA | 1,556 | n. def. | 0 | 0 | 0 | NA | 3,969 | n. def. | 0 | 3 | 3 | 1,207 | 0,271 |
| 2,657 | 1,410 | 42,17 | 2 | 2 | 2 | 0,718 | -0,479 | 9,46 | 2 | 2 | 2 | 1,691 | 0,758 | 25,27 | 2 | 2 | 2 | 8,526 | 3,092 | 18,11 | 3 | 0 | 0 | NA | -0,821 |
| 2,647 | 1,404 | 39,98 | 16 | 6 | 6 | 0,246 | -2,023 | 17,06 | 6 | 5 | 5 | 0,751 | -0,414 | 29,65 | 6 | 2 | 2 | 2,818 | 1,494 | 35,93 | 2 | 0 | 0 | NA | 1,387 |
| 1,623 | 0,698 | 10,53 | 4 | 1 | 1 | 0,655 | -0,609 | n. def. | 1 | 2 | 2 | 1,332 | 0,414 | 3,22 | 2 | 0 | 0 | NA | 3,011 | n. def. | 0 | 0 | 0 | NA | 1,299 |
| 2,396 | 1,261 | 13,71 | 8 | 11 | 11 | 0,268 | -1,902 | 18,06 | 11 | 11 | 11 | 0,627 | -0,674 | 49,81 | 11 | 13 | 13 | 3,138 | 1,650 | 65,34 | 13 | 2 | 2 | 1,463 | 0,549 |
| 1,216 | 0,282 | 30,20 | 26 | 13 | 13 | 0,297 | -1,752 | 19,97 | 15 | 11 | 11 | 0,779 | -0,360 | 19,45 | 12 | 8 | 8 | 5,457 | 2,448 | 43,97 | 9 | 0 | 0 | NA | 1,952 |

| | | Cor1 - Replicate 2 | | | | | | Cor1 - Replicate 3 | | | | | |
|---|---|---|---|---|---|---|---|---|---|---|---|---|---|
| Ratio Variability [%] | Ratio Count | Peptides | Unique Peptides | Ratio Cor1/WT | log2 Ratio Cor1/WT | Ratio Variability [%] | Ratio Count | Peptides | Unique Peptides | Ratio Cor1/WT | log2 Ratio Cor1/WT | Ratio Variability [%] | Ratio Count |
| 89,40 | 22 | 13 | 13 | 15,531 | 3,957 | 33,27 | 34 | 14 | 14 | 15,078 | 3,914 | 57,58 | 30 |
| 102,48 | 138 | 51 | 51 | 15,155 | 3,922 | 85,06 | 375 | 52 | 52 | 14,008 | 3,808 | 93,33 | 311 |
| 123,67 | 185 | 60 | 60 | 18,100 | 4,178 | 101,08 | 458 | 66 | 66 | 15,167 | 3,923 | 115,10 | 436 |
| 252,87 | 21 | 8 | 8 | 77,610 | 6,278 | 187,20 | 29 | 8 | 8 | 38,518 | 5,267 | 205,32 | 30 |
| 103,87 | 12 | 15 | 15 | 9,316 | 3,220 | 74,12 | 27 | 11 | 11 | 8,372 | 3,066 | 52,33 | 22 |
| 54,42 | 4 | 3 | 3 | 24,258 | 4,600 | 78,27 | 6 | 2 | 2 | 28,287 | 4,822 | 21,30 | 4 |
| 141,87 | 16 | 6 | 6 | 424,737 | 8,730 | 164,13 | 22 | 8 | 8 | 196,406 | 7,618 | 162,83 | 18 |
| 31,77 | 2 | 1 | 1 | 3,403 | 1,767 | 45,82 | 4 | 2 | 2 | 3,982 | 1,993 | 5,25 | 2 |
| 162,63 | 40 | 13 | 13 | 84,360 | 6,398 | 152,45 | 96 | 15 | 15 | 85,215 | 6,413 | 160,85 | 75 |
| 120,77 | 3 | 11 | 11 | 7,902 | 2,982 | 37,96 | 18 | 3 | 3 | 8,269 | 3,048 | 16,28 | 4 |
| 161,09 | 28 | 12 | 12 | 118,756 | 6,892 | 187,94 | 68 | 11 | 11 | 78,309 | 6,291 | 179,82 | 55 |
| 233,42 | 27 | 13 | 13 | 52,676 | 5,719 | 197,88 | 57 | 12 | 12 | 47,923 | 5,583 | 187,79 | 42 |
| 180,82 | 49 | 23 | 21 | 62,294 | 5,961 | 151,26 | 80 | 28 | 25 | 41,305 | 5,368 | 129,72 | 63 |
| 161,49 | 19 | 12 | 12 | 394,867 | 8,625 | 156,93 | 43 | 12 | 12 | 186,979 | 7,547 | 154,93 | 33 |
| 77,52 | 22 | 9 | 8 | 0,696 | -0,523 | 35,07 | 36 | 9 | 8 | 0,779 | -0,361 | 34,47 | 29 |
| n. def. | 1 | 0 | 0 | NA | 3,638 | n. def. | 0 | 1 | 1 | 10,006 | 3,323 | n. def. | 1 |
| 165,45 | 24 | 9 | 9 | 163,924 | 7,357 | 168,41 | 38 | 9 | 9 | 43,225 | 5,434 | 181,49 | 35 |
| n. def. | 0 | 6 | 6 | 5,342 | 2,417 | 74,79 | 10 | 1 | 1 | 4,925 | 2,300 | n. def. | 1 |
| n. def. | 0 | 1 | 1 | 1,983 | 0,988 | 62,17 | 3 | 0 | 0 | NA | 2,202 | n. def. | 0 |
| 182,54 | 42 | 15 | 15 | 173,593 | 7,440 | 181,08 | 97 | 17 | 17 | 80,847 | 6,337 | 150,23 | 83 |
| 187,40 | 61 | 16 | 16 | 40,820 | 5,351 | 178,54 | 141 | 14 | 14 | 22,662 | 4,502 | 169,48 | 96 |
| 72,39 | 20 | 9 | 8 | 0,684 | -0,547 | 66,36 | 34 | 9 | 8 | 0,730 | -0,455 | 24,73 | 30 |
| n. def. | 0 | 0 | 0 | NA | 0,575 | n. def. | 0 | 0 | 0 | NA | 0,637 | n. def. | 0 |
| n. def. | 0 | 0 | 0 | NA | 0,947 | n. def. | 0 | 0 | 0 | NA | 2,247 | n. def. | 0 |
| 114,49 | 21 | 16 | 16 | 8,451 | 3,079 | 45,47 | 48 | 12 | 12 | 8,262 | 3,046 | 71,80 | 34 |
| n. def. | 0 | 0 | 0 | NA | 4,083 | n. def. | 0 | 0 | 0 | NA | -0,324 | n. def. | 0 |
| n. def. | 0 | 0 | 0 | NA | 0,700 | n. def. | 0 | 0 | 0 | NA | 1,235 | n. def. | 0 |
| 69,62 | 3 | 4 | 4 | 7,391 | 2,886 | 66,52 | 9 | 1 | 1 | 6,132 | 2,616 | n. def. | 1 |
| n. def. | 0 | 0 | 0 | NA | -2,093 | n. def. | 0 | 0 | 0 | NA | 4,262 | n. def. | 0 |
| 142,73 | 5 | 3 | 3 | 306,007 | 8,257 | 66,14 | 6 | 3 | 3 | 82,041 | 6,358 | 141,91 | 6 |
| n. def. | 0 | 0 | 0 | NA | 0,119 | n. def. | 0 | 0 | 0 | NA | 1,093 | n. def. | 0 |
| 166,37 | 12 | 8 | 8 | 20,326 | 4,345 | 55,61 | 20 | 10 | 10 | 16,201 | 4,018 | 92,60 | 14 |
| 5,97 | 3 | 6 | 6 | 2,693 | 1,429 | 19,60 | 15 | 5 | 5 | 3,244 | 1,698 | 24,27 | 9 |
| n. def. | 0 | 0 | 0 | NA | 1,689 | n. def. | 0 | 0 | 0 | NA | 2,133 | n. def. | 0 |
| 104,76 | 3 | 4 | 4 | 5,279 | 2,400 | 10,38 | 7 | 4 | 4 | 5,200 | 2,379 | 25,02 | 7 |
| n. def. | 0 | 0 | 0 | NA | 5,193 | n. def. | 0 | 0 | 0 | NA | 0,068 | n. def. | 0 |
| 186,35 | 7 | 7 | 5 | 91,785 | 6,520 | 251,05 | 16 | 6 | 3 | 78,487 | 6,294 | 268,06 | 8 |
| n. def. | 0 | 0 | 0 | NA | 2,326 | n. def. | 0 | 1 | 1 | NA | 2,828 | n. def. | 0 |
| n. def. | 0 | 0 | 0 | NA | 1,499 | n. def. | 0 | 0 | 0 | NA | 3,063 | n. def. | 0 |
| 37,80 | 4 | 13 | 13 | 8,267 | 3,047 | 93,11 | 16 | 7 | 7 | 8,293 | 3,052 | 109,73 | 11 |
| 0,14 | 3 | 1 | 1 | 40,165 | 5,328 | 67,21 | 3 | 1 | 1 | 38,040 | 5,249 | 8,76 | 3 |
| n. def. | 1 | 1 | 1 | 1,320 | 0,400 | n. def. | 1 | 0 | 0 | NA | 2,219 | n. def. | 0 |
| n. def. | 0 | 0 | 0 | NA | -0,715 | n. def. | 0 | 0 | 0 | NA | 2,384 | n. def. | 0 |
| 4,19 | 2 | 3 | 3 | 2,523 | 1,335 | 73,81 | 6 | 1 | 1 | 0,875 | -0,192 | n. def. | 1 |
| n. def. | 1 | 1 | 1 | 12,030 | 3,589 | 60,13 | 2 | 1 | 1 | 5,695 | 2,510 | n. def. | 1 |
| 19,87 | 2 | 1 | 1 | NA | 0,330 | n. def. | 0 | 1 | 1 | 0,982 | -0,026 | n. def. | 1 |
| n. def. | 0 | 1 | 1 | 2,074 | 1,053 | 8,99 | 2 | 0 | 0 | NA | 1,421 | n. def. | 0 |
| 43,60 | 5 | 3 | 3 | 21,100 | 4,399 | 81,88 | 9 | 3 | 3 | 29,631 | 4,889 | 76,01 | 9 |
| n. def. | 1 | 3 | 3 | 22,474 | 4,490 | 298,54 | 3 | 3 | 3 | 3,091 | 1,628 | n. def. | 1 |
| n. def. | 1 | 3 | 3 | 9,844 | 3,299 | n. def. | 1 | 1 | 1 | 8,214 | 3,038 | 63,20 | 2 |
| 162,42 | 3 | 11 | 11 | 13,194 | 3,722 | 126,98 | 16 | 7 | 7 | 4,413 | 2,142 | 117,26 | 4 |
| n. def. | 0 | 0 | 0 | NA | 1,680 | n. def. | 0 | 0 | 0 | NA | 4,202 | n. def. | 0 |
| 3,42 | 3 | 6 | 6 | 2,478 | 1,309 | 12,65 | 8 | 3 | 3 | 2,140 | 1,097 | 11,92 | 5 |
| n. def. | 0 | 0 | 0 | NA | 3,560 | n. def. | 0 | 0 | 0 | NA | 2,687 | n. def. | 0 |
| n. def. | 0 | 2 | 2 | 5,618 | 2,490 | n. def. | 1 | 0 | 0 | NA | 3,902 | n. def. | 0 |
| 54,71 | 2 | 3 | 3 | 150,709 | 7,236 | 21,72 | 2 | 4 | 4 | 40,196 | 5,329 | 140,88 | 2 |
| 151,30 | 4 | 3 | 3 | 0,919 | -0,122 | 135,85 | 7 | 2 | 2 | 2,965 | 1,568 | 122,45 | 7 |
| 76,18 | 14 | 11 | 11 | 0,882 | -0,180 | 20,45 | 39 | 7 | 7 | 0,867 | -0,206 | 14,29 | 19 |
| 139,78 | 49 | 40 | 40 | 51,948 | 5,699 | 127,36 | 101 | 42 | 42 | 41,874 | 5,388 | 123,52 | 69 |
| 44,89 | 7 | 7 | 7 | 9,811 | 3,294 | 172,72 | 18 | 5 | 5 | 4,473 | 2,161 | 215,22 | 7 |
| 80,77 | 7 | 5 | 5 | 0,898 | -0,155 | 36,44 | 15 | 2 | 2 | 0,968 | -0,047 | 9,02 | 12 |
| 66,38 | 12 | 4 | 4 | 0,935 | -0,098 | 31,03 | 19 | 4 | 4 | 1,038 | 0,054 | 8,82 | 12 |
| 25,04 | 2 | 0 | 0 | NA | -0,782 | n. def. | 0 | 1 | 1 | 0,949 | -0,075 | n. def. | 1 |
| 21,19 | 16 | 13 | 13 | 0,694 | -0,526 | 30,87 | 41 | 7 | 7 | 0,814 | -0,298 | 28,24 | 24 |
| n. def. | 1 | 2 | 2 | 2,649 | 1,405 | 26,14 | 4 | 0 | 0 | NA | -2,032 | n. def. | 0 |
| n. def. | 0 | 0 | 0 | NA | 1,664 | n. def. | 0 | 0 | 0 | NA | 1,944 | n. def. | 0 |
| 147,45 | 13 | 21 | 21 | 10,390 | 3,377 | 136,65 | 36 | 16 | 16 | 8,511 | 3,089 | 148,51 | 23 |
| n. def. | 0 | 1 | 1 | 1,060 | 0,084 | 3,36 | 2 | 1 | 1 | 1,061 | 0,086 | 12,58 | 2 |
| n. def. | 0 | 5 | 5 | 17,280 | 4,111 | 58,41 | 3 | 1 | 1 | NA | 2,459 | n. def. | 0 |
| n. def. | 0 | 0 | 0 | NA | 0,921 | n. def. | 0 | 0 | 0 | NA | 3,831 | n. def. | 0 |
| n. def. | 1 | 1 | 1 | 6,957 | 2,798 | 4,99 | 3 | 1 | 1 | 4,785 | 2,259 | 7,53 | 2 |
| 71,16 | 53 | 23 | 23 | 0,968 | -0,046 | 34,13 | 114 | 25 | 25 | 1,018 | 0,025 | 17,57 | 99 |
| n. def. | 0 | 1 | 1 | 0,723 | -0,468 | n. def. | 1 | 1 | 1 | 1,259 | 0,332 | n. def. | 1 |
| n. def. | 0 | 0 | 0 | NA | 1,938 | n. def. | 0 | 0 | 0 | NA | -2,804 | n. def. | 0 |
| 59,12 | 39 | 15 | 15 | 0,903 | -0,147 | 35,14 | 87 | 16 | 16 | 0,971 | -0,042 | 14,26 | 64 |
| 97,55 | 4 | 2 | 2 | 1,070 | 0,098 | 16,45 | 5 | 2 | 2 | 1,089 | 0,124 | 5,73 | 2 |
| n. def. | 1 | 1 | 1 | NA | -1,141 | n. def. | 0 | 2 | 2 | 4,798 | 2,262 | 29,05 | 4 |
| n. def. | 1 | 2 | 2 | 1,261 | 0,334 | 20,27 | 2 | 0 | 0 | NA | 1,741 | n. def. | 0 |
| 135,33 | 3 | 9 | 9 | 9,281 | 3,214 | 99,37 | 12 | 5 | 5 | 21,482 | 4,425 | 110,25 | 5 |
| 353,20 | 7 | 10 | 10 | 18,243 | 4,189 | 146,23 | 13 | 10 | 10 | 10,063 | 3,331 | 351,59 | 11 |
| 52,75 | 7 | 6 | 6 | 11,462 | 3,519 | 62,58 | 9 | 5 | 5 | 11,243 | 3,491 | 41,70 | 11 |
| 120,78 | 31 | 15 | 15 | 65,317 | 6,029 | 91,34 | 65 | 14 | 14 | 58,052 | 5,859 | 103,69 | 35 |
| n. def. | 0 | 0 | 0 | NA | -1,408 | n. def. | 0 | 1 | 0 | NA | 3,876 | n. def. | 0 |
| n. def. | 0 | 2 | 2 | 12,378 | 3,630 | n. def. | 1 | 2 | 2 | 6,671 | 2,738 | 81,21 | 2 |
| n. def. | 0 | 0 | 0 | NA | 2,125 | n. def. | 0 | 0 | 0 | NA | 2,718 | n. def. | 0 |
| n. def. | 0 | 0 | 0 | NA | -0,037 | n. def. | 0 | 0 | 0 | NA | 3,884 | n. def. | 0 |
| 291,58 | 9 | 7 | 7 | 8,949 | 3,162 | 213,53 | 16 | 8 | 8 | 16,020 | 4,002 | 268,73 | 14 |
| n. def. | 0 | 0 | 0 | NA | 1,527 | n. def. | 0 | 0 | 0 | NA | 3,067 | n. def. | 0 |
| n. def. | 1 | 1 | 1 | 5,025 | 2,329 | n. def. | 1 | 1 | 1 | 2,071 | 1,050 | n. def. | 1 |
| n. def. | 0 | 0 | 0 | NA | 2,221 | n. def. | 0 | 0 | 0 | NA | 4,053 | n. def. | 0 |
| 156,87 | 31 | 11 | 8 | 30,630 | 4,937 | 151,35 | 46 | 13 | 9 | 50,541 | 5,659 | 166,11 | 46 |
| n. def. | 0 | 0 | 0 | NA | 0,713 | n. def. | 0 | 0 | 0 | NA | 0,041 | n. def. | 0 |
| 129,91 | 2 | 5 | 5 | 7,130 | 2,834 | 69,57 | 2 | 5 | 5 | 5,154 | 2,366 | 48,30 | 6 |
| 95,56 | 4 | 5 | 5 | 1,071 | 0,098 | 20,52 | 12 | 2 | 2 | 0,932 | -0,102 | 21,17 | 3 |
| n. def. | 0 | 0 | 0 | NA | 2,849 | n. def. | 0 | 0 | 0 | NA | 1,112 | n. def. | 0 |
| n. def. | 0 | 1 | 1 | 4,179 | 2,063 | 5,96 | 2 | 1 | 1 | 3,445 | 1,784 | 7,66 | 2 |
| n. def. | 0 | 0 | 0 | NA | 0,778 | n. def. | 0 | 0 | 0 | NA | 1,929 | n. def. | 0 |
| n. def. | 0 | 1 | 1 | 4,592 | 2,199 | n. def. | 1 | 1 | 1 | NA | -0,289 | n. def. | 0 |
| 11,67 | 3 | 6 | 6 | 1,565 | 0,646 | 24,92 | 21 | 4 | 4 | 1,619 | 0,695 | 12,05 | 10 |
| 220,92 | 3 | 9 | 6 | 15,395 | 3,944 | 204,32 | 17 | 9 | 5 | 8,331 | 3,059 | 185,09 | 5 |
| n. def. | 0 | 0 | 0 | NA | 5,848 | n. def. | 0 | 0 | 0 | NA | 5,409 | n. def. | 0 |
| 13,21 | 2 | 4 | 4 | 1,374 | 0,458 | 21,56 | 13 | 3 | 3 | 1,514 | 0,598 | 17,70 | 10 |
| 11,48 | 6 | 8 | 8 | 0,895 | -0,161 | 23,94 | 17 | 6 | 6 | 0,994 | -0,009 | 62,76 | 9 |
| n. def. | 0 | 5 | 5 | 6,711 | 2,747 | 104,24 | 3 | 2 | 2 | 6,862 | 2,779 | n. def. | 1 |
| 17,59 | 2 | 3 | 3 | 1,585 | 0,664 | 19,62 | 8 | 1 | 1 | 1,574 | 0,654 | 12,32 | 3 |
| n. def. | 0 | 1 | 1 | NA | 4,199 | n. def. | 0 | 0 | 0 | NA | 3,222 | n. def. | 0 |
| n. def. | 0 | 0 | 0 | NA | 3,601 | n. def. | 0 | 0 | 0 | NA | 0,818 | n. def. | 0 |
| n. def. | 0 | 1 | 1 | 0,117 | -3,099 | 202,24 | 3 | 0 | 0 | NA | 2,570 | n. def. | 0 |
| 10,66 | 4 | 3 | 3 | 2,240 | 1,163 | 26,15 | 7 | 3 | 3 | 2,344 | 1,229 | 18,87 | 7 |
| 22,71 | 4 | 3 | 3 | 0,997 | -0,005 | 26,45 | 10 | 4 | 4 | 1,011 | 0,015 | 10,65 | 7 |
| 59,07 | 17 | 17 | 17 | 1,271 | 0,346 | 27,68 | 51 | 15 | 15 | 1,265 | 0,339 | 27,19 | 37 |
| n. def. | 0 | 0 | 0 | NA | 3,140 | n. def. | 0 | 0 | 0 | NA | 3,240 | n. def. | 0 |
| n. def. | 0 | 0 | 0 | NA | 3,385 | n. def. | 0 | 0 | 0 | NA | 0,892 | n. def. | 0 |
| n. def. | 0 | 4 | 4 | 2,426 | 1,278 | 365,58 | 2 | 4 | 4 | 7,742 | 2,953 | 120,96 | 3 |
| n. def. | 0 | 1 | 1 | NA | 3,380 | n. def. | 0 | 1 | 1 | NA | 3,358 | n. def. | 0 |
| 93,92 | 3 | 1 | 1 | 1,613 | 0,690 | n. def. | 1 | 1 | 1 | 1,376 | 0,460 | n. def. | 1 |
| 85,35 | 6 | 3 | 3 | 76,441 | 6,256 | 89,62 | 12 | 3 | 3 | 245,284 | 7,938 | 108,84 | 4 |
| n. def. | 0 | 0 | 0 | NA | 2,036 | n. def. | 0 | 0 | 0 | NA | 3,176 | n. def. | 0 |
| n. def. | 1 | 5 | 5 | 11,618 | 3,538 | 36,70 | 5 | 4 | 4 | 11,283 | 3,496 | 39,43 | 3 |
| n. def. | 0 | 0 | 0 | NA | 1,930 | n. def. | 0 | 0 | 0 | NA | 1,530 | n. def. | 0 |
| 52,15 | 5 | 5 | 3 | 169,503 | 7,405 | 219,08 | 17 | 4 | 3 | 206,646 | 7,691 | 169,25 | 12 |
| n. def. | 0 | 0 | 0 | NA | 0,855 | n. def. | 0 | 0 | 0 | NA | 0,843 | n. def. | 0 |
| n. def. | 0 | 0 | 0 | NA | 1,786 | n. def. | 0 | 0 | 0 | NA | 3,623 | n. def. | 0 |
| 1,30 | 2 | 3 | 3 | 1,331 | 0,412 | 20,91 | 5 | 2 | 2 | 1,430 | 0,516 | 6,91 | 3 |
| n. def. | 0 | 0 | 0 | NA | 1,198 | n. def. | 0 | 0 | 0 | NA | -2,458 | n. def. | 0 |
| n. def. | 0 | 2 | 2 | 7,759 | 2,956 | n. def. | 1 | 2 | 2 | 2,557 | 1,355 | n. def. | 1 |
| n. def. | 0 | 0 | 0 | NA | 5,754 | n. def. | 0 | 0 | 0 | NA | 1,290 | n. def. | 0 |
| n. def. | 1 | 1 | 1 | NA | 4,153 | n. def. | 0 | 1 | 1 | NA | 2,025 | n. def. | 0 |
| 132,77 | 13 | 8 | 8 | 17,643 | 4,141 | 134,39 | 22 | 12 | 12 | 52,138 | 5,704 | 117,32 | 17 |
| n. def. | 0 | 0 | 0 | NA | 4,326 | n. def. | 0 | 0 | 0 | NA | 1,138 | n. def. | 0 |
| n. def. | 0 | 0 | 0 | NA | 1,006 | n. def. | 0 | 0 | 0 | NA | 3,764 | n. def. | 0 |
| 183,64 | 6 | 2 | 2 | 18,427 | 4,204 | 200,84 | 11 | 2 | 2 | 74,102 | 6,211 | 186,46 | 12 |
| 18,89 | 3 | 4 | 4 | 1,043 | 0,060 | 22,22 | 15 | 4 | 4 | 1,161 | 0,216 | 14,23 | 12 |
| n. def. | 0 | 0 | 0 | NA | 0,681 | n. def. | 0 | 0 | 0 | NA | 2,813 | n. def. | 0 |
| n. def. | 0 | 0 | 0 | NA | -0,599 | n. def. | 0 | 0 | 0 | NA | -0,021 | n. def. | 0 |
| n. def. | 0 | 0 | 0 | NA | 2,085 | n. def. | 0 | 0 | 0 | NA | 0,672 | n. def. | 0 |
| n. def. | 0 | 1 | 1 | 1,467 | 0,553 | 23,21 | 2 | 1 | 1 | 1,504 | 0,589 | 5,89 | 3 |
| n. def. | 0 | 1 | 1 | NA | 0,859 | n. def. | 0 | 0 | 0 | NA | 3,172 | n. def. | 0 |
| 79,05 | 20 | 10 | 9 | 0,859 | -0,220 | 22,70 | 36 | 10 | 9 | 0,897 | -0,157 | 8,27 | 26 |
| 62,77 | 8 | 4 | 4 | 0,980 | -0,029 | 18,46 | 15 | 4 | 4 | 0,982 | -0,026 | 10,02 | 9 |
| n. def. | 0 | 7 | 7 | 4,692 | 2,230 | 124,06 | 9 | 2 | 2 | 14,551 | 3,863 | n. def. | 1 |
| n. def. | 0 | 1 | 1 | 0,736 | -0,443 | n. def. | 1 | 0 | 0 | NA | 2,681 | n. def. | 0 |
| n. def. | 0 | 0 | 0 | NA | 1,193 | n. def. | 0 | 0 | 0 | NA | 1,722 | n. def. | 0 |
| 68,11 | 12 | 12 | 12 | 0,954 | -0,067 | 62,91 | 34 | 8 | 8 | 0,983 | -0,025 | 15,59 | 16 |
| 58,30 | 5 | 4 | 4 | 1,140 | 0,189 | 53,79 | 5 | 3 | 3 | 0,868 | -0,204 | 43,45 | 6 |
| 6,93 | 2 | 2 | 2 | 0,969 | -0,046 | n. def. | 1 | 1 | 1 | 0,899 | -0,154 | 17,98 | 3 |
| 48,01 | 11 | 14 | 14 | 1,066 | 0,092 | 22,26 | 35 | 11 | 11 | 1,125 | 0,170 | 18,00 | 23 |
| n. def. | 0 | 0 | 0 | NA | 5,917 | n. def. | 0 | 0 | 0 | NA | 3,289 | n. def. | 0 |
| 2,55 | 2 | 1 | 1 | 0,882 | -0,181 | 5,60 | 3 | 1 | 1 | NA | -1,014 | n. def. | 0 |
| n. def. | 0 | 0 | 0 | NA | 1,769 | n. def. | 0 | 0 | 0 | NA | 3,728 | n. def. | 0 |
| n. def. | 0 | 1 | 1 | 1,383 | 0,468 | n. def. | 1 | 1 | 1 | 0,735 | -0,443 | n. def. | 1 |
| 22,15 | 10 | 8 | 8 | 0,939 | -0,090 | 53,58 | 30 | 9 | 9 | 0,955 | -0,066 | 15,70 | 17 |
| 187,47 | 10 | 12 | 12 | 2,285 | 1,192 | 52,23 | 21 | 7 | 7 | 2,143 | 1,100 | 132,49 | 13 |
| 67,77 | 9 | 4 | 4 | 1,011 | 0,016 | 16,28 | 10 | 5 | 5 | 1,195 | 0,258 | 46,56 | 8 |
| n. def. | 0 | 0 | 0 | NA | 0,983 | n. def. | 0 | 0 | 0 | NA | 2,501 | n. def. | 0 |
| n. def. | 0 | 1 | 1 | 1,290 | 0,367 | n. def. | 1 | 0 | 0 | NA | 0,763 | n. def. | 0 |
| 55,21 | 27 | 16 | 15 | 1,393 | 0,478 | 26,01 | 62 | 14 | 14 | 1,383 | 0,467 | 16,30 | 44 |
| n. def. | 1 | 2 | 2 | 13,765 | 3,783 | 8,16 | 3 | 1 | 1 | 12,190 | 3,608 | 40,27 | 2 |
| n. def. | 0 | 1 | 1 | 1,219 | 0,286 | n. def. | 1 | 1 | 1 | 1,016 | 0,023 | n. def. | 1 |
| n. def. | 0 | 0 | 0 | NA | 4,948 | n. def. | 0 | 0 | 0 | NA | 2,715 | n. def. | 0 |
| n. def. | 0 | 0 | 0 | NA | -0,799 | n. def. | 0 | 0 | 0 | NA | 4,163 | n. def. | 0 |
| 68,99 | 6 | 2 | 2 | 1,125 | 0,170 | 14,71 | 2 | 1 | 1 | 1,303 | 0,382 | n. def. | 1 |
| n. def. | 0 | 0 | 0 | NA | -0,758 | n. def. | 0 | 0 | 0 | NA | 2,555 | n. def. | 0 |
| n. def. | 0 | 0 | 0 | NA | 3,569 | n. def. | 0 | 0 | 0 | NA | 4,443 | n. def. | 0 |
| n. def. | 0 | 0 | 0 | NA | 1,209 | n. def. | 0 | 0 | 0 | NA | 4,803 | n. def. | 0 |
| n. def. | 1 | 2 | 2 | 1,361 | 0,445 | n. def. | 1 | 2 | 2 | 1,129 | 0,175 | n. def. | 1 |
| n. def. | 0 | 0 | 0 | NA | -0,192 | n. def. | 0 | 0 | 0 | NA | 1,506 | n. def. | 0 |
